# Dynamic merging of enhancer- and promoter-associated assemblies is coupled to transcriptional bursting

**DOI:** 10.64898/2026.10.26.761858

**Authors:** Qinhua Lin, Bo Wang, Yiwen Liu, Zuhui Wang, Junwei Zhang, Yiting Wang, Kairui Guo, Shuang Hui, Xuejie Zhang, Cheng Ni, Zhixing Chen, Wulan Deng

## Abstract

Classical models attribute distal enhancer function to chromatin looping, whereas emerging models propose that spatially organized assemblies of transcriptional regulators also facilitate enhancer-promoter communication. To investigate how these processes relate to transcription, we developed an endogenous-tagging-based four-color live-cell imaging platform and an integrated quantitative analysis pipeline in mouse embryonic stem cells (mESCs), enabling simultaneous visualization of regulatory assembly dynamics, chromatin organization and transcriptional bursting at the endogenous *Sox2* locus. We find that the pluripotency transcription factor OCT4 and the coactivator BRD4 form numerous spatially distinct but dynamically interacting assemblies. At non-bursting *Sox2* alleles, the promoter and the distal *Sox2* enhancer SCR occupy distinct assembly environments, with stronger BRD4 association at the SCR than at the promoter. At bursting *Sox2* alleles, transcription is coupled to transient merging and partial overlap of enhancer- and promoter-associated assemblies, with the SCR, promoter and nascent RNA frequently sharing OCT4/BRD4-enriched assembly environments across a broad range of SCR-promoter distances. Pharmacological perturbation of bromodomain-dependent BRD4 function preferentially suppresses long-duration, high-amplitude bursts. Together, our findings support a model in which transcriptional bursting is coupled to the dynamic reorganization and merging of enhancer- and promoter-associated regulatory assemblies, linking chromatin-looping and assembly-based views of enhancer-promoter communication.

## Main Text

Gene expression is controlled in part through communication between distal enhancers and promoters, a process classically attributed to chromatin looping^1–10^. Our previous work demonstrated that engineered enhancer-promoter looping is sufficient to activate silent genes^11,12^, establishing a causal role in gene activation. However, subsequent imaging studies have reported apparent enhancer-promoter distances extending to hundreds of nanometers at some active genes^13–15^. Although such measurements should be interpreted in light of potentially confounding technical factors such as labeling offsets and localization uncertainty, they raise the question of how regulatory communication occurs when the measured distance exceeds what would be expected for direct molecular contact. Thus, how distal enhancers communicate with promoters in living cells remains poorly understood.

In parallel, an emerging model proposes that transcription factors, coactivators and RNA polymerase II (Pol II) form spatially concentrated assemblies, commonly referred to as transcriptional condensates, through multivalent interactions, often involving intrinsically disordered regions (IDRs)^16–21^. These assemblies have been proposed to amplify transcription by concentrating Pol II and regulatory factors within specialized microenvironments^22–25^, and advanced fluorescence microscopy has detected such assemblies at or near active genes^20,23,25^. Because transcription occurs in bursts^26,27^, understanding how these assemblies contribute to enhancer action requires resolving not only where regulatory factors are located, but also how their dynamics are coordinated with burst initiation, duration and amplitude. Recent live-cell imaging studies have further linked assembly dynamics to transcriptional output, suggesting that transient interactions between regulatory assemblies and target genes may modulate bursting^25^. However, these studies have largely focused on Pol II and Mediator, whose enrichment at active genes is closely coupled to transcriptional output, making it difficult to distinguish upstream regulatory events from the transcriptional process itself. Consequently, it remains unclear whether upstream transcription factors and coactivators form a single mixed compartment or distinct, dynamically interacting assemblies, and how their organization contributes to enhancer-promoter communication and coordinates with transcriptional bursting in living cells.

Here, we used the endogenous *Sox2* locus in mESCs as a model system, as its transcription is strongly dependent on the distal Sox2 Control Region (SCR) enhancer cluster. We established mESC lines carrying up to five endogenous knock-ins to label OCT4, BRD4, the *Sox2* promoter, the SCR, and nascent *Sox2* RNA. We combined complementary super-resolution imaging modalities with an integrated quantitative analysis pipeline to selectively monitor up to four of these five components simultaneously across different labeling configurations. This approach enabled us to dissect the interplay between regulatory assemblies, cis-regulatory elements and transcriptional bursting. Using this system, we identify a dynamic regulatory transition in which enhancer- and promoter-associated assemblies transiently merge during transcriptional bursting. Our results support a model in which higher-order chromatin organization and regulatory assembly dynamics jointly contribute to transcriptional activation, thereby linking chromatin-looping and assembly-based views of enhancer-promoter communication.

## Results

### Endogenous OCT4 and BRD4 form distinct yet dynamically interacting assemblies

To investigate the link between regulatory assembly dynamics and gene regulation, we focused on the upstream regulators, transcription factor OCT4 and coactivator BRD4, which provide regulatory inputs to Pol II-mediated transcription^25^. To visualize these low-abundance proteins without perturbing their native behavior, we endogenously tagged OCT4 and BRD4 with self-labeling tags (HaloTag or SNAP-tag) and labeled them with the newly developed live-cell-compatible BD dyes^28^. These dyes offer exceptional brightness and high photostability, enabling detection of weak endogenous signals. Endogenous tagging was essential, as overexpression artifacts can substantially inflate apparent clustering and local enrichment.

We employed two complementary imaging modalities: sparse deconvolution structured illumination microscopy (Sparse-SIM; ∼60 nm lateral resolution)^29^ and spinning-disk confocal microscopy equipped with a Live-SR super-resolution module (∼110 nm lateral resolution)^30,31^. This dual approach was essential because, while Sparse-SIM provided superior spatial resolution, it exhibited relatively rapid photobleaching under these labeling conditions and was used primarily for high-resolution structural characterization and validation. Together with high-performance dyes, Live-SR imaging enabled multicolor imaging and time-lapse acquisition of low-abundance endogenous proteins, and served as the primary platform for most quantitative analyses and dynamic measurements.

To enable quantitative analysis of low endogenous signal intensities in multicolor live-cell images, we further developed **Condensate-quant** (Extended Data Fig. 1 and Methods), an integrated computational framework that combines image denoising, PSF-informed deconvolution, HMRF-based segmentation and watershed separation to identify and resolve individual regulatory assemblies. The framework extracts assembly boundaries, centroids, sizes and spatial overlap, enabling quantitative measurement of their relationships with DNA loci and nascent RNA. We performed extensive and systematic validation of assembly segmentation and boundary determination, as well as the localization accuracy of assembly centroids and DNA/RNA foci (Extended Data Fig. 1l and Supplementary Notes 1 and 2). Together with chromatic aberration correction across fluorescence channels along the x, y, z axes (Extended Data Fig. 2 and Methods), these validations establish the reliability of the assembly measurements and spatial distance analyses used throughout this study.

We began by investigating how endogenous OCT4 and BRD4 molecules organize in mESCs. Using CRISPR/Cas9-mediated genome editing, we introduced a HaloTag at the C-terminus of OCT4 or BRD4 in mESCs (Fig. 1a and Extended Data Fig. 3a, b). By labeling the endogenously tagged proteins with BD566_HTL_ and imaging them using Sparse-SIM, we observed that both OCT4-Halo and BRD4-Halo each formed numerous small, spatially separated regions of local enrichment in mESCs (Fig. 1b). Here, we use the term “assembly” to denote such a spatially confined region of local OCT4 or BRD4 enrichment, without making assumptions about its mechanism of formation or material state. These results were supported by Sparse-SIM imaging of fluorescently immunostained cells (Extended Data Fig. 3c), consistent with previous reports^17,18,32^. We further examined their distribution using Live-SR imaging in living cells (Extended Data Fig. 3d, e). Similar spatial patterns of assemblies were observed, with BRD4 exhibiting more pronounced local enrichment. Time-lapse Live-SR imaging further showed that OCT4 and BRD4 assemblies underwent dynamic spatial reorganization on timescales of tens of seconds, including apparent merging and separation events (Fig. 1c and Video S1). We use “merging” to describe the imaging-observed transition of initially spatially distinct OCT4- and BRD4-enriched assemblies into a shared or partially overlapping spatial configuration, without implying liquid-phase coalescence or any other specific physical mechanism. Consistent with prior observations of nanoscale clustering in cells^17,18,32^, these dynamics may reflect contributions from chromatin organization, protein-DNA and protein-protein interactions, and local concentration effects. To examine whether OCT4 and BRD4 form shared assemblies as proposed by the transcriptional-condensate model^16–21^, we stained endogenously tagged BRD4-Halo with BD566_HTL_, and labeled endogenous OCT4 with an AF488-conjugated antibody. Two-color 3D Sparse-SIM revealed that while OCT4 and BRD4 assemblies colocalized in certain locations, they exhibited largely distinct spatial distributions, with Manders’ coefficients of approximately 0.1 (Fig. 1d, e, Extended Data Fig. 3f and Video S2). Quantitative analysis revealed an average assembly radius of ∼100 nm and thousands of such assemblies per nucleus (Fig. 1e). To eliminate the effect of signal amplification with antibody staining and to examine their relationship in living cells, we generated double knock-in mESCs expressing endogenously tagged OCT4-Halo and BRD4-SNAP, and selected BD566_HTL_ and BD626_STL_ to optimize dual-color imaging quality (Fig. 1f). Imaging with either Sparse-SIM or Live-SR in living cells, we observed a consistent spatial organization in which OCT4 and BRD4 assemblies were only partially colocalized throughout the nucleus (Fig. 1g, h, Extended Data Fig. 3g). The spatial relationship between the two assembly types was examined in 3D, confirming that their apparent overlap was not caused by axial projection (Extended Data Fig. 3h). Condensate-quant, which incorporates denoising and deconvolution, further enhanced Live-SR image quality and facilitated quantitative analysis of assemblies (Fig. 1h, i). Time-lapse imaging of OCT4 and BRD4 assemblies revealed that their proximity and partial overlap were transient and reversible (Fig. 1j and Video S3). In fact, they did not remain as a single stable entity, but frequently separated again over time.

**Fig. 1.**
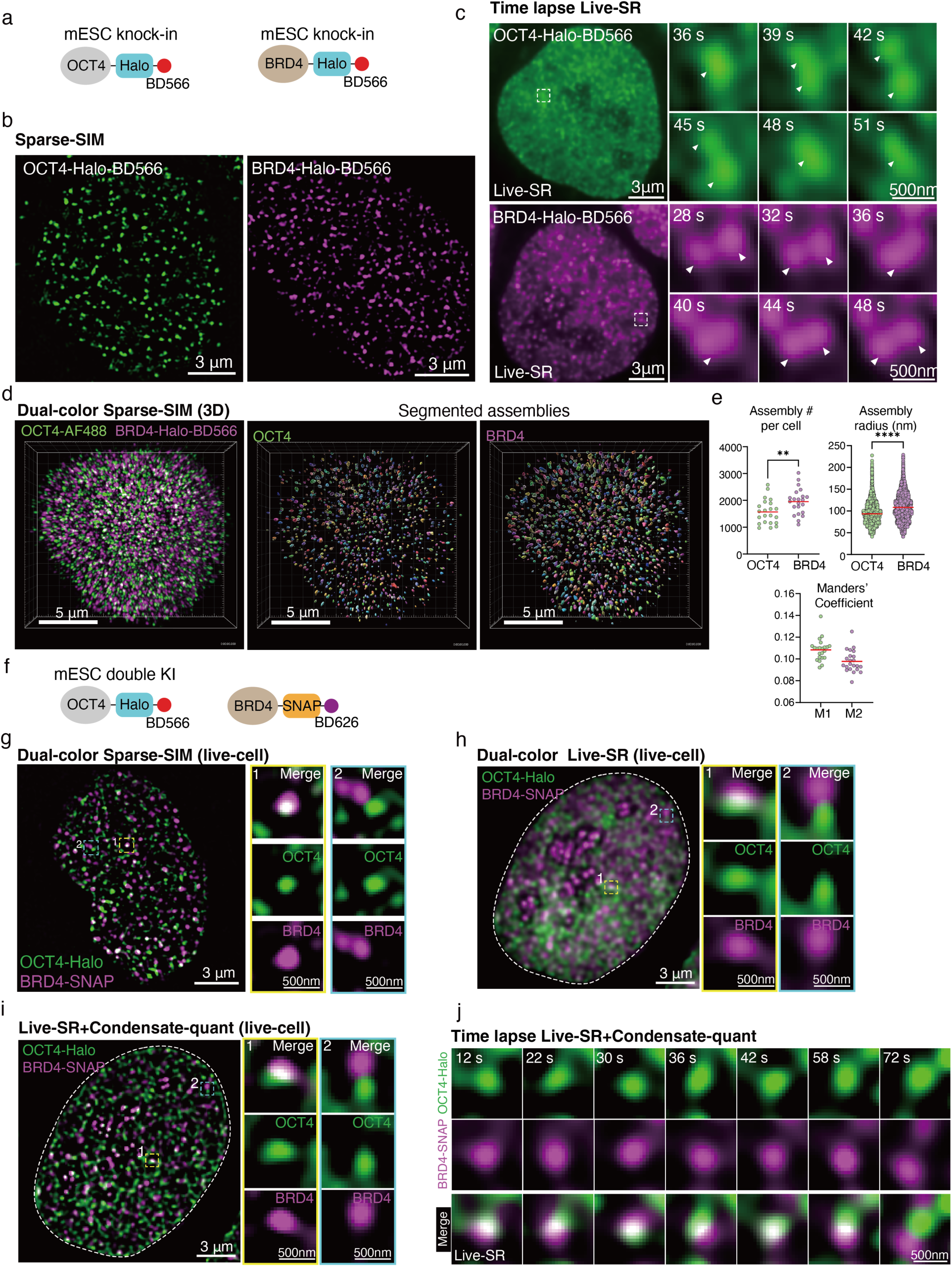
Spatial organization and dynamic interactions of endogenous OCT4 and BRD4 assemblies in mESCs. (**a**) Schematics of OCT4-Halo (heterozygous) and BRD4-Halo (homozygous) knock-in mESC lines and labeling with BD566_HTL_. (**b**) Sparse structured illumination microscopy (Sparse-SIM) images of fixed knock-in mESCs labeled with BD566. (**c**) Live-cell time-lapse super-resolution (Live-SR) images acquired at 1-s intervals; selected frames show assembly merging and separation events (white arrows). (**d**) Representative 3D Sparse-SIM image of a fixed BRD4-Halo mESC, with BRD4-Halo labeled with BD566_HTL_ and OCT4 immunostained with AF488; images were segmented in 3D. (**e**) Quantification of OCT4 and BRD4 assemblies from 3D Sparse-SIM, including the number of assemblies per nucleus, average assembly radius and colocalization (Manders’ coefficients M1 and M2). Data include 42,965 BRD4 assemblies and 34,534 OCT4 assemblies from 22 cells. (**f**) Schematic of double knock-in mESCs (OCT4-Halo, heterozygous; BRD4-SNAP, homozygous) labeled with BD566_HTL_ and BD626_STL_. (**g-i**) Dual-color live-cell Sparse-SIM, Live-SR and Condensate-quant-processed Live-SR images; zoomed ROIs show (1) merged and (2) separated assemblies. (**j**) Time-lapse Live-SR ROIs showing dynamic OCT4-BRD4 interactions (2-s intervals). All images are pseudocolored. Statistical significance: two-sided Student’s t-test; **P < 0.01, ****P < 0.0001; n.s., not significant.

These observations revealed that endogenous OCT4 and BRD4 organize into distinct yet dynamically interacting assemblies. This spatial organization differs from that observed in *in vitro* reconstitution systems, in which transcription factors and coactivators co-condense within a common droplet phase^18,22^. The partial overlap between OCT4- and BRD4-enriched assemblies in living cells suggests that the regulatory environment is spatially and compositionally heterogeneous rather than uniformly mixed. We speculate that their organization is at least in part shaped by the underlying DNA sequences, the distinct protein-interaction networks associated with OCT4 and BRD4, and other local chromatin features^33^. Nevertheless, this dynamic and heterogeneous behavior prompted us to investigate how transient interactions between assemblies relate to transcriptional regulation at endogenous loci.

### Sox2 bursting coincides with transient merging of OCT4 and BRD4 assemblies

Given the dynamic properties of regulatory assemblies, we investigated the relationship between assembly dynamics and transcriptional regulation using *Sox2* as a model locus. In mESCs, *Sox2* expression is controlled by the distal SCR enhancer cluster, located approximately 110 kb downstream of the promoter^34^ and displaying the characteristic features of a super-enhancer^35^. The SCR forms a chromatin loop with the *Sox2* promoter and contributes to over 90% of its transcriptional activity^34–36^. Thus, the *Sox2* locus serves as a suitable model for studying the spatiotemporal dynamics of regulatory assemblies in transcriptional activation. We engineered a triple knock-in mESC line (3KI: BRD4-SNAP, OCT4-Halo, *Sox2*-24×MS2) for simultaneous three-color live-cell tracking of endogenous assemblies and nascent RNA production. Specifically, building on our double knock-in line, we inserted 24 copies of MS2 repeats into the 3’UTR of both endogenous *Sox2* alleles and stably expressed the fluorescent MS2-binding protein stdMCP-EGFP (Fig. 2a). Single-cell clones with high signal-to-noise nascent RNA signals were selected to ensure robust detection (Fig. 2b). We examined the effects of genome editing on steady-state *Sox2* expression by RT-qPCR. OCT4-Halo/BRD4-SNAP double knock-in did not significantly affect *Sox2* expression (Extended Data Fig. 4a). Although the MS2 insertion reduced steady-state *Sox2* expression likely owing to reduced RNA stability^13^, robust nascent RNA foci and transcriptional bursting remained readily detectable, enabling analysis of assembly organization in relation to the *Sox2* transcriptional state (Video S4). Substantial heterogeneity of bursting states was observed between alleles within a cell, across cells, and across imaging fields (Extended Data Fig. 4b, c). Consistent with prior reports^25^, *Sox2* transcription in mESCs occurred in bursts. In randomly selected snapshots, detectable Sox2 nascent RNA foci were observed in 39% of cells and 25% of alleles (Fig. 2c).

**Fig. 2.**
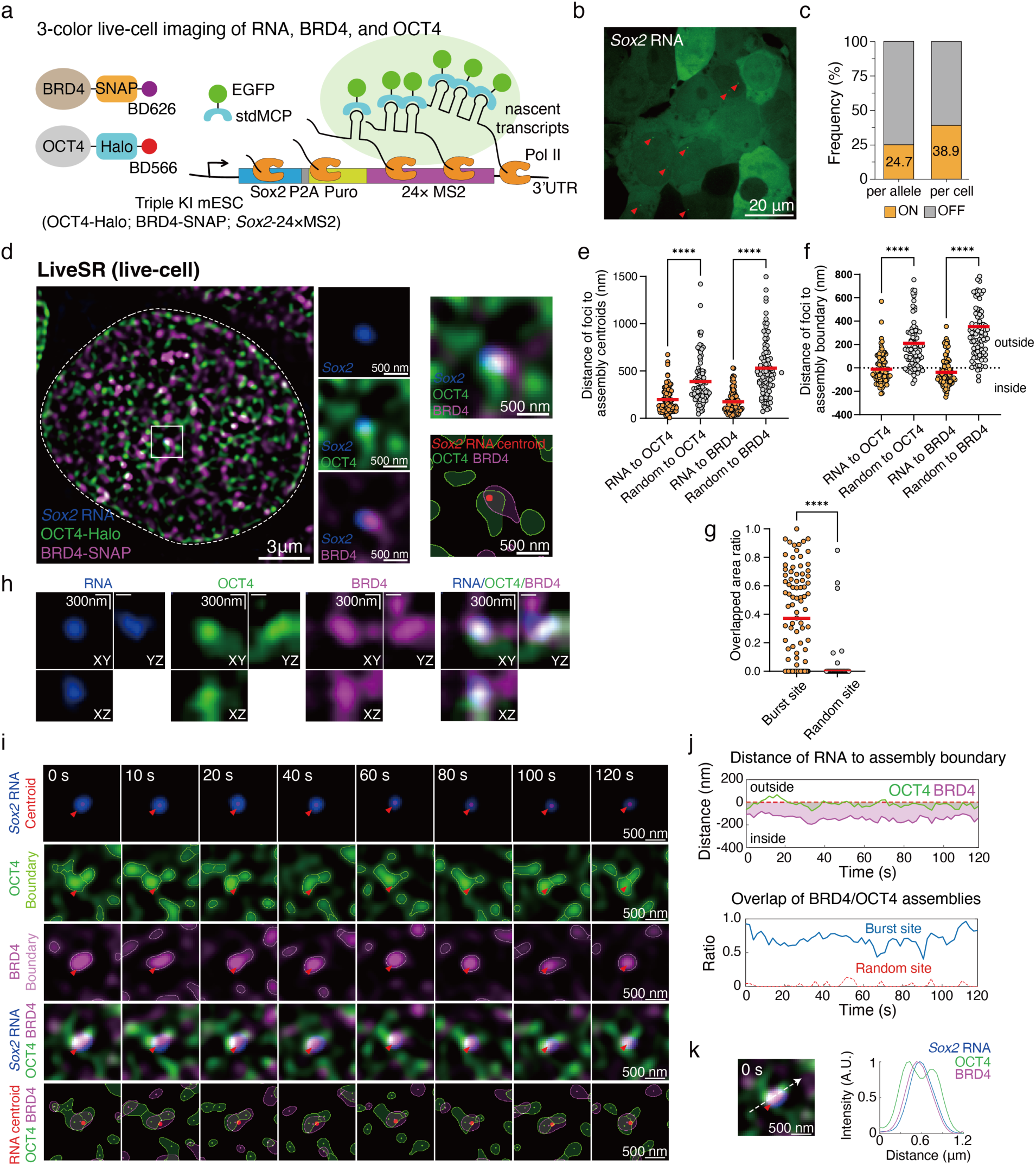
Spatial association and dynamic merging of OCT4 and BRD4 assemblies at bursting *Sox2* loci. **(a)** Schematic of three-color live-cell imaging; a triple knock-in clone with homozygous 24×MS2 integration was used. (**b**) Representative view showing *Sox2* RNA bursts (red arrows) in mESCs. (**c**) Quantification of the fractions of cells and alleles with detectable Sox2 nascent RNA foci in randomly selected snapshots (n = 807 cells). (**d**) Condensate-quant-processed live-cell Live-SR images showing RNA bursts and regulatory assemblies; selected ROIs illustrate the association between RNA bursts and merged OCT4-BRD4 assemblies. (**e–g**) Distances from RNA centroids or control random sites to the nearest OCT4 or BRD4 assembly centroids (**e**) and boundaries (**f**), together with the overlap ratios of the nearest OCT4 and BRD4 assemblies at bursting or random sites (**g**), calculated as the overlapping area divided by the BRD4 assembly area, were quantified from live-cell Live-SR images (n = 93 cells). (**h**) 3D intensity projection of RNA (blue), OCT4 (green) and BRD4 (magenta). (**i**) Time-lapse three-color confocal imaging (2-s intervals; three-slice z-stacks); *bottom*, resolved RNA centroids and assembly boundaries. (**j**) Time-course quantification of distances from RNA foci to assembly boundaries (top) and OCT4-BRD4 assembly overlap ratios (bottom, calculated as the overlap area divided by the BRD4 assembly area), corresponding to (**i**). (**k**) Snapshot and line intensity profiles across the bursting site and adjacent assemblies. Statistical significance: Mann-Whitney U test; ****P < 0.0001.

To visualize the spatial relationships between regulatory assemblies and the actively transcribing gene, we used the Live-SR module to acquire z-stack images of nascent RNA (GFP labeled), OCT4-Halo (BD566_HTL_ labeled) and BRD4-SNAP (BD626_STL_ labeled) assemblies in living cells (see Methods). Because *Sox2* is a short gene, its nascent RNA focus can serve as a spatial marker of the transcription site^13,25^. Notably, although OCT4 and BRD4 formed largely segregated assemblies across the nucleus, bursting *Sox2* loci were preferentially associated with overlapping OCT4- and BRD4-enriched assemblies (Fig. 2d). To quantify these spatial relationships across many cells (Fig. 2d-g), we used Condensate-quant to segment protein assemblies and identify the centroids of nascent RNA foci and assemblies, as well as assembly boundaries. Using complementary metrics, we measured the distance from each nascent RNA focus to either the centroid (Fig. 2e) or the boundary (Fig. 2f) of the nearest OCT4 or BRD4 assembly. Compared with randomly selected nuclear positions, bursting *Sox2* loci were significantly closer to both OCT4 and BRD4 assembly centroids and were preferentially located within or adjacent to both assembly types. Moreover, the OCT4 and BRD4 assemblies associated with bursting loci showed substantial but incomplete overlap (Fig. 2d, and g). Orthogonal inspection of 3D image stacks confirmed that this overlapping configuration was not caused by axial projection (Fig. 2h).

To determine whether this overlapping assembly configuration persisted during minutes-long bursts, we performed three-color time-lapse imaging. Although the positions of the RNA focus relative to the OCT4 and BRD4 assemblies fluctuated during individual bursts, the overlapping OCT4-BRD4 configuration was detected at an average of approximately 70% of RNA-positive time points across the analyzed cells (Fig. 2i-k, Extended Data Fig. 4d, and Video S5). We also acquired longer time-lapse recordings to capture the relatively infrequent transitions associated with burst onset and termination. In one illustrative recording, changes in OCT4-BRD4 assembly organization were observed during an interval surrounding the appearance of a *Sox2* RNA focus; in another, BRD4 dissociation was observed near the loss of RNA signal (Extended Data Fig. 5a-f; Videos S6-S7). However, the limited fluorescence signals from endogenously tagged OCT4 and BRD4 constrained our ability to capture enough of these infrequent transitions to establish a population-level temporal relationship between assembly reorganization and burst onset or termination. Together with the enrichment of simultaneous OCT4 and BRD4 association at RNA-positive loci, these observations support an association between transient OCT4-BRD4 assembly overlap and the *Sox2* bursting state.

We next asked whether bursting *Sox2* loci were also associated with a broader set of transcriptional regulators beyond OCT4 and BRD4, by combining MS2-based nascent RNA labeling with immunofluorescence staining of key transcription factors (SOX2 and NANOG), coactivators (MED1 and p300), and RNA polymerase II (RPB1). Through quantification of many cells, all examined factors formed puncta that exhibited stronger association with nascent RNA foci than with random nuclear positions (Extended Data Fig. 4e, f). Collectively, these analyses indicate that bursting Sox2 loci are associated with a multicomponent regulatory environment containing transcription factors, coactivators, and the core transcriptional machinery.

### Cis-regulatory elements associate with distinct regulatory assemblies at non-bursting alleles

We next investigated the spatial relationship between assemblies and *cis*-regulatory elements to determine how these assemblies engage with the *Sox2* locus architecture. We engineered two quadruple knock-in mESC lines (4KI: BRD4-SNAP, OCT4-Halo, *Sox2*-24×MS2, Prom/SCR-144×CuO) for simultaneous four-color live-cell imaging of *cis*-regulatory DNA elements, transcriptional activity, and endogenous OCT4 and BRD4 assemblies. Specifically, building on the 3KI cell, we inserted a *CuO* repeat array (144 repeats) at either a promoter-proximal position (8 kb upstream of the TSS) or an SCR-proximal position (4 kb downstream of the SCR) on one *Sox2* allele (Fig. 3a and Extended Data Fig. 6a; see Methods). Because these arrays are offset from their corresponding regulatory elements, they serve as promoter-proximal and SCR-proximal positional labels, hereafter referred to as the promoter and SCR labels. To visualize DNA and RNA, we stably co-expressed the CuO-binding protein CymR-3×mTagBFP2 and the MS2-binding protein stdMCP-EGFP, and selected single-cell clones with robust labeling suitable for imaging. The promoter and SCR foci were located in the vicinity of the nascent RNA foci, as expected from their positions within the *Sox2* locus (Extended Data Fig. 6b). Cells displaying closely duplicated DNA foci, indicative of separated sister chromatids, were excluded, leaving only loci with a single resolvable DNA focus for analysis.

**Fig. 3.**
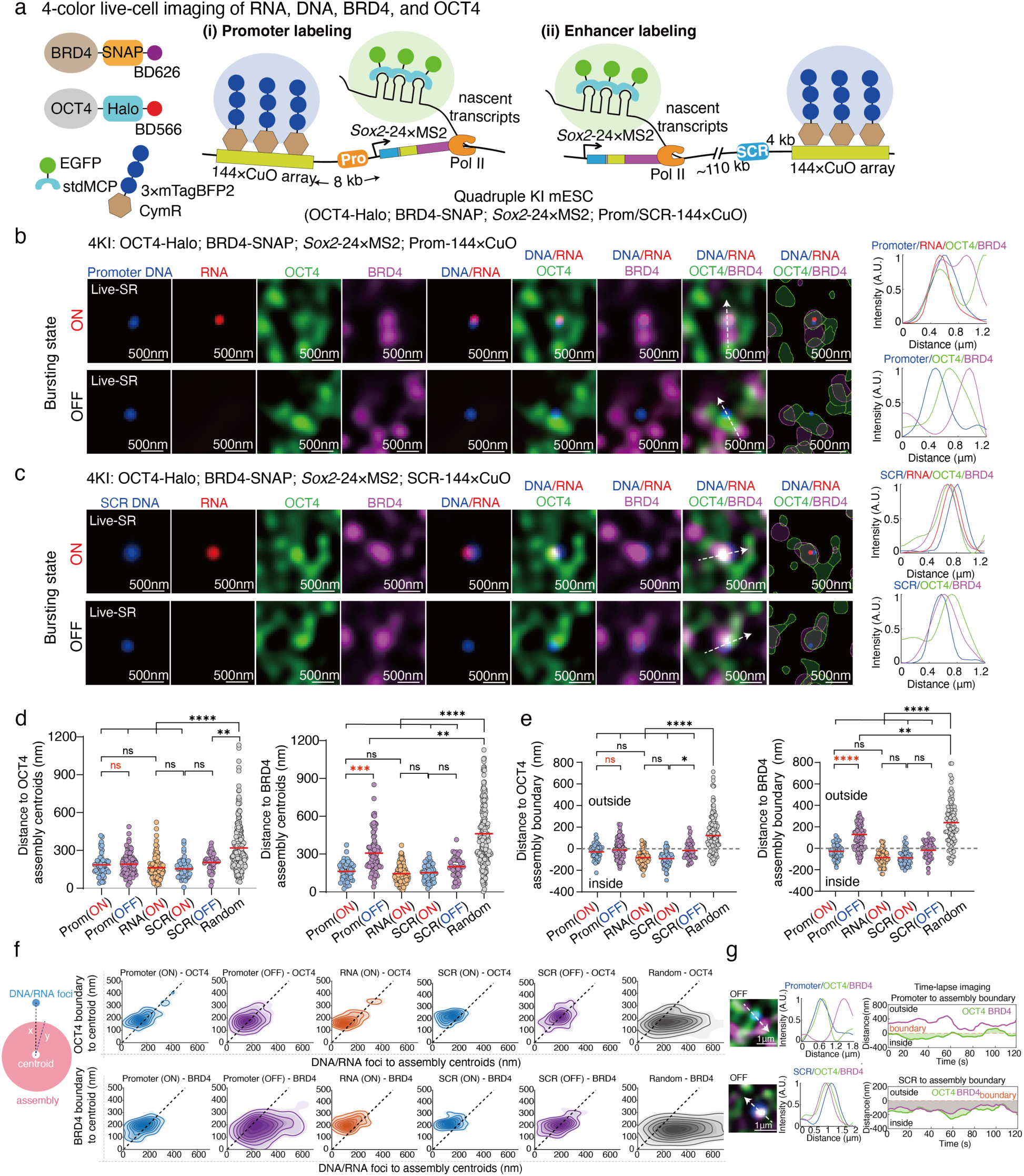
Distinct associations of regulatory assemblies with the Sox2 promoter and SCR. **(a)** Schematic of four-color live-cell imaging; two quadruple knock-in clones with heterozygous integration of a CuO array near the *Sox2* promoter or SCR were used. (**b**, **c**) Live-cell Live-SR images of DNA elements (blue; promoter in **b**, SCR in **c**), RNA (red), OCT4 (green) and BRD4 (magenta) assemblies at bursting (ON) or non-bursting (OFF) alleles (five-slice z-stacks). Resolved DNA/RNA centroids and assembly boundaries and line intensity profiles are shown on the right. (**d**) Distances from DNA/RNA centroids to the nearest assembly centroid (left, OCT4, right, BRD4) in promoter-labeled or SCR-labeled cells at ON or OFF alleles. Promoter: n = 48 (ON) and 92 (OFF). SCR: n = 57 (ON) and 44 (OFF). (**e**) Distances from DNA/RNA centroids to the nearest assembly boundaries (left, OCT4, right, BRD4) in promoter-labeled or SCR-labeled cells at ON or OFF alleles. (**f**) Contour plots of distances from DNA/RNA centroids to assembly centroids versus assembly radii, measured as the distances from assembly boundaries to their centroids, at ON or OFF alleles, corresponding to Extended Data Fig. 7c. (**g**) Time-lapse imaging of OFF alleles in promoter-labeled (top) or SCR-labeled (bottom) cells (2-s intervals). Each panel shows a representative snapshot (left), corresponding line intensity profiles (middle) across the labeled locus and adjacent assemblies, and time-lapse distance measurements (right). Related time-lapse image sequences are in Extended Data Fig. 8b, d. Statistical significance: Kruskal–Wallis test followed by Dunn’s multiple-comparisons test; *P < 0.05, **P < 0.01, ***P < 0.001, ****P < 0.0001; n.s., not significant.

This four-color system (DNA: BFP; RNA: GFP; OCT4: BD566_HTL_; BRD4: BD626_STL_) enabled us to simultaneously visualize OCT4- and BRD4-enriched assemblies, a labeled DNA element, and nascent RNA at single-locus resolution and, importantly, to compare their spatial organization between bursting and non-bursting states (Fig. 3 and Extended Data Fig. 7). At bursting *Sox2* loci, identified by bright stdMCP-EGFP foci, OCT4 and BRD4 assemblies showed a high degree of spatial overlap, with both the nascent RNA focus and the labeled SCR or promoter associated with a shared OCT4/BRD4-enriched assembly environment (Fig. 3b, c, top panels). This pattern was consistent with our observation in the 3KI cells (Fig. 2). At non-bursting *Sox2* loci, identified by promoter or SCR labeling in the absence of a detectable stdMCP-EGFP focus, promoter association with OCT4 appeared largely maintained relative to bursting loci, whereas its association with BRD4 was reduced. In contrast, SCR association with both OCT4 and BRD4 appeared largely maintained between the two transcriptional states (Fig. 3b, c, bottom panels).

We next assessed whether these transcription-state-dependent spatial patterns were consistent across the population using two complementary distance metrics: the distance from each DNA/RNA focus to the centroid or boundary of the nearest assembly (Fig. 3d, e). Compared with bursting loci, promoters at non-bursting loci were significantly farther from the nearest BRD4 assembly centroid and also showed significantly reduced association with the nearest BRD4 assembly in the boundary-based analysis. To assess these spatial relationships in relation to assembly size, we plotted the distance from each DNA/RNA focus to the nearest assembly centroid against the corresponding assembly radius (Fig. 3f and Extended Data Fig. 7c). These plots further supported reduced promoter-BRD4 association at non-bursting relative to bursting alleles, whereas promoter-OCT4 association remained largely unchanged between the two states. In contrast, SCR association with OCT4 or BRD4 showed modest differences between bursting and non-bursting alleles, although these differences were not consistently significant across the two distance metrics.

Representative time-lapse recordings further illustrated these state-dependent spatial relationships over time (Fig. 3g and Extended Data Fig. 8a-e; Videos S8-S12). At non-bursting alleles, the stronger BRD4 association with the SCR than with the promoter label was consistent with the higher BRD4 ChIP-seq signal at the SCR than at the promoter-proximal region (Extended Data Fig. 8f).

Together, these differential associations support distinct regulatory assembly environments for the promoter and SCR at non-bursting alleles. Together with the population-level enrichment of OCT4-BRD4 overlap at bursting loci (Fig. 2), these results support a model in which dynamic merging of enhancer- and promoter-associated regulatory assemblies is coupled to transcriptional bursting.

### SCR and promoter share a regulatory assembly environment despite variable distances at bursting loci

Having established that transcriptional bursting is coupled to the merging of enhancer- and promoter-associated assemblies, we next investigated how assembly organization relates to the classical enhancer-promoter looping model. To this end, simultaneous monitoring of the enhancer, promoter, regulatory assemblies and transcriptional activity is critical. Therefore, we generated a quintuple-knock-in cell line (5KI: BRD4-SNAP, OCT4-Halo, *Sox2*-24×MS2, Prom-CuO, SCR-TetO). To generate this line, we inserted a *CuO* array at the promoter-proximal position (8 kb upstream of the TSS) and a *TetO* array at an SCR-proximal position (4 kb downstream of the SCR) on the same allele. The arrays were visualized using the fluorescent DNA-binding proteins CymR and TetR, respectively (Fig. 4a and Extended Data Fig. 9a; see Methods). Importantly, steady-state *Sox2* mRNA abundance was similar between the 5KI and *Sox2*-24×MS2 3KI cells, suggesting that insertion of the promoter-proximal *CuO* and SCR-proximal *TetO* arrays alone did not cause detectable changes in *Sox2* expression (Extended Data Fig. 4a). By expressing distinct fluorescently tagged DNA- and RNA-binding proteins and labeling HaloTag and SNAP-tag with spectrally distinct fluorophores, we could selectively image any four of the five components to dissect their spatiotemporal relationships. To ensure consistent high-quality imaging of this multi-component system, we established cell lines expressing appropriate levels of the requisite fluorescent DNA- and RNA-binding proteins.

**Fig. 4.**
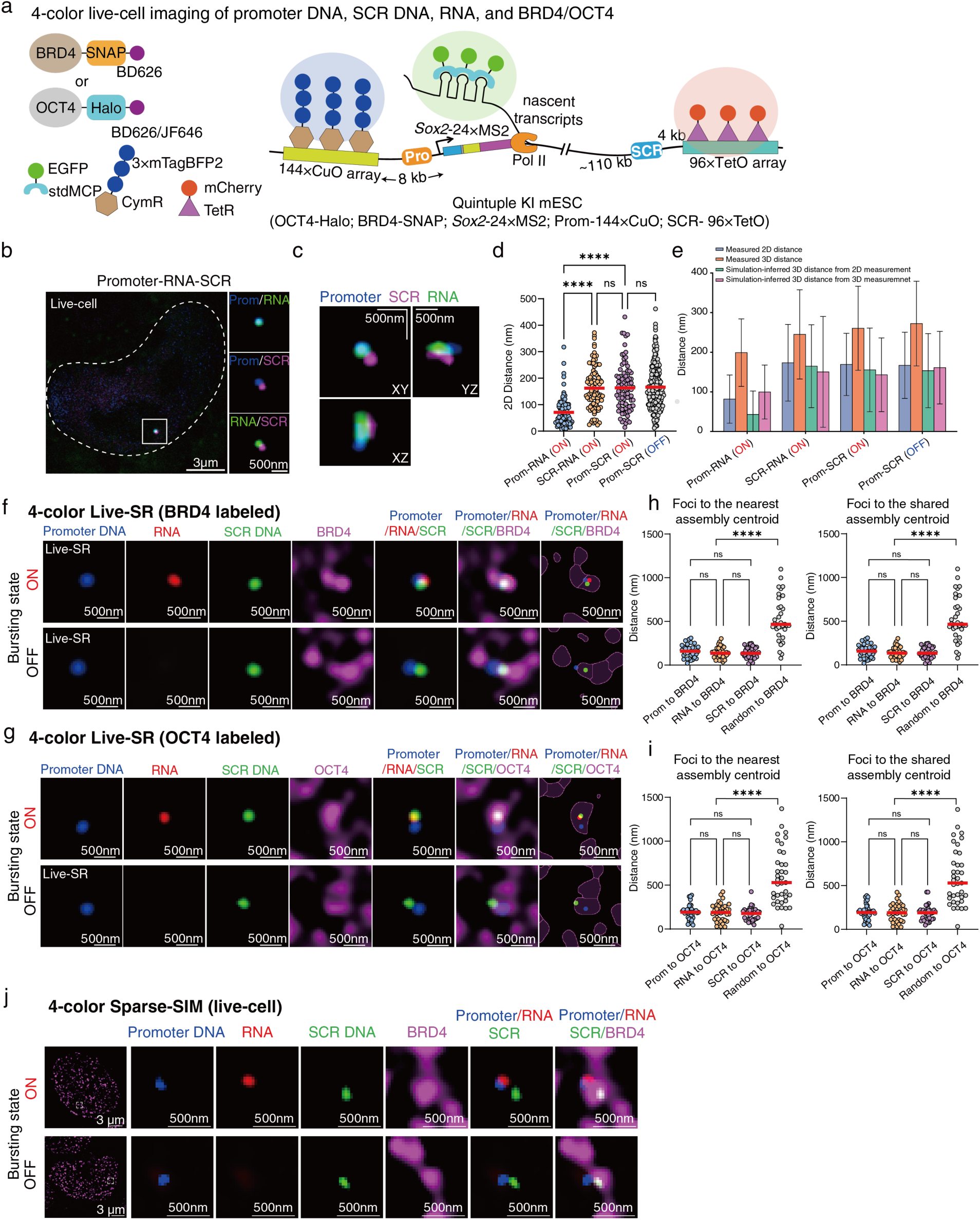
Variable SCR–promoter distance and shared assembly occupancy at bursting *Sox2* loci. **(a)** Schematic of flexible four-color live-cell imaging; a quintuple knock-in clone with CuO and TetO arrays integrated on the same *Sox2* allele (heterozygous) was used. (**b**) Live-cell Live-SR images of RNA, promoter and SCR; zoomed ROIs show their pairwise spatial relationships. (**c**) 3D intensity projections of Live-SR images. (**d**) 2D distances between the promoter and SCR, promoter and RNA, and SCR and RNA at ON alleles (n = 90), and between the promoter and SCR at OFF alleles (n = 221). (**e**) Comparison of measured distances and simulation-inferred underlying distances for DNA/RNA element pairs. Error bar represents standard deviation. (**f**, **g**) Live-cell Live-SR images of promoter (blue), RNA (red), SCR (green) and assemblies (magenta; BRD4 labeled with BD626_STL_ in **f**, OCT4 labeled with BD626_HTL_ or JF646_HTL_ in **g**) at ON and OFF alleles (five-slice z-stacks). (**h, i**) Distances from DNA/RNA centroids to the nearest corresponding individual assembly centroid (left) or to the centroid of the shared assembly (right); random sites served as controls. n = 37 (OCT4) and 34 (BRD4) cells. The shared assembly was operationally defined as the assembly closest to the geometric center of the three labeled loci. (**j**) Multicolor Sparse-SIM images of promoter, RNA, SCR, and BRD4 assemblies at ON and OFF alleles with ROIs. Statistical significance: one-way ANOVA; **P < 0.01, ****P < 0.0001; n.s., not significant.

We first examined the spatial relationships among the promoter, SCR, and nascent RNA using three-color 3D Live-SR imaging. Although the promoter (labeled by CymR-3×mTagBFP2) was consistently in close proximity to the nascent RNA focus (labeled by stdMCP-EGFP) at bursting loci, the SCR (labeled by TetR-mCherry) did not consistently colocalize with the promoter and was often spatially separated from it (Fig. 4b, c). We further quantified their pairwise distances across the population by localizing their centroids in 3D and measuring either 3D distances (Extended Data Fig. 9b) or distances after 2D projection (Fig. 4d) (see Methods). Both analyses showed that although all the SCR, promoter, and RNA foci were located in spatial proximity, within a few hundreds of nanometers of one another, the SCR-promoter and SCR-RNA distances were significantly greater than promoter-RNA distances. Moreover, at the population level, SCR-promoter distances were broadly distributed, with substantial overlap between the distributions at bursting and non-bursting loci, consistent with previous studies^13^. However, these population-level measurements do not exclude short-lived changes in SCR-promoter distance during individual burst events^37^.

We observed that the difference between SCR-RNA and promoter-RNA distances was less pronounced in the 3D measurements than in the 2D measurements. To investigate this discrepancy, we performed simulations incorporating measurement errors and found that, within the distance range and under the measurement-error conditions examined here, 2D projection-based distance estimates were more precise because the error introduced by projection was smaller than the additional axial localization uncertainty associated with direct 3D measurements (Supplementary Note 3). To account for genomic offsets between the inserted repeat arrays and their corresponding regulatory elements^38,39^, as well as localization uncertainty, we used chromatin-polymer simulations to infer the underlying distances between the functional elements (Supplementary Note 4). The inferred SCR-promoter distance showed a broad distribution centered at approximately 150 nm in both bursting and non-bursting alleles (Fig. 4e). It is worth noting that very short SCR-promoter distances, potentially compatible with direct contact as envisioned in classical looping models, were present in both states, indicating that such close configurations can occur but do not uniquely specify the bursting state. Together with previous imaging studies at *Sox2* and other loci^13,14,40^, these results support a model in which transcriptional bursting does not require sustained direct SCR-promoter contact.

We next examined which features of assembly organization were associated with transcriptional bursting. Specifically, we asked whether the spatial relationships of the two regulatory elements to OCT4- and BRD4-enriched assemblies differed with transcriptional state, using four-color live-cell imaging at single-allele resolution. At bursting *Sox2* loci, despite variable SCR-promoter distances, the SCR, promoter, and nascent RNA were simultaneously associated with the same regulatory assembly environment, whether defined by OCT4- or BRD4-enriched assemblies (Fig. 4f, g and Extended Data Fig. 9c, d). To assess this relationship across the population, we compared the distance from each SCR, promoter and RNA focus to its nearest assembly with its distance to the assembly closest to the geometric center of the three foci. The two measurements were highly similar, supporting a shared association of the three foci with the same OCT4- or BRD4-enriched assembly (Fig. 4h, i). This shared association was further supported by visual inspection of additional examples (Extended Data Fig. 10a, b). We also used an alternative four-color imaging scheme labeling RNA, OCT4, BRD4, and either the promoter or SCR DNA, and again observed a consistent pattern of shared association (Extended Data Fig. 10c, d). The nanoscale spatial organization of the *Sox2* locus and BRD4 assemblies was further resolved by 3D multicolor Sparse-SIM imaging (Fig. 4j). Together, these results indicate that bursting is associated with a shared regulatory assembly environment across a broad range of SCR-promoter distances. Thus, SCR-promoter distance alone does not uniquely distinguish the bursting state. These findings support a model in which dynamic reorganization and merging of regulatory assemblies are coupled to transcriptional bursting.

### Perturbation of BRD4 function preferentially suppresses long-duration, high-amplitude transcriptional bursts

The multicolor live-cell imaging results revealed a close association between assembly merging and transcriptional bursting. To perturb bromodomain-dependent BRD4 function, we treated cells with JQ1, a BET bromodomain inhibitor known to disrupt BRD4 interactions with acetylated chromatin and enhancer activity^41,42^. JQ1 rapidly dispersed most BRD4-enriched assemblies, and the BRD4 signal became largely diffuse throughout the nucleoplasm, although a small number of unusually large assemblies remained (Extended Data Fig. 11a). Live-cell imaging showed that JQ1 dispersed BRD4-enriched assemblies associated with the *Sox2* locus (Extended Data Fig. 11b), while ChIP-qPCR showed reduced BRD4 occupancy at the locus (Extended Data Fig. 11c). JQ1 treatment also significantly reduced *Sox2* RNA abundance, as measured by RT-qPCR (Extended Data Fig. 11d), and reduced detectable nascent RNA bursting by live-cell imaging (Extended Data Fig. 11e,f).

To further characterize the effect of JQ1 treatment on bursting kinetics at single-allele resolution, we performed long-term time-lapse imaging of nascent RNA production over one hour in hundreds of cells. We defined active alleles as those that exhibited at least one burst within the one-hour imaging window. We found that JQ1 treatment greatly reduced the fraction of time that active alleles spent in the bursting state (Fig. 5a, Extended Data Fig. 11e, and Video S13), while only mildly reducing the overall fraction of alleles classified as active (Extended Data Fig. 11g). To quantify the kinetic effects of JQ1 on transcriptional bursting, we applied a hidden Markov model (HMM) framework to define bursts and quantify burst parameters, including burst number, duration, amplitude and size, at single-allele resolution (Fig. 5b-d and Extended Data Fig. 11h). In untreated mESCs, *Sox2* alleles exhibited both short- and long-duration bursts, whereas in JQ1-treated cells, *Sox2* alleles exhibited predominantly short-duration bursts (Fig. 5e and Extended Data Fig. 11i). Notably, the long-duration, high-amplitude bursts that accounted for the majority of RNA production were preferentially lost upon treatment. Burst quantification showed that JQ1 significantly reduced burst amplitude, duration and size (Fig. 5f), but had only a minor effect on the number of bursts detected within the one-hour imaging time (Fig. 5g).

**Fig. 5.**
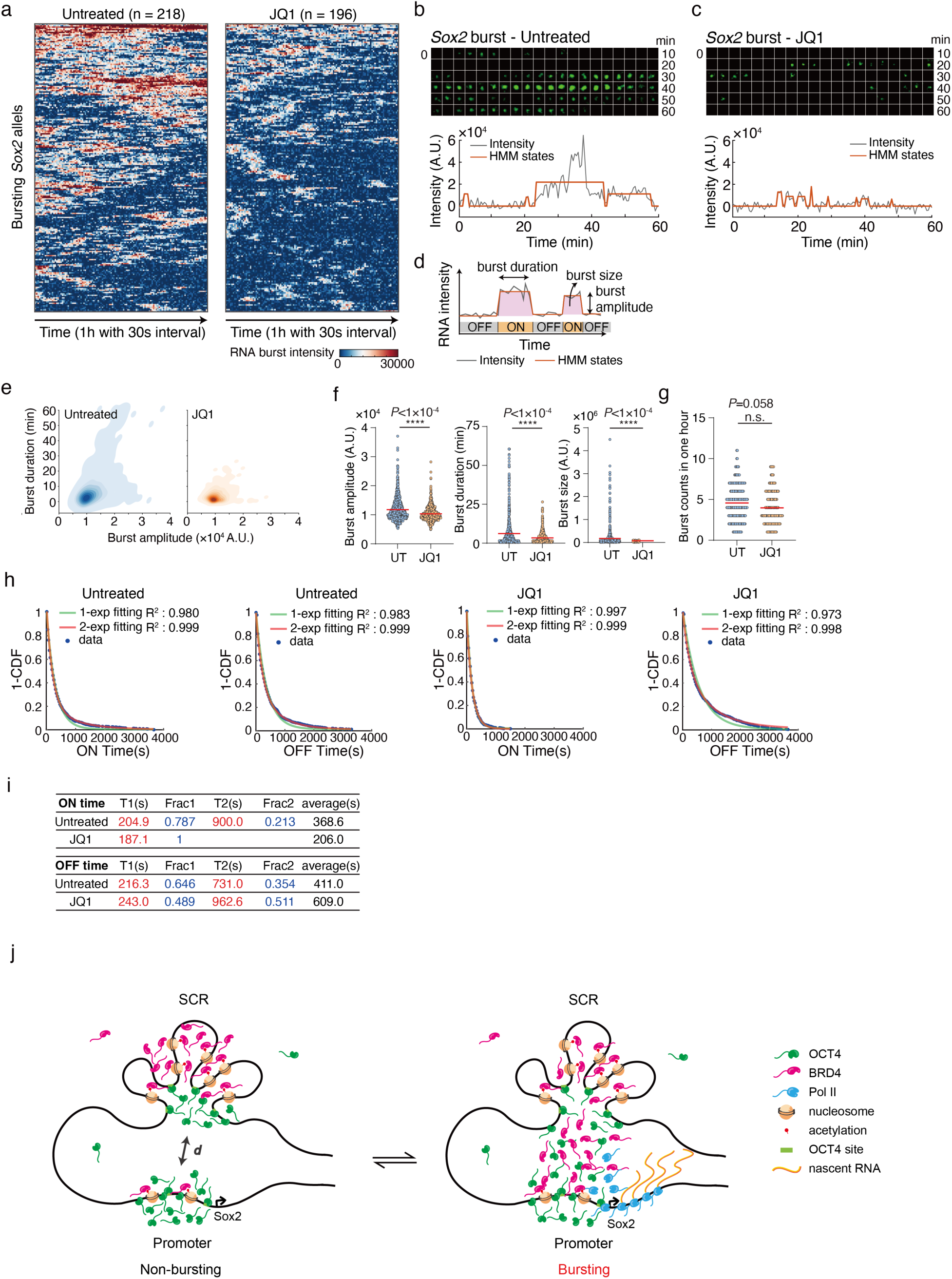
Effects of BRD4 perturbation on Sox2 transcriptional bursting kinetics. **(a)** RNA intensity heatmap of bursting alleles from untreated cells (n = 218) and JQ1-treated cells (1 μM, 30 min; n = 196), analyzed from one-hour time-lapse confocal imaging acquired at 30-s intervals using nine-slice z-stacks. (**b, c**) Representative time-lapse image sequences of *Sox2* bursting alleles and overlay of RNA intensity with bursting features analyzed by a Hidden Markov Model (HMM) from untreated (**b**) or JQ1-treated cells (**c**). (**d**) Temporal RNA intensity profiles of representative bursting alleles from untreated or JQ1-treated cells, overlaid with HMM-defined bursting states. (**e**) Kernel density plots of burst amplitude versus duration in untreated and JQ1-treated cells. (**f**) Characteristics of *Sox2* bursts, including amplitude, duration and size. Mean values are indicated by red lines. (**g**) Scatter plot of the number of bursts per allele during the one-hour recording window; the red line indicates the mean. (**h**) Complementary cumulative distribution functions (1-CDFs) of ON- and OFF-state durations, with single- or double-exponential fits. (**i**) Fitted kinetic parameters of bursting states from two-component fits, except for the ON-time distribution in JQ1-treated cells, which was fitted with a single component. (**j**) Model in which chromatin organization establishes spatial proximity of SCR and promoter, while dynamic assembly merging is coupled to long-duration transcriptional bursting. *d* denotes the variable distances between the SCR and promoter. Statistical significance: two-sided Student’s t-test; n.s., not significant.

To further resolve these distinct kinetic states quantitatively, we analyzed the cumulative distribution functions (CDFs) of burst durations (ON times) and inter-burst intervals (OFF times) (Fig. 5h, i and Extended Data Fig. 11j). In untreated cells, the ON-time distribution was better described by a double-exponential fit, consistent with two kinetic components: short bursts (ON1 state, average ∼247 s) and long bursts (ON2 state, average ∼1078 s). In JQ1-treated cells, the ON-time distribution was adequately described by a single-exponential fit (average ∼211 s), consistent with the loss of the detectable long-burst component. In contrast, analysis of OFF-time distributions showed only modest changes upon JQ1 treatment, with a slight shift toward longer inter-burst intervals but no clear emergence of additional kinetic components. Together, these kinetic analyses indicate that JQ1 preferentially suppresses the long-duration burst population, whereas short-duration bursts persist and OFF-state kinetics are only modestly affected. Because JQ1 perturbs bromodomain-mediated BRD4 chromatin engagement and, in our imaging experiments, also disperses BRD4-enriched assemblies at the *Sox2* locus, these results support an important role for bromodomain-dependent BRD4 function in sustaining long-duration, high-amplitude *Sox2* bursts.

## Discussion

The mechanistic basis of distal enhancer-promoter communication in living cells remains a central question in gene regulation. By developing a four-color endogenous-tagging live-cell imaging system and the quantitative Condensate-quant analysis framework, our study shows that dynamic merging of enhancer- and promoter-associated assemblies is coupled to transcriptional bursting at the endogenous *Sox2* locus. These findings are not readily explained by a simplified model in which transcription factors and coactivators invariably form a single, uniformly mixed transcriptional compartment. Rather, OCT4- and BRD4-enriched assemblies can remain spatially separated, associate differentially with cis-regulatory elements and undergo transient merging during transcriptional bursting. This spatial and compositional heterogeneity does not exclude contributions from multivalent interactions, intrinsically disordered regions or other condensation mechanisms to assembly formation and reorganization. At non-bursting *Sox2* alleles, the promoter and distal SCR enhancer occupy distinct assembly environments characterized by differential OCT4 and BRD4 enrichment, with BRD4 enriched at the SCR relative to the promoter. During transcriptional bursts, BRD4 association with the promoter increases, and the SCR, promoter and nascent RNA co-reside within a shared OCT4/BRD4-enriched assembly environment despite variable SCR-promoter distances. The bursting locus is also associated with a broader set of transcriptional regulators, including SOX2, NANOG, MED1, p300 and RNA polymerase II, supporting its association with a multicomponent regulatory environment extending beyond OCT4 and BRD4. Together, these observations support a model in which transcriptional bursting at *Sox2* is coupled to the dynamic merging and reorganization of enhancer- and promoter-associated assemblies, without requiring all active alleles to adopt a uniformly close SCR-promoter configuration (Fig. 5j).

We propose a two-layer model in which higher-order chromatin organization and regulatory assembly dynamics operate at distinct but complementary levels of regulation. At the chromatin level, a pre-existing spatial proximity between the SCR and promoter may provide a permissive configuration for regulatory communication. At the assembly level, dynamic interactions and merging of enhancer- and promoter-associated assemblies may create a transient, coactivator-enriched microenvironment that supports transcriptional bursting, potentially supporting long-duration bursts. Although SCR-promoter distance alone did not uniquely distinguish the bursting state, we speculate that baseline proximity may facilitate dynamic overlap between enhancer- and promoter-associated assemblies, possibly by constraining the relative motion of the SCR and promoter. The relatively short SCR-promoter distances observed in our study and in previous imaging studies^43^ are consistent with this possibility, although the proposed role of spatial confinement remains to be tested. Thus, baseline proximity may be permissive for enhancer-promoter communication, whereas dynamic assembly overlap is coupled to transcriptional bursting. This two-layer model integrates previous observations that *Sox2* transcription is enhancer-dependent but not tightly coupled to stable SCR-promoter proximity^13^ with studies linking transcriptional output to Pol II- or Mediator-associated assembly dynamics^25^.

This model raises several important mechanistic questions. BRD4-enriched assemblies at the SCR may act as local reservoirs of coactivator activity that become accessible to the promoter through transient assembly interactions, potentially supporting sustained Pol II engagement, transcriptional efficiency or long-duration bursting^20,44^. Consistent with a role for BRD4 in sustaining transcription, bromodomain inhibition with JQ1 preferentially suppresses long-duration, high-amplitude transcriptional bursts. However, because JQ1 affects both BRD4 chromatin engagement and BRD4-enriched assemblies, these experiments do not distinguish their respective contributions. The molecular determinants controlling the timing, frequency and duration of OCT4-BRD4 assembly interactions remain unclear. OCT4, BRD4 and other regulatory assembly components contain intrinsically disordered regions, whose multivalent interactions may contribute to assembly reorganization, merging, separation and heterotypic interactions in cells. Thus, enhancer potency may be encoded not only in DNA sequence and chromatin state, but also in the dynamic interaction properties of enhancer- and promoter-associated regulatory assemblies. Finally, although Sox2 is a powerful model for long-range enhancer-promoter regulation, extending this imaging approach to other loci will help establish the broader applicability of the proposed model and reveal potential mechanistic diversity in enhancer-promoter communication.

In summary, our live-cell imaging study reveals a spatiotemporal dimension of enhancer function in which regulatory assembly organization is coupled to transcriptional bursting across a broad range of enhancer-promoter distances. The resulting two-layer model connects chromatin spatial organization with regulatory assembly dynamics and provides a framework for testing how these features jointly shape transcriptional output.

## Acknowledgements

We thank Dr. Xiaohua Shen for sharing CJ9 mESCs, Dr. Yulong Li for sharing PhiC31 integrase plasmid, and Dr. Luke Lavis for sharing JF646_HTL_ dye. We thank the National Center for Protein Sciences at Peking University for providing the FACS service, and the High-Performance Computing Platform of the Center for Life Sciences for part of the data analysis.

## Funding

National Key Research and Development Program of China 2020YFA0509502 (WD)

National Natural Science Foundation of China 32170566 (WD)

Beijing Advanced Innovation Center for Genomics at Peking University (WD)

Peking-Tsinghua Center for Life Sciences (WD)

## Author contributions

Conceptualization: WD, QL, BW, YL

Methodology: WD, QL, BW, YL, JZ, ZC

Investigation: WD, QL, BW, YL, XZ, YW, KG, SH, CN

Visualization: BW, YL, ZW

Funding acquisition: WD

Project administration: WD

Supervision: WD

Writing-original draft: WD, QL, BW, YL

Writing-review & editing: WD, QL, BW, YL

## Data availability

The underlying raw and processed data will be made publicly available upon peer-reviewed publication.

## Code availability

The custom analysis code and trained models used in this study will be made publicly available upon peer-reviewed publication.

## Competing interests

Peking University has filed a patent application relating to the Condensate-quant analysis framework described in this study, with WD, BW, and YL listed as inventors. The remaining authors declare no competing interests.

## Supplementary Materials

Materials and Methods

Extended Data Fig. 1–11

Supplementary Notes 1-4

Tables S1 to S3

Video S1 to S13

