## Supplementary materials for "Dynamic merging of enhancer- and promoter-associated assemblies is coupled to transcriptional bursting"

**The supplementary materials file includes:**

Materials and Methods

Extended Data Fig. 1–11

Tables S1 to S3

Video S1 to S13

#### Materials and Methods

##### Cell culture

All cell lines used in this study were derived from mouse embryonic stem cells (CJ9, 46C)[45]. Cells were cultured on gelatin-coated plates in standard mESC medium, composed of DMEM (Gibco, C11995500BT) supplemented with 15% heat-inactivated fetal bovine serum (VisTech, SE100-011), 1% GlutaMAX (Gibco, 35050061), 1% penicillin/streptomycin (Corning, 30-002-CI), 1% EmbryoMax nucleosides (Millipore, ES-008-D), 0.1 mM 2-mercaptoethanol (Gibco, 21985023), 1% MEM non-essential amino acids (Corning, R25-0250CI), and 1000 U/ml recombinant leukemia inhibitory factor (Millipore, ESG1107). Cells were maintained at 37 °C with 5% $CO_2$ in a humidified incubator, and the culture medium was changed every 24 h.

##### Knock-in of HaloTag or SNAP-tag into OCT4 or BRD4

Single-guide RNAs (sgRNAs) targeting the genomic regions near the translation stop codons of the mouse *Pou5f1* and *Brd4* genes were designed using the CRISPR tool in Benchling (https://www.benchling.com). These sgRNAs were cloned into a vector containing expression cassettes for Cas9 and the Venus fluorescent protein, with sgRNA expression driven by the U6 promoter. Homology-directed repair (HDR) donor constructs were generated by flanking either the Halo-3×Flag or SNAP-HA sequences with two ~800 bp homology arms amplified from genomic DNA.

For each knock-in experiment, 1.25 µg of the donor plasmid and 1.25 µg of the sgRNA/Cas9 plasmid were co-transfected into $1\times10^6$ mESCs using Lipofectamine 3000 (Thermo Fisher, L3000015). 48 hours after transfection, Venus-positive cells were isolated by fluorescence-activated cell sorting (FACS), plated at low density (5,000 cells per 10 cm dish), and cultured for clonal expansion. Individual clones were manually isolated. A fraction of cells from each clone were lysed using a direct mouse genotyping kit (APE×BIO), and screened by PCR to identify positive knock-in events. Candidate clones were then expanded, and genomic DNA was extracted using the TIANamp Genomic DNA Kit (TIANGEN). The genotype of each clone was validated by Sanger sequencing, and protein expression was confirmed by western blot analysis. Finally, clones exhibiting proper mESC morphology and expected protein expression levels were selected for subsequent experiments.

Using this approach, we first generated single knock-in mESC lines expressing either OCT4-Halo-3×Flag (heterozygous clone) or BRD4-Halo-3×Flag (homozygous clone). Subsequently, using the OCT4-Halo-3×Flag heterozygous clone as the parental line, we generated a double knock-in mESC line expressing OCT4-Halo-3×Flag (heterozygous) and BRD4-SNAP-HA (homozygous). All primer sequences are listed in Supplementary Table 1.

##### Knock-in of the 24×MS2 cassette into the *Sox2* gene for RNA labeling

For knock-in, an sgRNA targeting the 3’UTR of the *Sox2* gene was designed and cloned into the Cas9 expression vector described above. An HDR donor construct was generated by flanking a P2A-PuroR-24×MS2 cassette with two ~800 bp homology arms amplified from genomic DNA. In this construct, the 24×MS2 cassette comprises synonymous MS2 RNA motifs (MBSV5)[46], and PuroR encodes the puromycin-resistance gene used for positive-clone selection. For knock-in, 1.25 µg of the donor plasmid and 1.25 µg of the sgRNA/Cas9 plasmid were co-transfected into $1\times10^6$ cells from the double knock-in mESC line (OCT4-Halo; BRD4-SNAP) using Lipofectamine 3000 (Thermo Fisher). 48 hours after transfection, Venus-positive cells were isolated by FACS, plated at low density, and cultured for about 10 days under selection with 1 µg/mL puromycin (Selleck, S7417). Positive clones were identified by PCR screening following the procedures described above. We used a homozygous clone with the MS2 cassette inserted into both *Sox2* alleles for subsequent experiments.

To enable live-cell visualization of MS2-tagged transcripts, a piggyBac transposon system (System Biosciences) was subsequently used to generate stable cell lines expressing EGFP fused to the synonymous tandem MS2 coat protein (stdMCP-EGFP)[46] under the control of the moderate-strength L30 promoter[38]. Cells with low expression levels were isolated by FACS to minimize background signal. Single-cell clones exhibiting an optimal signal-to-noise ratio for RNA imaging were selected for all subsequent experiments. All primer sequences are listed in Supplementary Table 1.

**Insertion of the CuO repeat array into the *Sox2* promoter or SCR for DNA labeling**

A two-step strategy for array insertion was adapted from a previous publication[13] (Extended Data Fig. 6a). All subsequent genome editing was performed using the triple knock-in mESC line (OCT4-Halo; BRD4-SNAP, *Sox2*-24×MS2) as the starting material. First, Bxb1 *attP* and PhiC31 *attP* sites were knocked into the genomic loci near the promoter (~8 kb upstream) and SCR region (~4 kb downstream), respectively, via CRISPR/Cas9-mediated HDR. Homozygous clones carrying the *attP* insertions were obtained and validated by PCR screening and Sanger sequencing.

Subsequently, a 144×CuO repeat array (Addgene #119903) was integrated into the pre-installed *attP* sites using the corresponding *attB* donor constructs and integrases. For this purpose, two donor constructs were generated, each containing the 144×CuO array, a *loxP*-flanked PGK promoter-driven EGFP expression cassette, and either a Bxb1 or PhiC31 *attB* site. For integration, cells were co-transfected with 1.25 μg of the corresponding *attB* donor plasmid and 1.25 μg of either the Bxb1 integrase plasmid (Addgene #119901) or the PhiC31 integrase plasmid (a gift from Dr. Yulong Li's lab). EGFP-positive cells were isolated by FACS and plated for clonal isolation.

Candidate clones were first screened by PCR, and further validated by imaging. The CuO-binding fusion protein CymR-3×mTagBFP2 was transiently expressed in each clone to identify the clones exhibiting bright, focal BFP signals. Selected clones were further validated by transient co-expression of stdMCP-mCherry (to label *Sox2* nascent RNA) and CymR-3×mTagBFP2. Correct genomic integration was confirmed by observing the expected proximity between mCherry-labeled RNA foci and mTagBFP2-labeled DNA loci. Only clones with heterozygous integration were obtained. The EGFP cassette was subsequently excised from validated clones via transient Cre recombinase expression, and EGFP-negative cells were isolated by FACS. This process yielded two quadruple knock-in mESC lines (OCT4-Halo; BRD4-SNAP; *Sox2*-24×MS2; Promoter-144×CuO or SCR-144×CuO).

For stable visualization, we established cell lines expressing the CymR-3×mTagBFP2 fusion protein. Cells were subjected to FACS and single-cell cloning to isolate clones with optimal signal-to-noise ratios for live-cell imaging, following a strategy analogous to that used for RNA imaging. All primer sequences are listed in Supplementary Table 1.

**Insertion of the TetO repeat array for dual DNA labeling at the *Sox2* locus**

We employed the same two-step strategy to introduce the TetO repeat array into the SCR in the established quadruple knock-in line (OCT4-Halo; BRD4-SNAP; *Sox2*-24×MS2; Promoter-144×CuO) (Extended Data Fig. 9a). Specifically, a PhiC31 *attP* site was first knocked into the genomic locus near the SCR via CRISPR/Cas9-mediated HDR, and homozygous clones were obtained. A donor construct was generated containing the 96×TetO repeat array (Addgene #174884), a *loxP*-flanked PGK promoter-driven EGFP expression cassette, and a PhiC31 *attB* site. Using the same procedure described above, EGFP-positive cells were isolated by FACS and plated for clonal isolation. Candidate clones were first screened by PCR, and further validated by imaging after transient co-expression of CymR-3×mTagBFP2 (to label the *Sox2* promoter) and TetR-mCherry (to label the SCR). Clones with correct genomic integration of the promoter and SCR labels on the same allele were selected by observing the expected proximity between the TagBFP2- and mCherry-labeled loci. Clones carrying the TetO insertion on one Sox2 allele were obtained. The EGFP cassette was subsequently excised from validated clones via transient Cre recombinase expression, and EGFP-negative cells were isolated by FACS.

For stable visualization, we established stable cell lines expressing the requisite fluorescent fusion proteins. Cells were subjected to FACS and single-cell cloning to isolate clones with optimal signal-to-noise ratios for live-cell imaging. All primer sequences are listed in Supplementary Table 1.

**Fluorescent dye labeling for live-cell imaging**

Eight-well chambered coverglasses (Cellvis, C8-1.5H-N) were coated with Matrigel overnight. Cells were seeded in the coated chambers at ~70% confluence in standard culture medium one day before imaging. For single-color imaging with HaloTag, cells were stained with 200 nM $BD566_{HTL}$ for 30 min in the cell incubator, and gently rinsed with pre-warmed PBS once to remove excess dye. After dye labeling, cells were incubated in pre-warmed fresh medium for at least 30 min, and the medium was changed to pre-warmed phenol red-free medium immediately before live-cell imaging. For multicolor imaging with both HaloTag and SNAP-tag, cells were first stained with 200 nM $BD626_{STL}$ for 2 hours in the cell incubator, and gently rinsed with pre-warmed PBS once to remove excess dye. Subsequently, cells were stained with 200 nM $BD566_{HTL}$ for 30 min, and processed as above. For co-imaging of HaloTag and three types of DNA/RNA foci, cells were stained with 200 nM $BD566_{HTL}$, $BD626_{HTL}$, or $JF646_{HTL}$ for 30 min.

**Immunofluorescence staining**

mESCs were cultured on Matrigel (Corning, 356231)-coated coverslips and pre-treated according to experimental requirements. After two washes with PBS, samples were fixed with pre-warmed 4% paraformaldehyde for 10 min at 37 °C, and then permeabilized with 0.1% Triton X-100 for 10 min at room temperature. Cells were subsequently blocked for 1 hour at room temperature with blocking buffer (3% BSA/0.1% Triton X-100/PBS). Primary antibodies, diluted in blocking buffer, were incubated with the samples overnight at 4 °C, followed by incubation with species-matched secondary antibodies for 2 hours at room temperature. After optional DAPI staining and final PBS washes, coverslips were mounted in glycerol and imaged by confocal microscopy. All primary and secondary antibodies are listed in Supplementary Table 3.

**Confocal and Live-SR imaging**

All confocal imaging was performed on a custom-built Nikon Ti2-E microscope equipped with a Yokogawa CSU-X1 spinning-disk confocal scanner, a Live-SR super-resolution module (GATACA Systems), a Nikon CFI Plan Apo VC 100× oil-immersion objective (NA 1.40), an EM-CCD camera (Andor iXon Life 897, 160 nm pixel size), and a stage-top incubator (Tokai Hit) maintained at 37 °C and 5% $CO_2$. Excitation was provided by OBIS laser modules at 405 nm (maximum 110 mW), 488 nm (maximum 165 mW), 561 nm (maximum 110 mW), and 642 nm (maximum 110 mW), coupled with a quad-band dichroic mirror (405/488/561/633 nm). Emission signals were filtered using bandpass filters specific to each fluorophore: 447/60 nm for BFP, 525/50 nm for EGFP, 617/73 nm for BD566, and 685/40 nm for BD626. The Perfect Focus System (PFS) was engaged throughout all time-lapse acquisitions to maintain stable focus. All hardware components were synchronized and controlled using NIS-Elements AR software (version 5.42.03) for automated and coordinated image acquisition.

For single-time-point imaging, the Live-SR module was employed to acquire images at a resolution beyond that of conventional confocal microscopy. To capture three-dimensional spatial information, z-stacks of 5 or 9 steps (with step sizes of 0.2 or 0.3 μm) were collected. Protein channels were imaged with an exposure time of 500 ms and four-frame averaging, whereas DNA and RNA were imaged with an exposure time of 200 ms. Multicolor 3D images were acquired sequentially in a plane-by-plane manner: at each z-position, all fluorescence channels were acquired before the microscope moved to the next z-plane. No beam splitters or simultaneous multi-camera acquisition were used. Super-resolution reconstruction was performed automatically by the Live-SR system through optical demodulation of the disk-stripe pattern, resulting in a final pixel size of 94.5 nm.

For the four-color imaging experiment of the promoter, BRD4, SCR, and RNA, the acquisition order at each z-plane was promoter, BRD4, SCR, and RNA. The promoter, SCR, and RNA channels were each acquired with a 200 ms exposure, whereas BRD4 was acquired as four 500 ms frames that were averaged to improve the signal-to-noise ratio. For averaged acquisitions, the midpoint of the integration window was used as the effective acquisition time. Based on this calculation, the effective midpoint time difference between BRD4 and the DNA/RNA channels was approximately 1.1-1.3 s.

For time-lapse imaging, the Live-SR module was used where indicated. For single- or two-channel imaging of protein assemblies, the Live-SR module was used with an exposure time of 300 ms for HaloTag fusion proteins and 500 ms for SNAP-tag fusion proteins. For three- or four-channel imaging of protein assemblies and DNA/RNA, the conventional spinning-disk confocal module was used with an exposure time of 500 ms for BRD4, 300 ms for OCT4, and 200 ms for DNA/RNA. Time-lapse movies were acquired either as single planes at 1- or 2-second intervals, or as three-step z-stacks (0.3 μm step size) at 3-second intervals. Movie durations ranged from 2 to 10 min. For long-term RNA burst imaging, nine-step z-stacks (0.5 μm step size) were acquired every 30 seconds for 60 minutes, with an exposure time of 100 ms per frame.

**Condensate-quant: a deep learning-assisted framework for imaging and analyzing weak assembly signals**

Live-cell imaging of endogenous biomolecular assemblies is challenged by intrinsically low signal intensity, even with bright fluorescent labels, as well as by imaging noise and the diffraction limit imposed by the point-spread function (PSF). To reliably extract quantitative information from such weak confocal time-lapse data, we developed **Condensate-quant** (Extended Data Fig. 1), a computational framework that sequentially applies a deep-learning-based denoising model and Lucy-Richardson (LR) deconvolution to enhance image quality, followed by a model-based intensity-classification step to segment assemblies. This framework enables robust quantification of assembly properties and their spatial relationships with other nuclear structures.

*Self-supervised image denoising with Noise2Void (N2V)*

To suppress noise and provide a cleaner input for subsequent deconvolution, which is highly sensitive to noise, we applied a deep-learning-based denoising approach. After comparing several algorithms, we selected the self-supervised Noise2Void (N2V) model[47] (Extended Data Fig. 1b) based on its performance on our data. The model was implemented using a U-Net architecture (Extended Data Fig. 1c). During training, image stacks were divided into 128 × 128 patches with a 32-pixel overlap to reduce boundary artifacts. Optimization was performed with the Adam optimizer (learning rate $5 \times 10^{-5}$; $\beta_1 = 0.5$, $\beta_2 = 0.99$) using a composite loss defined as the arithmetic mean of the L1 and L2 norms. The model typically converged within 10 epochs.

Performance was evaluated by comparing denoised images with high-SNR reference images generated from five-frame averages. Denoised outputs showed higher similarity to the reference than the raw images, as measured by the structural similarity index (SSIM) and Pearson correlation coefficient (Extended Data Fig. 1d, e).

*PSF-guided Lucy-Richardson deconvolution*

Insufficient deconvolution iterations limit resolution recovery, whereas excessive iterations often amplify noise and introduce artifacts. To quantitatively determine the appropriate number of iterations, we employed residual autocorrelation (RAC) analysis. The point-spread function (PSF) of the imaging system was empirically measured using 3D images of 100 nm TetraSpeck™ microspheres (Invitrogen) and fitted as described previously (Extended Data Fig. 1f)[48]. At each iteration, the deconvolved image was reconvolved with the measured PSF and subtracted from the original to generate a residual image. The RAC value initially decreased as image resolution improved and subsequently increased as excessive iteration amplified noise. The iteration yielding the minimum RAC was selected as the optimal stopping point. We found that prior denoising significantly improved deconvolution stability, allowing more iterations (approximately 20 in our implementation) without generating noise-related artifacts (Extended Data Fig. 1g, h).

*HMRF-based assembly segmentation*

Endogenous regulatory assemblies are small relative to the imaging pixel size. To facilitate smooth boundary delineation at the subpixel scale, we adopted a strategy combining up-sampling and intensity-based classification (Extended Data Fig. 1j). We first resized the images by a factor of 16 for confocal data (reducing the pixel size from 160 nm to 10 nm) or by a factor of 10 for Live-SR data (from 95 nm to 9.5 nm). Assembly regions were then segmented using a hidden Markov random field (HMRF) model, which classifies each pixel into one of seven intensity-based levels[49,50]. Pixels assigned to the selected highest-intensity classes (in this study, classes 5 to 7) were designated as assemblies. Adjacent assemblies within the same segmented region were further separated by marker-controlled watershed segmentation using LoG-detected assembly centroids, as described in the following section (Extended Data Fig. 1k). We systematically evaluated the performance of this method using simulated and experimental data, and compared it with alternative thresholding methods (Extended Data Fig. 1l and Supplementary Note 1).

*LoG-based assembly centroid localization*

Within each HMRF-defined mask, assembly centroids were detected using two-dimensional Laplacian-of-Gaussian (LoG) filtering followed by quadratic fitting for subpixel localization (Extended Data Fig. 1i). Following N2V denoising and LR deconvolution, images were processed using LoG filtering to enhance punctate structures while suppressing background signals. Candidate positions were identified as local maxima in the LoG response map, and centroid coordinates were refined by two-dimensional quadratic fitting of the local intensity distribution. We quantitatively assessed LoG localization accuracy using synthetic datasets with known ground truth (Supplementary Note 2).

When an HMRF-defined mask contained multiple adjacent assemblies, the LoG-detected centroids were used as markers for watershed segmentation, thereby separating the merged region into individual assemblies (Extended Data Fig. 1k). The resulting masks defined the final assembly boundaries, whereas the quadratically refined LoG coordinates served as assembly centroids for subsequent spatial analyses.

#### Chromatic aberration correction via multi-channel registration

Chromatic aberration was calibrated using fluorescent bead samples. Beads were prepared by depositing 0.1 μm TetraSpeck microspheres (Thermo Fisher, #T7284) on #1.5H precision coverslips (Thorlabs, #CG15XH1), except for calibration involving the 405 nm channel, for which 0.5 μm microspheres were used to improve bead signal intensity. Samples were mounted in ProLong Glass Antifade Mountant (Thermo Fisher, #P36982). Z-stack images of the beads were acquired in four fluorescence channels. Microspheres that were spatially well separated and exhibited high-quality localization fits were selected for analysis. Their positions were localized in 3D using the TrackMate plugin in Fiji with a Laplacian-of-Gaussian (LoG) detector (with the scale set to the estimated bead diameter), followed by quadratic fitting to achieve subpixel precision.

After obtaining corresponding 3D coordinate sets across channels, a 3D affine transformation was computed using the fitgeotform3d function in MATLAB to register all channels to a common reference frame. Following registration, residual inter-channel alignment errors had standard deviations of 5-10 nm laterally and 13-27 nm axially, depending on the channel pairs (Extended Data Fig. 2).

#### Quantification of assembly and foci properties and their spatial relationships

*Localization and distance measurement*

To quantify the spatial relationships between transcriptional assemblies and DNA/RNA foci, all experimental image stacks were first aligned using the 3D chromatic aberration correction transformation, and subsequently processed by denoising and deconvolution as described above. DNA and RNA foci were localized in 3D from the denoised, non-deconvolved images using a 3D LoG detector followed by 3D quadratic fitting to achieve subpixel localization precision, enabling 3D distance measurements between DNA and RNA foci (Supplementary Note 2).

To analyze assembly-foci spatial relationships, we adopted a hybrid 2D-3D analysis strategy. Because accurate 3D reconstruction of assembly boundaries is highly sensitive to axial PSF distortion and noise, assembly segmentation was performed on a restricted 2D projection rather than the full z-stack. Specifically, when two DNA/RNA foci were analyzed, the z-slices spanning their localized axial positions were averaged into a single projection plane for assembly analysis; when a single focus was analyzed, the z-plane containing that focus was used. Assembly boundaries were delineated using the HMRF-based segmentation method described above, and assembly centroids were subsequently identified using a 2D LoG detector. DNA and RNA focus coordinates were then projected onto the same 2D plane for distance calculations.

Two complementary spatial metrics were calculated: (1) the distance from each DNA or RNA focus to the nearest assembly centroid, and (2) the signed distance from each focus to the nearest assembly boundary, where positive values indicate positions outside the assembly boundary and negative values indicate positions inside the assembly. Because centroid localization is substantially less sensitive to PSF broadening than boundary determination, centroid-based measurements provide a more robust metric for spatial proximity analysis.

*Assembly enrichment level measurement*

The fluorescence enrichment level of each assembly was calculated as the ratio of the peak fluorescence intensity within the segmented assembly to the local background intensity. The local background was calculated as the mean intensity of pixels located two to three PSF standard deviations outside the segmented assembly boundary. This region was selected to minimize contributions from PSF-broadened assembly fluorescence while providing a representative measurement of the surrounding fluorescence intensity.

*RNA intensity measurement*

In time-lapse experiments analyzing the dynamic association between assemblies and transcription sites, RNA intensity was quantified as the integrated intensity within a 7 × 7-pixel window centered on the focus centroid, after subtracting the local mean background intensity. Photobleaching was corrected based on the temporal change in background intensity over the imaging series. For frames in which no distinct RNA focus was detected, the gene locus position was estimated by linear interpolation using the nearest preceding and subsequent detected foci.

**RNA burst kinetics analysis**

To quantify *Sox2* transcriptional bursting kinetics, long-term time-lapse images were acquired from the *Sox2*-24×MS2 cell line. More than 200 cells per condition were imaged using nine-step z-stacks (0.5 µm step size) every 30 seconds for 60 minutes, with an exposure time of 100 ms per frame. Cells exhibiting significant drift or phototoxicity were excluded from analysis. For image stacks from each cell, a maximum-intensity-z-projection was generated, from which the bursting transcription sites were then segmented at all time points using a random forest-based pixel classifier in Ilastik[51]. ROIs containing positions of each allele across all frames were manually created. In each ROI, the positions of bursting sites were identified as the pixels of maximum intensity within the Ilastik-generated masks, while positions during non-bursting states were estimated by linear interpolation. The burst intensity was defined as the integrated intensity of a 2D Gaussian fitted to the local signal. This yielded intensity trajectories of individual bursting alleles, as (t, x, y, I). All trajectories were visually inspected and manually curated before further analysis.

The intensity time series for each trajectory was subsequently analyzed with a hidden Markov model (HMM) implemented in custom MATLAB scripts. The OFF and ON transcriptional states were modeled as normal distributions. Model parameters (state distributions and transition probabilities) were optimized using the Baum-Welch algorithm, and the most likely sequence of states was inferred using the Viterbi algorithm. From the decoded state sequences, the mean burst amplitude, burst duration, and burst size were calculated.

Durations of ON and OFF states from multiple alleles were pooled to generate cumulative distribution functions (CDFs). Survival curves (1 − CDF) were fitted with either a single-exponential function

$$\mathrm{P}(t) = \mathrm{A}e^{-k_1 t}$$

or two-exponential function

$$\mathrm{P}(t) = \mathrm{A}e^{-k_1 t} + (1 - A)e^{-k_2 t}$$

where $1/k_1$ and $1/k_2$ represent the characteristic durations of the short and long lifetime components, respectively.

**Sparse-SIM imaging**

All structured-illumination microscopy (SIM) experiments were performed on a commercial Hessian-SIM system (His-SIM) equipped with a 100×/1.5 NA oil-immersion objective (Olympus) and excitation lasers at 405, 488, 561, and 638 nm. Image acquisition was controlled using IMAGER software. For each 3D-SIM dataset, z-stacks were acquired with a step size of 100 nm, using 5 phase shifts and 3 illumination angles (15 raw images per z-plane), yielding raw image stacks with a pixel size of 65 nm. SIM reconstructions were performed in IMAGER using the Sparse-SIM algorithm (Sparse level 1), yielding reconstructed images with a final pixel size of 32.5 nm.

Fluorescent bead samples were prepared using 0.1 μm and 0.5 μm TetraSpeck microspheres (Thermo Fisher, #T7279) deposited on #1.5H precision coverslips (Thorlabs, #CG15XH1) and mounted in ProLong Glass Antifade Mountant (Thermo Fisher, #P36982). For dual-color imaging (with excitation wavelengths of 488 and 561 nm), chromatic aberration was corrected by multichannel alignment using immobilized 0.1 μm TetraSpeck microspheres acquired under identical imaging conditions. For four-channel imaging, immobilized 0.5 μm TetraSpeck microspheres were used to improve the signal from 405-nm excitation.

In both cases, bead positions were localized in 3D using TrackMate (Fiji/ImageJ), and an affine transformation matrix was estimated to register the 405-, 488-, and 638-nm channels to the 561-nm channel. This transformation was applied to all experimental image stacks. After registration, the optimal z-range was manually selected to exclude reconstruction artifacts in low-SNR regions. To compensate for axial intensity decay during scanning, each z-slice was normalized by its mean background intensity.

**Sparse-SIM analysis**

Sparse-SIM images were analyzed for 3D cluster segmentation using custom scripts written in MATLAB. Briefly, intensities in each z-slice were rescaled to the range [0, 1] using the slice-specific minimum and maximum values. The resulting 3D stacks were then globally thresholded using Otsu's method (MATLAB function *graythresh* and *imbinarize*) to generate an initial binary mask. This mask was refined by morphological filtering, including removal of small connected components (MATLAB function *bwareaopen*) and 3D spherical closing (MATLAB function *imclose*). Candidate cluster centers were detected using a custom 3D Laplacian-of-Gaussian (LoG) filter, and only detections within the refined mask were retained. Marker-controlled watershed segmentation was then performed using the negative LoG response as the topographic surface and the detected centers as imposed minima, yielding individual 3D clusters. Clusters with volumes below the resolution limit were excluded from further analysis. For each retained cluster, a maximum-intensity projection along the z-axis was generated, and the equivalent radius in the xy plane was computed from the projected area.

For dual-color colocalization analysis, reconstructed Sparse-SIM image stacks were imported into Imaris (v10.1.0) and analyzed using the Colocalization module. Using this module, thresholded Manders' colocalization coefficients were computed after applying a fixed intensity threshold of 7 to both channels. SIM movie renderings were also performed in Imaris.

**RT-qPCR, ChIP-qPCR and ChIP-seq analysis**

Total RNA was extracted from $1\times10^6$ cells using the FastPure RNA Isolation Kit (Vazyme, RC113), and 1000 ng of RNA was reverse transcribed with the HiScript II cDNA Synthesis Kit (Vazyme, R212). ChIP-qPCR was performed according to standard protocols[52] in three experimental replicates; antibodies are listed in Supplementary Table S3. Quantitative PCR was carried out using ChamQ Universal SYBR qPCR Master Mix (Vazyme, Q711) on a Quantagene q225 real-time PCR system. All qPCR primers are provided in Supplementary Table 2.

Publicly available ChIP-seq datasets from mESCs were obtained from the GEO database, including GSE90893 (OCT4, SOX2, NANOG), GSE112806 (MED1, RPB1, BRD4), and GSE89211 (p300, H3K27ac, H3K4me1, H3K4me3). Raw reads were processed using a standard ChIP-seq analysis pipeline[50]. To compare factor enrichment at the promoter versus the SCR, we calculated the $\log_2$ enrichment score as the $\log_2$ ratio of the average read density in the SCR region (chr3:34,754,000–34,764,000; mm10) to that in the *Sox2* promoter-proximal region (chr3:34,645,000–34,655,000; mm10).

### Extended Data Figure 1

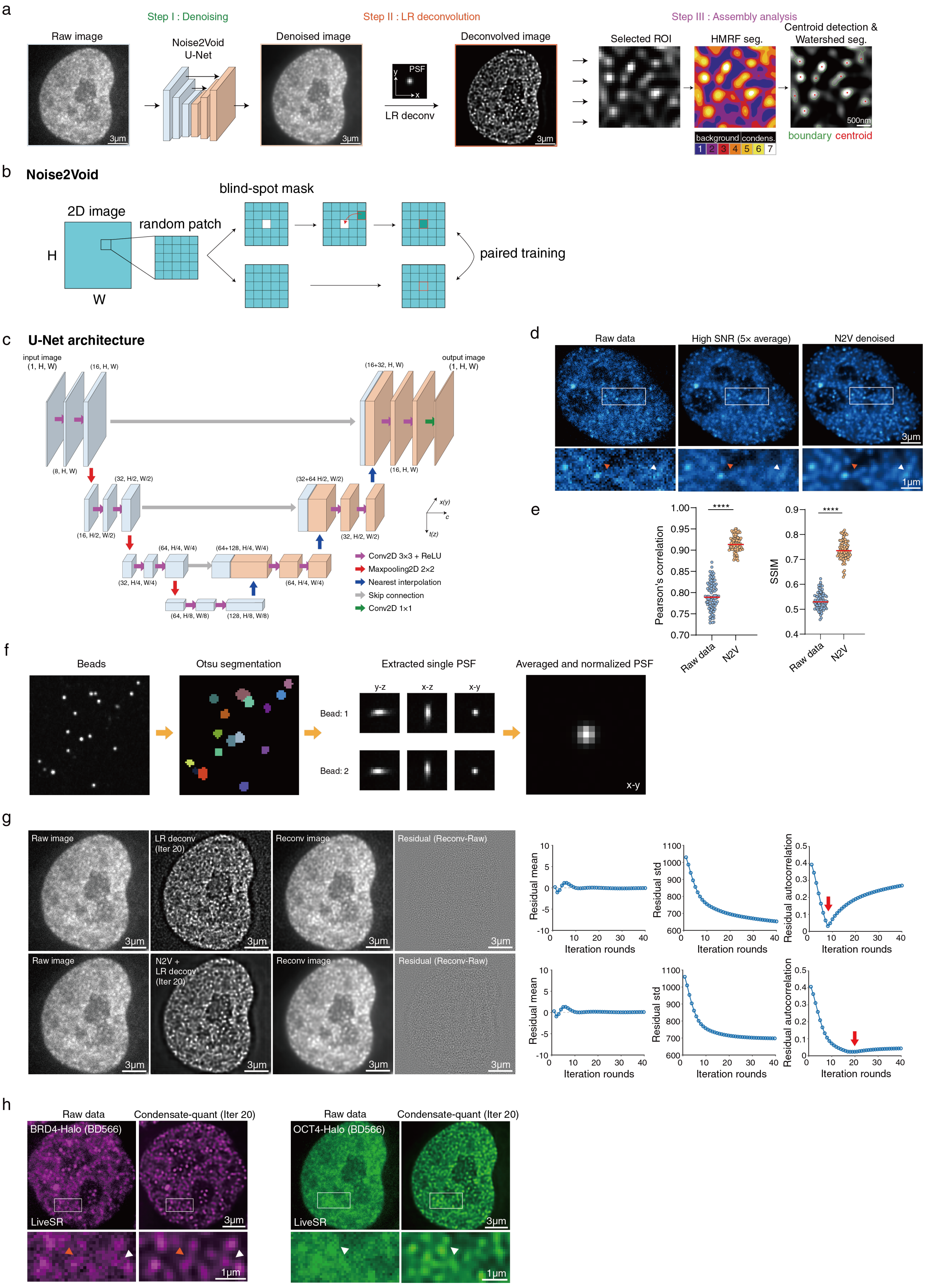


1

Extended Data Figure 1

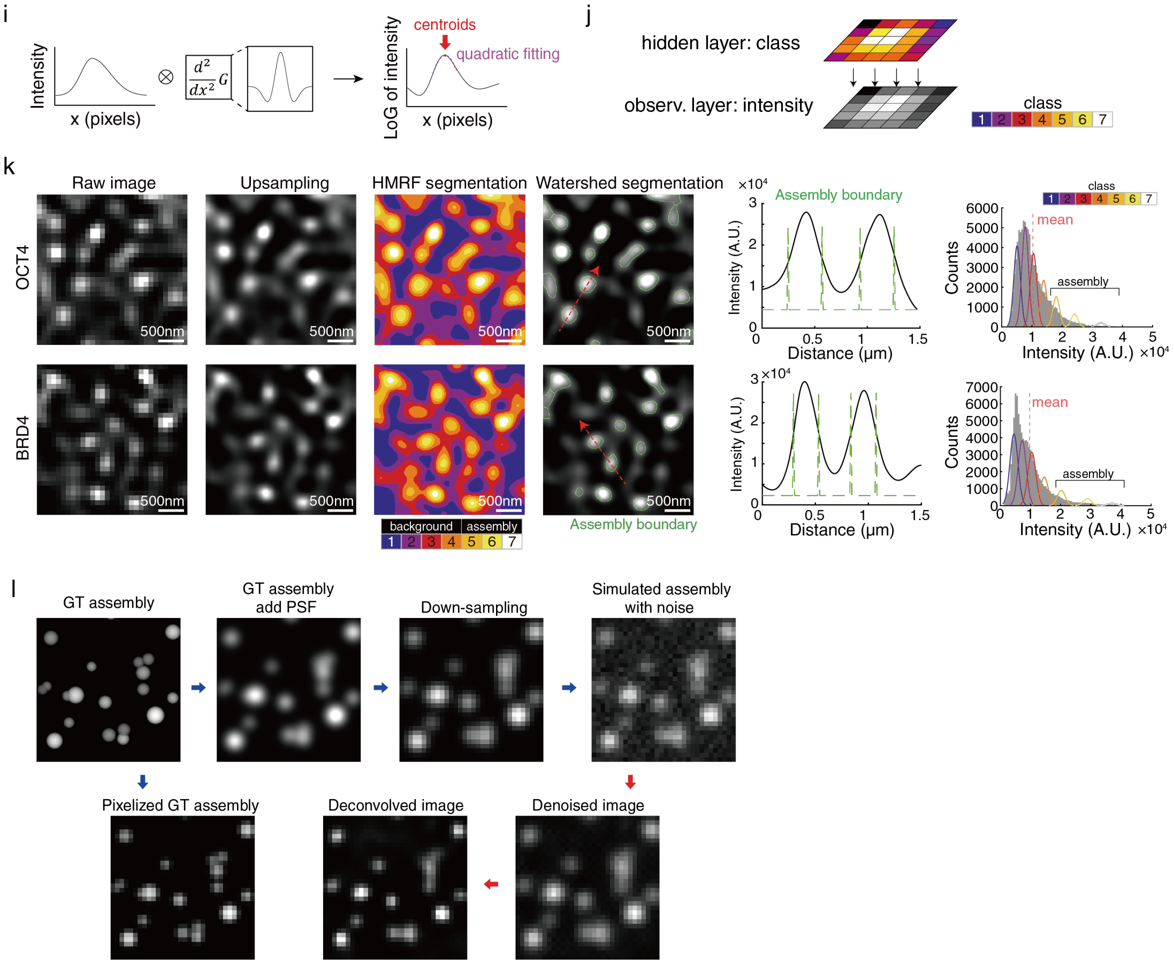


**Extended Data Fig. 1. Condensate-quant: an integrated pipeline for self-supervised denoising, deconvolution, and segmentation of transcriptional assemblies.** (**a**) Schematic overview of the Condensate-quant pipeline. Raw image sequences are first denoised using the N2V deep neural network, followed by Lucy-Richardson deconvolution using the experimentally measured PSF to obtain high signal-to-noise, super-resolved images. Individual assemblies are then segmented using the HMRF algorithm combined with watershed segmentation. **(b)** Schematic of the self-supervised Noise2Void (N2V) strategy. A random masking scheme replaces selected pixels with values sampled from their local neighborhood, allowing the network to learn denoising directly from noisy images without clean ground-truth data. (**c**) Architecture of the U-Net model. The network follows an encoder-decoder design with skip connections. The encoder extracts features via convolutional and down-sampling layers; the decoder restores resolution through up-sampling and skip-connected features. All convolutions use 3 × 3 kernels with ReLU activation. **(d)** Validation of the trained N2V network using time-lapse Live-SR images of endogenous BRD4-Halo. Single-frame raw images were denoised and compared against high-SNR reference images generated by averaging five consecutive frames. Insets (white boxes) show regions where denoising sharpens assembly boundaries (red arrows) and resolves adjacent assemblies (white arrows). (**e**) Scatter plots of Pearson correlation coefficients and structural similarity index (SSIM) between raw or N2V-denoised images and the corresponding high-SNR reference images (n = 90). Statistical significance: Wilcoxon paired signed-rank test; ****,

$p < 0.0001$. (**f**) Workflow for extracting the point-spread function (PSF) from fluorescent bead images. Large-field-of-view images of beads were acquired, and individual beads were segmented by Otsu thresholding to create masks. The PSF from each bead was then extracted, projected along the *x*-, *y*-, and *z*-axes, averaged across multiple beads, and normalized to generate the final PSF estimate. (**g**) Left: Comparison of Lucy-Richardson (LR) deconvolution applied to raw (top) and N2V-denoised (bottom) data over 20 iterations. Each panel shows the deconvolved image, the corresponding re-convolved result, and the residual map. Right: Plots illustrate the mean pixel value, standard deviation, and autocorrelation of residual images as functions of iteration number. Red arrow denotes the iteration at which RAC is minimized. **(h)** Live-cell images of mESCs with endogenous BRD4-Halo and OCT4-Halo labeled with BD566, shown before and after denoising and deconvolution (Condensate-quant pipeline). Insets (white boxes) display magnified regions where deconvolution resolves adjacent protein assemblies (white arrows). **(i)** Schematic of the Laplacian-of-Gaussian (LoG)-based localization algorithm. The input intensity profile is convolved with the second derivative of a Gaussian kernel to generate a LoG response, from which candidate peaks are identified. Sub-pixel centroid positions are then determined by quadratic fitting around the local maxima. **(j)** Schematic of the hidden Markov random field (HMRF)-based segmentation algorithm. The observation-layer *intensity* was modeled as a Gaussian distribution conditioned on the hidden-layer *class*. **(k)** Workflow for model-based assembly classification and analysis. Raw images (column 1) were up-sampled (column 2), segmented into seven intensity classes via HMRF (column 3), and finally processed by watershed segmentation to define assembly boundaries (column 4). Corresponding line-intensity profiles are shown on the right. (**l**) Schematic of the synthetic assembly image generation and processing pipeline. Ground-truth assemblies were generated, convolved with the experimental PSF, down-sampled, and corrupted with noise to mimic imaging conditions. The synthetic images were then processed through N2V denoising and LR deconvolution. The final output was compared to a directly down-sampled ground-truth reference.

Extended Data Figure 2

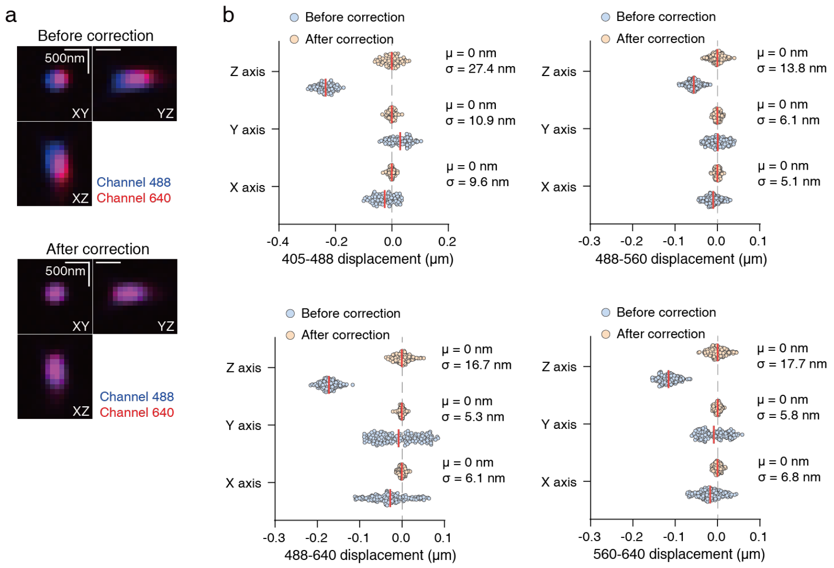


**Extended Data Fig. 2. Chromatic aberration correction.** (**a**) Visualization of chromatic aberration correction using TetraSpeck™ beads. Orthogonal views (XY, XZ, YZ) of fluorescent beads imaged in the 488 nm (blue) and 640 nm (red) channels are shown before (left) and after (right) correction, demonstrating the alignment of signals. (**b**) Quantification of correction efficacy. The displacement vectors between bead centroids localized in different channels are plotted before and after correction. The mean ± standard deviation of the inter-channel displacement is indicated on the graph, confirming a significant reduction post-correction.

#### Extended Data Figure 3

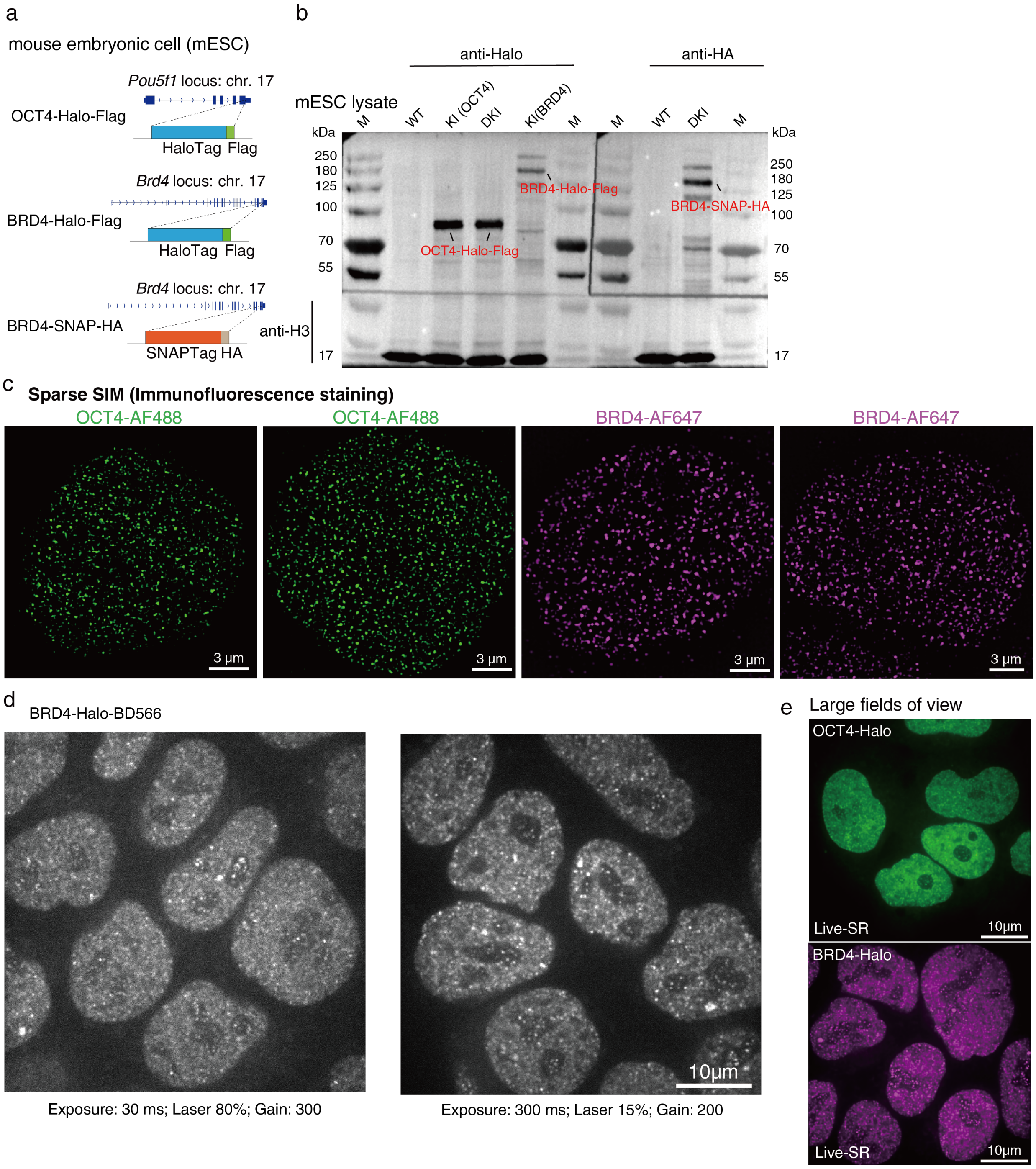


1

Extended Data Figure 3

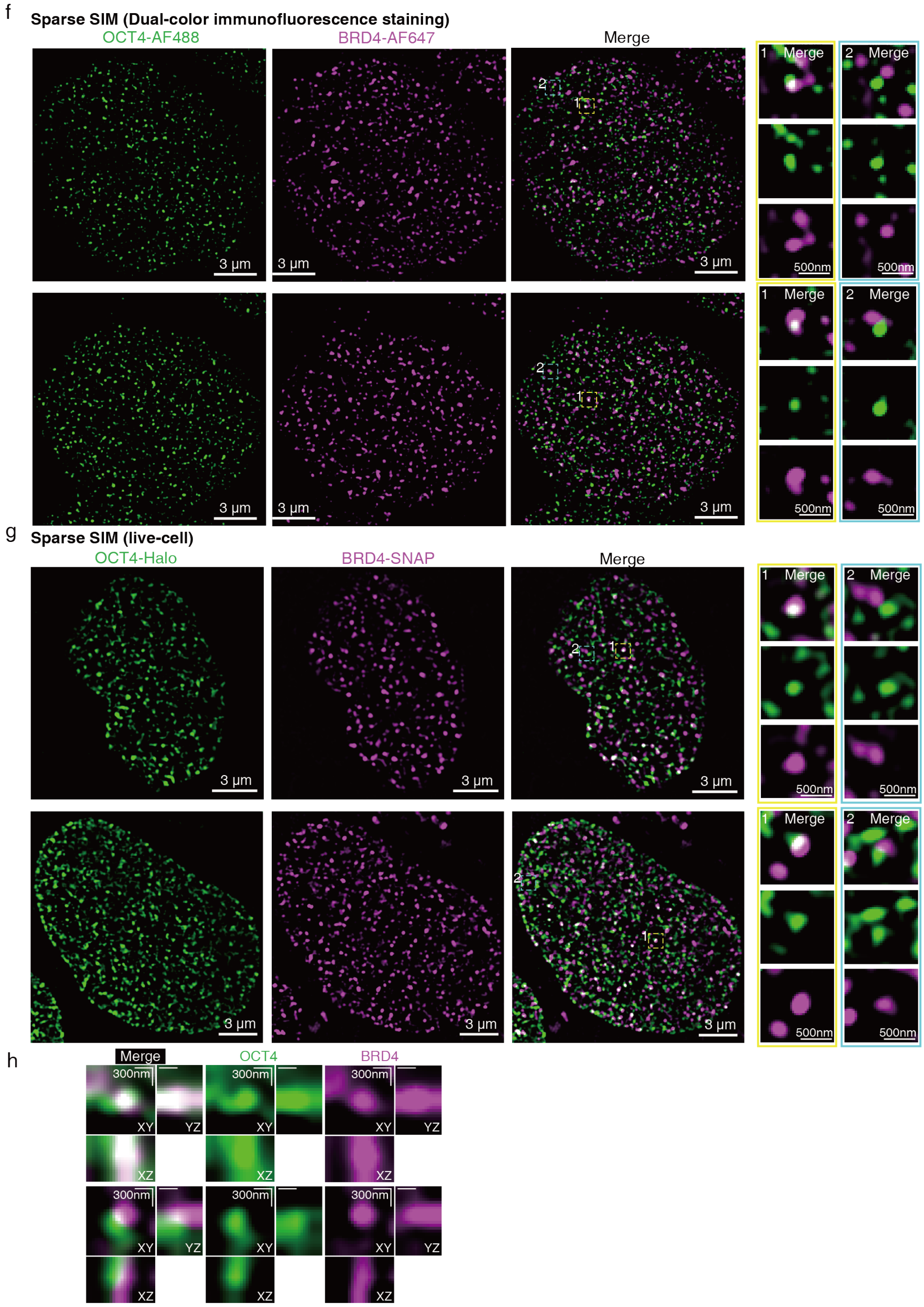


**Extended Data Fig. 3. Validation of endogenous OCT4 and BRD4 knock-in and imaging in mESCs.** (**a**) Schematics of CRISPR/Cas9-mediated HaloTag or SNAP-tag knock-in at the C-terminus of OCT4 or BRD4 in mESCs. (**b**) Western blot of indicated wild-type (WT) and edited mESCs using antibodies against HaloTag (left) and HA tag (right). KI (OCT4): OCT4-Halo-Flag knock-in; KI(BRD4): BRD4-Halo-Flag knock-in; DKI: OCT4-Halo and BRD4-SNAP double knock-in. (**c**) Sparse-SIM images of fixed mESCs immunostained for endogenous OCT4 (green) or BRD4 (magenta) individually. (**d**) Comparison of BRD4 assemblies imaged under low and high exposure conditions. BRD4-Halo was stained with $BD566_{HTL}$. (**e**) Large fields of view of endogenous OCT4-Halo (green pseudocolor) and BRD4-Halo (magenta pseudocolor) in live mESCs. HaloTag was stained with $BD566_{HTL}$. **(f)** Dual-color Sparse-SIM showing colocalization of endogenous OCT4 (green) and BRD4 (magenta) in fixed immunostained mESCs. **(g)** Dual-color live-cell Sparse-SIM of endogenous OCT4 (green) and BRD4 (magenta) in DKI mESCs, labeled with $BD566_{HTL}$ (OCT4-Halo, green) and $BD626_{STL}$ (BRD4-SNAP, magenta) dyes, respectively. **(h)** 3D intensity projections of overlapping (top) and adjacent (bottom) OCT4 and BRD4 assemblies, acquired using the Live-SR module with five-slice z-stacks with a 0.3 μm step size.

Extended Data Figure 4

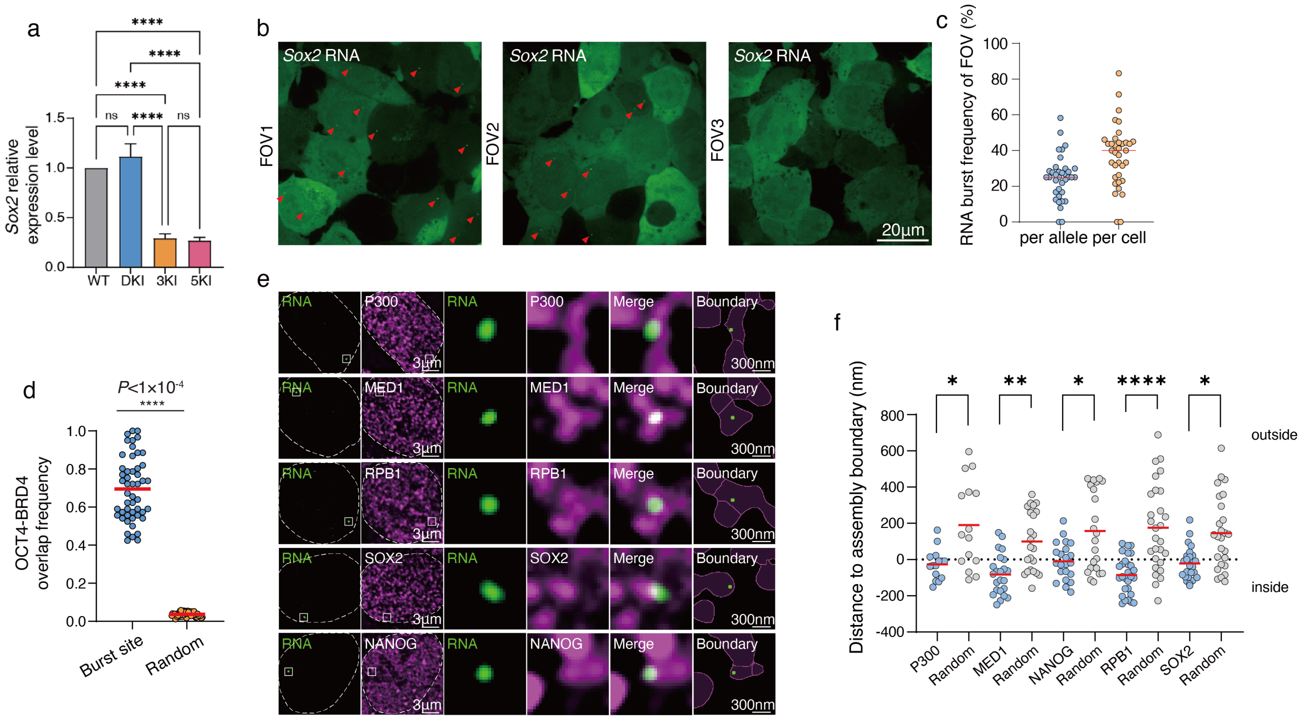


**Extended Data Fig. 4. Imaging and quantitative analysis of transcriptional bursting dynamics and their association with regulatory assemblies. (a)** Relative *Sox2* expression levels in edited cells. RT-qPCR analysis of *Sox2* transcripts, normalized to GAPDH. DKI: OCT4-Halo and BRD4-SNAP; 3KI: OCT4-Halo, BRD4-SNAP, *Sox2*-MS2; 5KI: OCT4-Halo, BRD4-SNAP, *Sox2*-MS2, Prom-CuO, SCR-TetO. **(b)** Representative live-cell snapshots demonstrating transcriptional heterogeneity of *Sox2* RNA bursts (red arrows) across different fields of views (FOVs). **(c)** Fractions of cells and alleles with detectable *Sox2* nascent RNA foci in randomly selected FOV snapshots. n = 37 snapshots. **(d)** Scatter plots of OCT4-BRD4 merging frequency (i.e., merging time proportion) at transcriptional bursting sites versus random sites during time series imaging. n = 50 cells. **(e)** Representative images showing P300, MED1, RPB1, SOX2, and NANOG localization (magenta) relative to *Sox2* RNA (green) in fixed mESCs. **(f)** Boxplots of distances from *Sox2* RNA centroids to assembly boundaries analyzed from (e). Distances from random sites to assembly boundaries were included as a negative control. Sample sizes: P300 (n=15 cells), MED1 (n=23), RPB1 (n=28), SOX2 (n=25), NANOG (n=21), and OCT4/BRD4 (n=18). Statistical significance: one-way ANOVA with Tukey's test; * p-value<0.05, ** p-value<0.01, **** p-value<0.0001, n.s., not significant.

Extended Data Figure 5

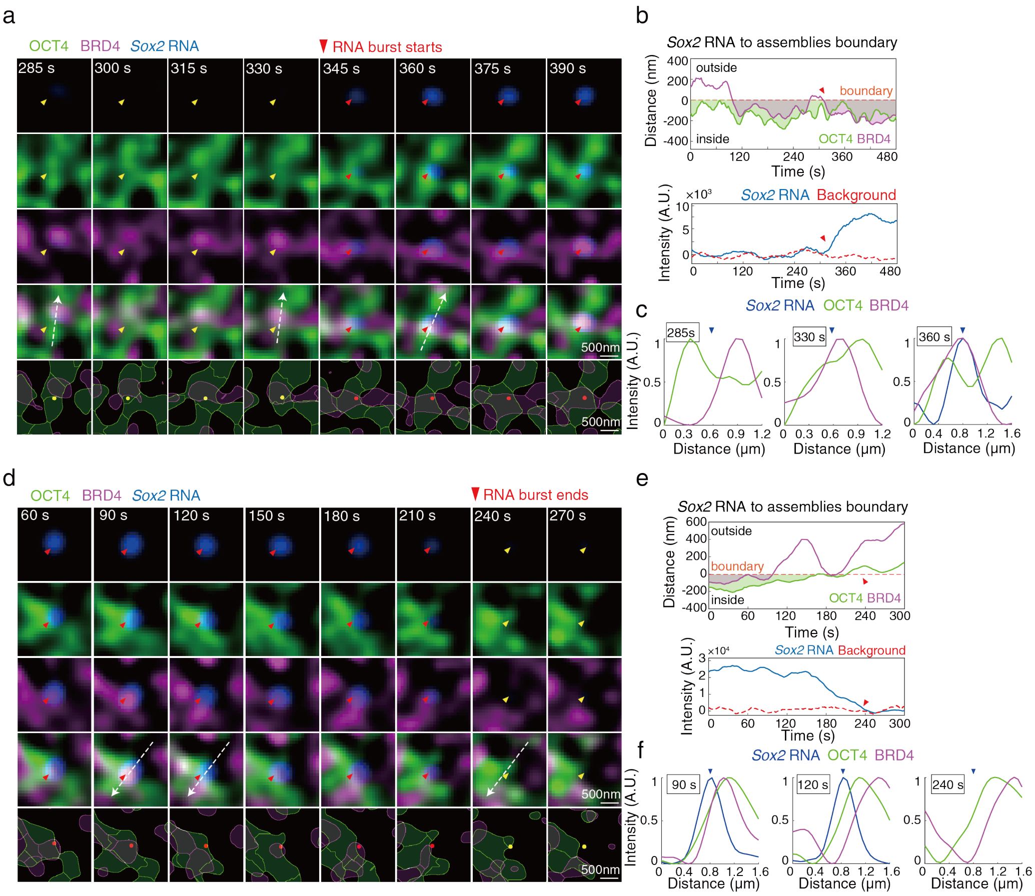


**Extended Data Fig. 5. Temporal and spatial dynamics of OCT4-BRD4 assembly merging during *Sox2* transcriptional bursting.** (**a-f**) Time-lapse confocal imaging capturing burst-onset (**a-c**) or burst-termination (**d-f**) events. Three-plane z-stacks with a step size of 0.3 μm were acquired at 3-second intervals. Images containing the *Sox2* transcription site in focus were shown and analyzed. RNA-to-assembly-boundary distance profiles (**b** and **e**, top), RNA intensity profiles (**b** and **e**, bottom) and line-intensity profiles at selected time points (**c** and **f**) are shown. Images were obtained by three-plane z-stack time-lapse imaging.

#### Extended Data Figure 6

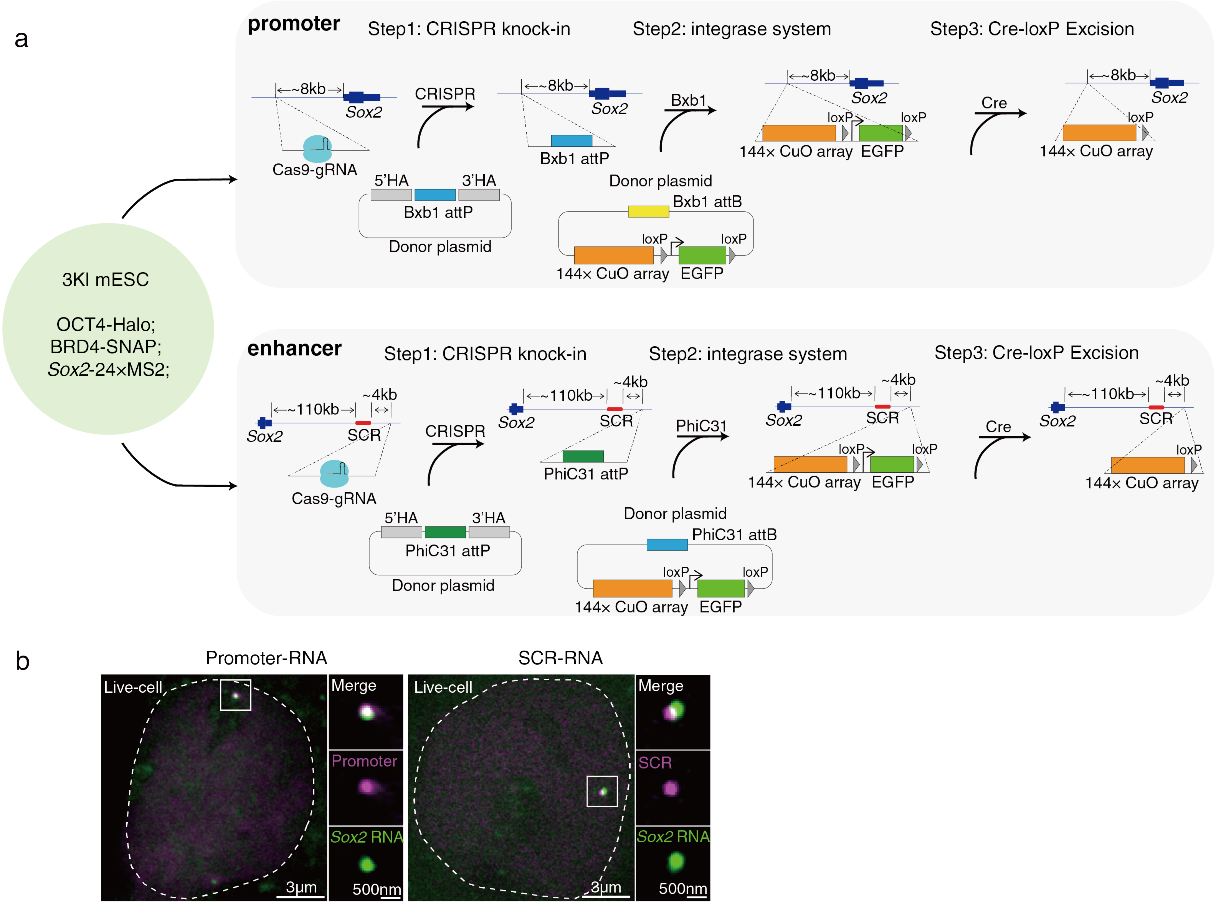


**Extended Data Fig. 6. Simultaneous imaging of *Sox2* DNA and RNA.** (**a**) CRISPR/Cas9-mediated knock-in strategy for inserting a 144×CuO array via Bxb1 attP (*Sox2* promoter) and PhiC31 attP (SCR) recombination sites for DNA labeling. Cre-loxP system was used for excision of EGFP expression cassette. (**b**) Representative live-cell images showing *Sox2* RNA colocalization with either the promoter or SCR.

Extended Data Figure 7

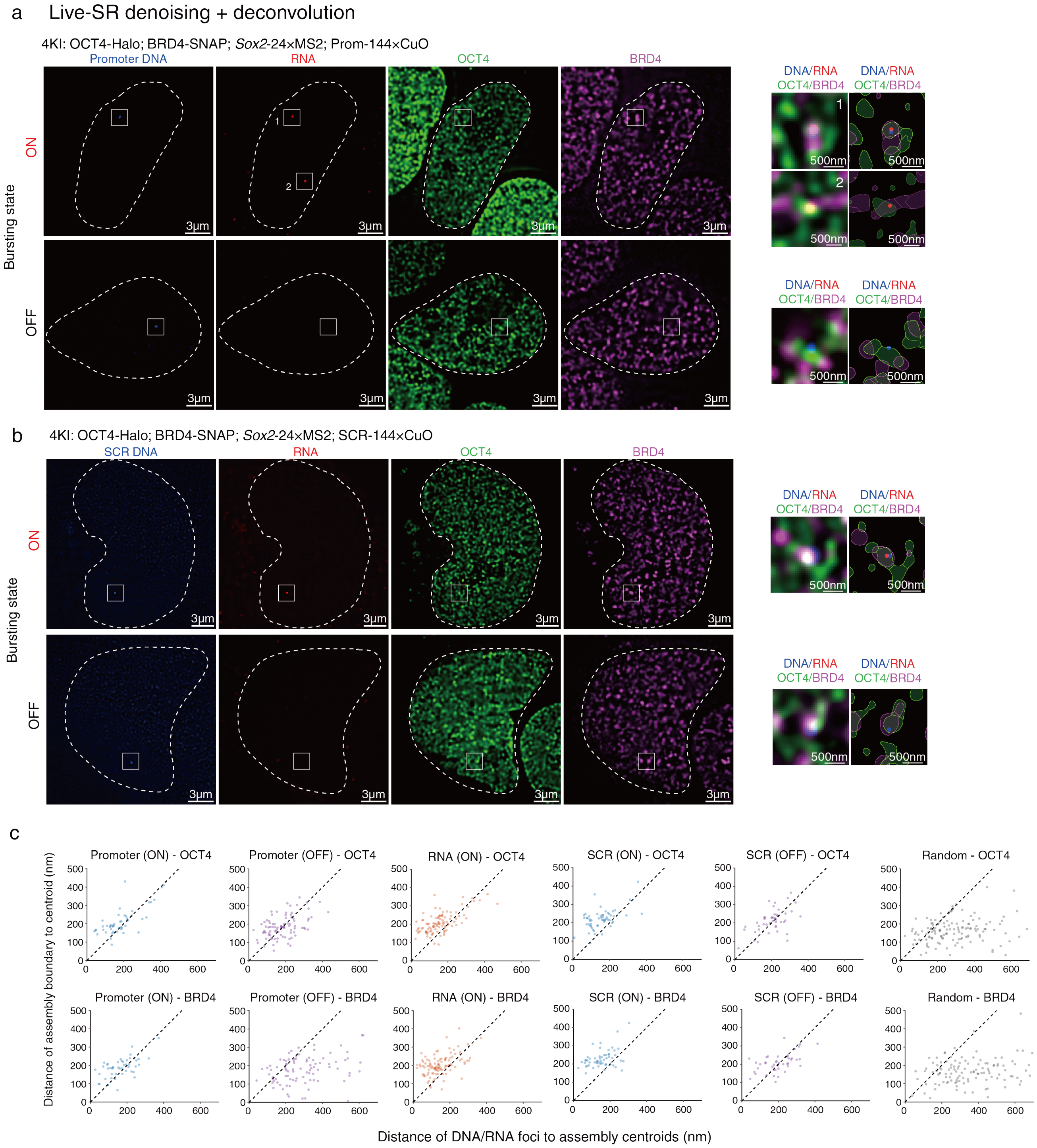


**Extended Data Fig. 7. Quantitative analysis of spatial relationships among *Sox2* DNA, RNA, BRD4 and OCT4 assemblies.** (**a-b**) Whole-cell Live-SR images (processed through denoising and deconvolution) of intact mESCs showing *Sox2* DNA elements (blue; promoter in **a**, SCR in **b**), *Sox2* RNA (red), OCT4 (green), and BRD4 (magenta) assemblies under transcriptionally active (ON) and inactive (OFF) states. ROIs on the right correspond to the magnified views presented in Fig 3b-c. (**c**) Scatter plot of distances from DNA/RNA centroids to assembly centroids versus assembly radii, measured as the distances from assembly boundaries to their centroids, at ON or OFF alleles, corresponding to Fig. 3f. Promoter: n = 48 (ON) and 92 (OFF). RNA: n = 105. SCR: n = 57 (ON) and 44 (OFF).

### Extended Data Figure 8

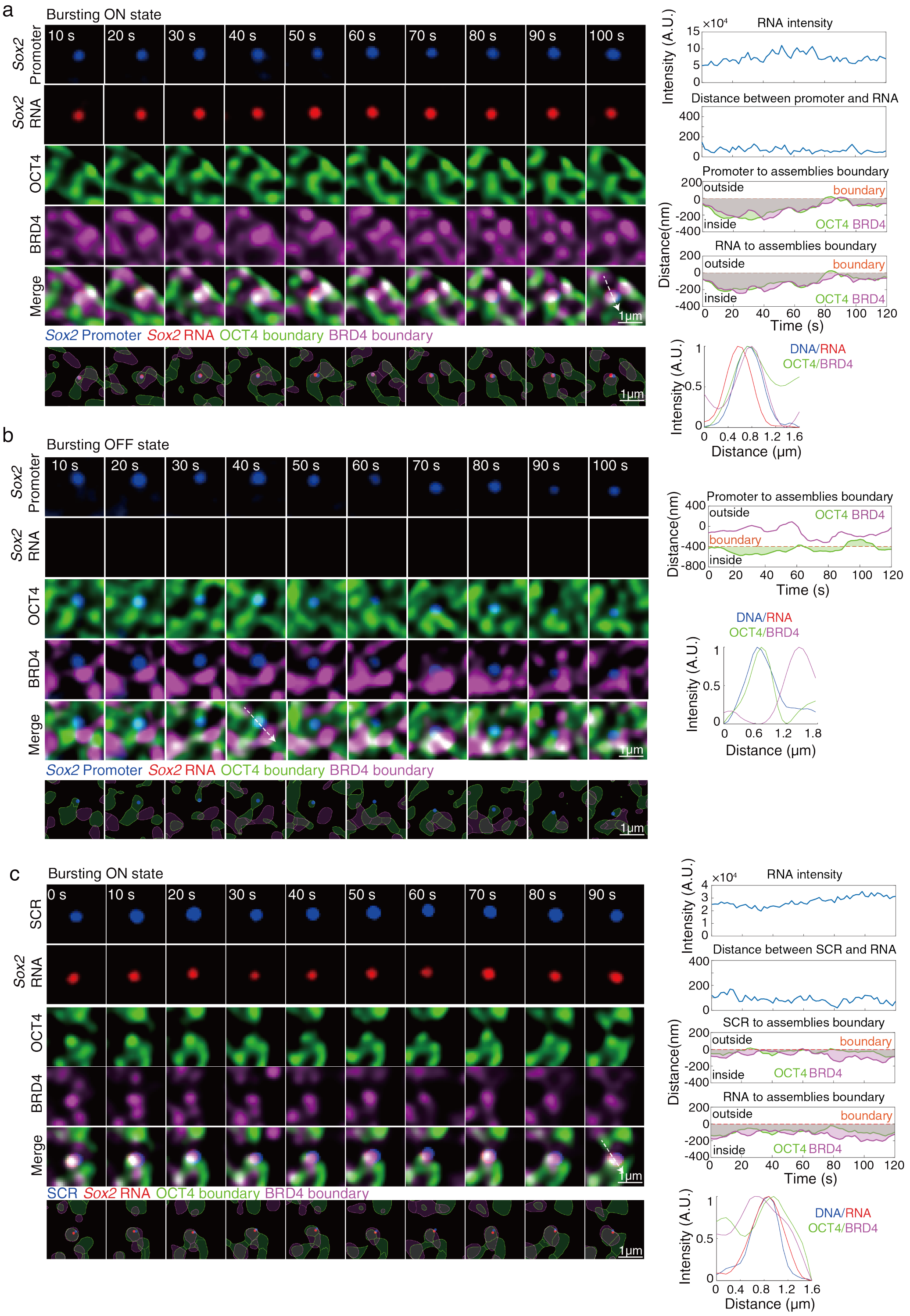

### Extended Data Figure 8

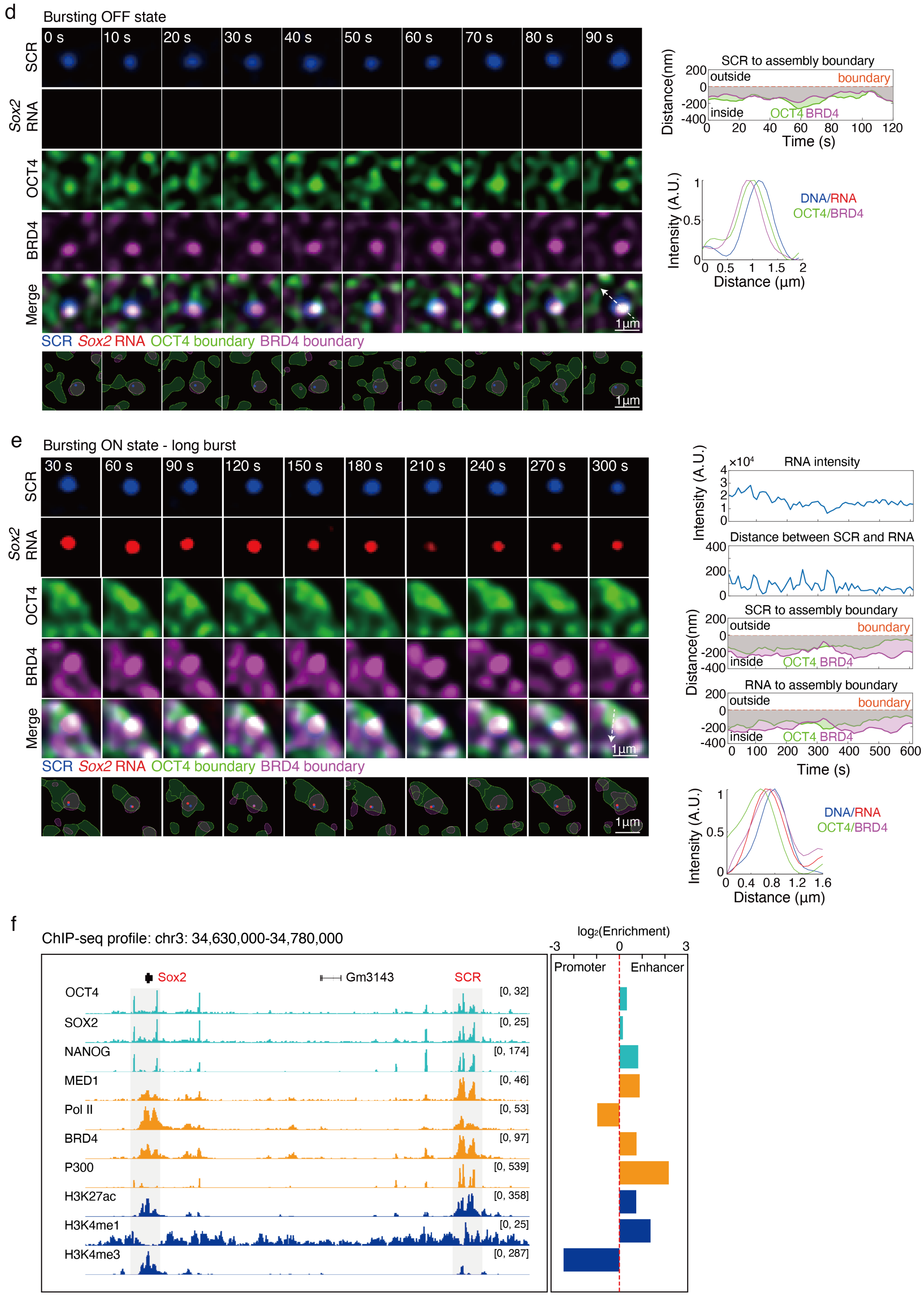

**Extended Data Fig. 8. Representative time-series imaging of *Sox2* transcriptional dynamics and assembly interactions.** (**a-d**) Live-cell time-series imaging of *Sox2* DNA elements (blue, promoter in a and b, and SCR in c and d), *Sox2* RNA (red), OCT4 (green) and BRD4 (magenta) in ON (a, c) and OFF (b, d, corresponding to Fig. 3g) states with a 2-second interval. Resolved *Sox2* DNA centroids, RNA centroids and indicated assembly boundaries are shown at the bottom. RNA intensity profiles, DNA-RNA distance profiles, DNA/RNA-assembly distance profiles and line intensity profiles are shown on the right. Images were obtained through single focal plane time-lapse imaging. In b and d, RNA intensity profiles and DNA-RNA distance profiles were not provided. **(e)** Live-cell time-series imaging of SCR (blue), *Sox2* RNA (red), OCT4 (green) and BRD4 (magenta) in long bursting state with a 10-second interval. **(f)** ChIP-seq enrichment of transcriptional regulators at the *Sox2* promoter-proximal region and SCR. Left, ChIP-seq profiles of selected transcription factors and histone modifications at the *Sox2* locus, analyzed using publicly available GEO datasets. Right, relative enrichment ($\log_2$[SCR/Promoter]) of each factor at the SCR region compared to the promoter region, calculated as the $\log_2$ ratio of average read density of the two regions. Statistical significance: two-sided Student's t-test, ** p-value<0.01, **** p-value<0.0001, n.s., not significant.

#### Extended Data Figure 9

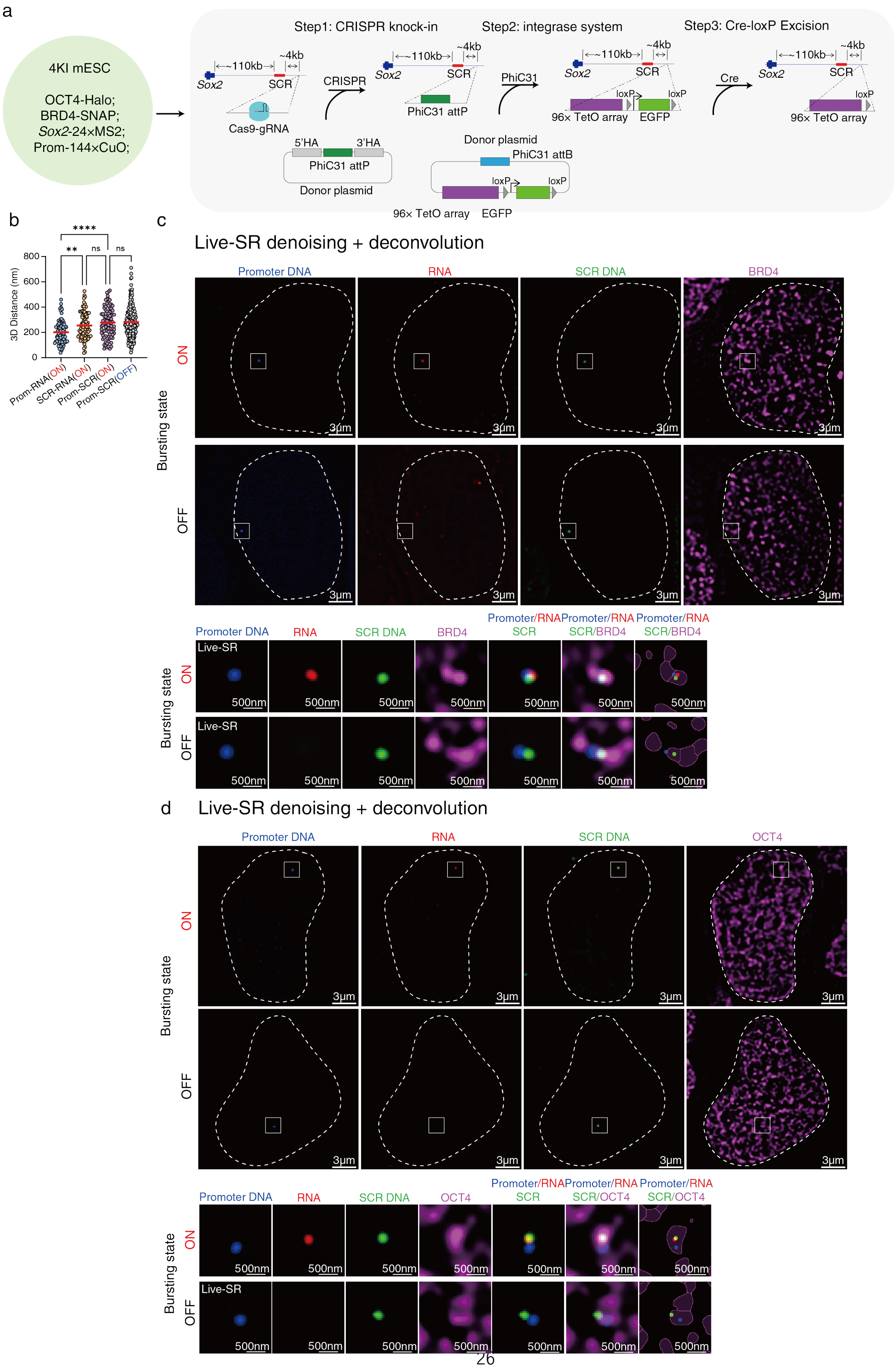


1

**Extended Data Fig. 9. Simultaneous DNA labeling of *Sox2* promoter and SCR.** (**a**) CRISPR/Cas9-mediated knock-in strategy for inserting a 96×TetO array via PhiC31 attP recombination sites to the SCR region for simultaneous DNA labeling of the *Sox2* promoter and SCR. Cre-loxP system was used for excision of EGFP expression cassette. (**b**) 3D distances between promoter-SCR, promoter-RNA, and SCR-RNA at ON alleles (n = 90), and promoter-SCR at OFF alleles (n = 221). (**c-d**) Whole-cell Live-SR images (processed through denoising and deconvolution) of intact mESCs showing *Sox2* promoter DNA (blue), *Sox2* RNA (red), SCR DNA (green), and assemblies (magenta; BRD4 in c, OCT4 in d) under transcriptionally active (ON) and inactive (OFF) states. ROIs on the bottom correspond to the magnified views presented in Fig 4**f, g**.

### Extended Data Figure 10

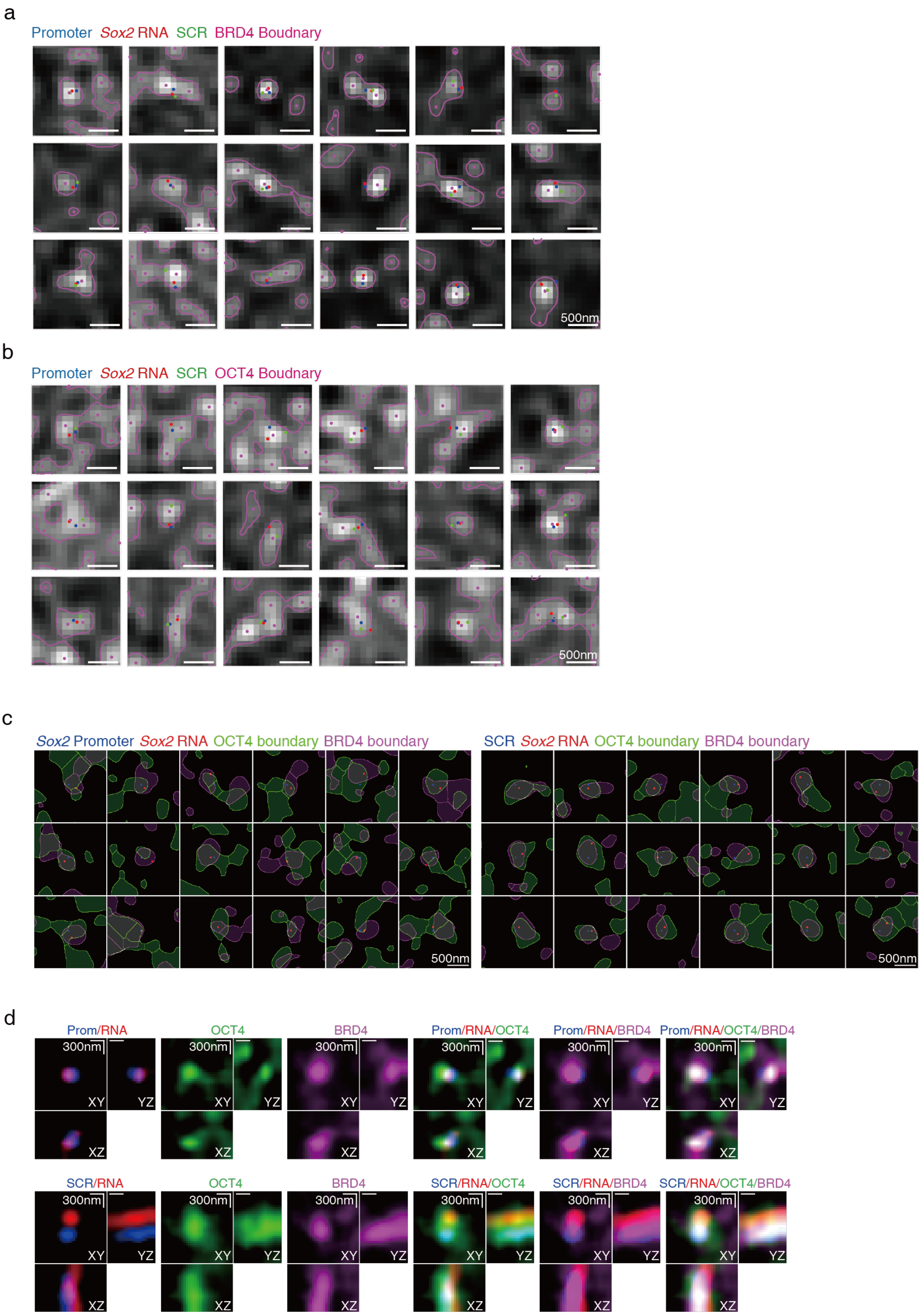

**Extended Data Fig. 10. Examples of the *Sox2* promoter, RNA and SCR within shared OCT4- and BRD4-enriched assemblies.** (**a**) Resolved *Sox2* promoter centroids (blue), RNA centroids (red), SCR centroids (green) and BRD4 (magenta) assembly boundaries. (**b**) Resolved *Sox2* promoter centroids (blue), RNA centroids (red), SCR centroids (green) and OCT4 (magenta) assembly boundaries. **(c)** Resolved *Sox2* DNA centroids (blue; left: promoter, right: SCR), RNA centroids (red), OCT4 (green) and BRD4 (magenta) assembly boundaries. **(d)** 3D intensity projection of a representative cell, displaying SCR and RNA foci associated with overlapping OCT4- and BRD4-enriched assemblies. Images in (**a-b**) were acquired with dual-DNA imaging. This strategy visualized both the *Sox2* promoter and SCR DNA locus, along with RNA and one assembly (BRD4 or OCT4), to assess their association within a single assembly. Images in (**c-d**) were acquired with dual-color assembly imaging. This strategy visualized one DNA locus (*Sox2* promoter or SCR), RNA and both OCT4 and BRD4 assemblies, to assess their association with an overlapping OCT4- and BRD4-enriched assembly environment.

#### Extended Data Figure 11

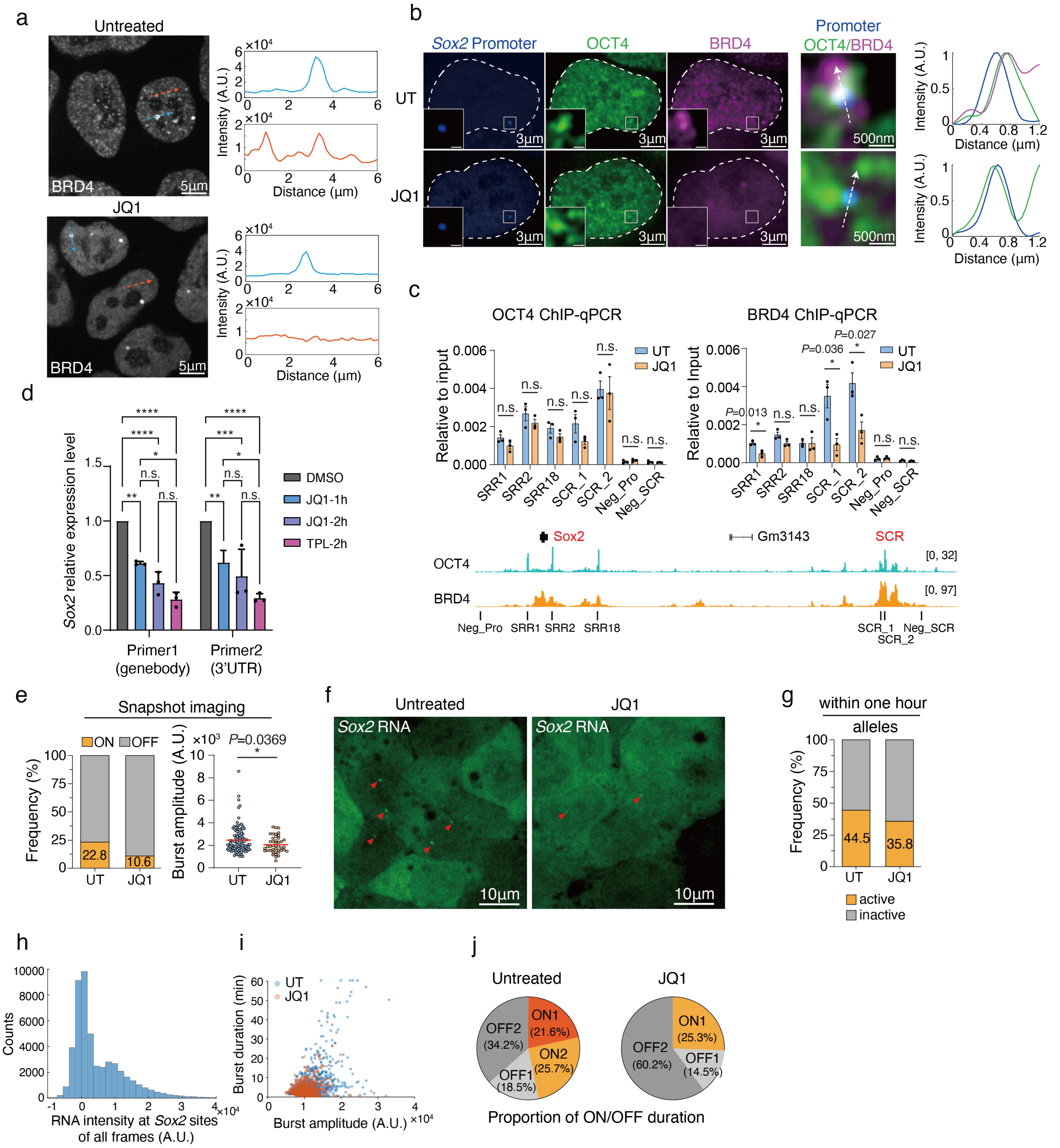


**Extended Data Fig. 11. Quantitative characterization of *Sox2* bursting and assembly behavior under JQ1 treatment.** (**a**) Live-SR images of endogenous BRD4-Halo assemblies in UT or JQ1-treated (1 μM for 30 min) cells. Line intensity profiles crossing large assemblies (blue arrow) or small assemblies (orange arrow) were shown on the right. (**b**) Live-cell imaging of *Sox2* promoter (blue), OCT4 (green) and BRD4 (magenta) assemblies in UT or JQ1-treated (1 μM for 30 min) cells. (**c**) Top, ChIP-qPCR results of OCT4 and BRD4 in untreated (UT) or JQ1-treated (1 μM for 30 min) cells, normalized to sonicated chromatin input. Bottom, qPCR primers at indicated genomic locations. (**d**) Relative *Sox2* expression levels in UT versus JQ1-treated (1 μM for 1 or 2 hours) and TPL-treated (1 μM for 2 hours) cells. RT-qPCR analysis of *Sox2* gene body and 3'UTR transcripts, normalized to

GAPDH. (**e**) Allele-level quantification of the fraction with detectable *Sox2* nascent RNA foci (left) and RNA-focus amplitude (right) in untreated (n = 298) or JQ1-treated (n = 267) cells. (**f**) Representative fields of view showing *Sox2* RNA bursts (red arrows) in UT or JQ1-treated cells. Images were smoothed for visualization purposes only. (**g**) Fraction of *Sox2* alleles classified as active during the one-hour imaging window. (**h**) Frequency distribution of RNA fluorescence intensities at *Sox2* genomic loci across all imaging frames. Histogram displays the counts of observed RNA signal intensities at single-allele resolution, facilitating HMM threshold setting. (**i**) Scatter plot of burst amplitude versus duration in untreated and JQ1-treated cells, corresponding to Fig. 5e. (**j**) Pie charts showing the integrated time-occupancy of each state, with sector sizes proportional to the product of duration and fraction. Statistical significance: two-sided Student's t-test, * p-value<0.05, **** p-value<0.0001, n.s., not significant.

**Supplementary Table 1**

sgRNAs and primers used in this study

| Name | Sequence | Usage |
|---|---|---|
| OCT4-sgRNA | TCTCCCATGCATTCAAACTG | Integrating HaloTag |
| BRD4-sgRNA | AATCTTTTTTGAGAGCACCC | Integrating HaloTag/SNAP-tag |
| *Sox2*-sgRNA | CCAGCCCTCACATGTGCGAC | Integrating MS2 to *Sox2* 3'UTR |
| *Sox2*-promoter | TATGTTAAGCTAGTTTCTGA | Integrating Bxb1 attP to *Sox2* promoter for CuO array knock in |
| *Sox2*-SCR | GTAAGCTATCTCATTGCCCG | Integrating PhiC31 attP to *Sox2* SCR for CuO array knock in |
| *Brd4*-5HA-forward primer | GCAGGAGCCAAAAACACCAGT | Genomic PCR for homology arms |
| *Brd4*-5HA-reverse primer | AAAAAGATTTTCTTCAAATATTGACAAAAGATCACTCTGG | Genomic PCR for homology arms |
| *Brd4*-3HA-forward primer | GAGCACCCAGGTGACTTCTGAC | Genomic PCR for homology arms |
| *Brd4*-3HA-reverse primer | TGACTTTGGTTGGTGGCCTTACT | Genomic PCR for homology arms |
| *POU5F1*-5HA-forward primer | GTGGAGTCTGGAGACCATGTT | Genomic PCR for homology arms |
| *POU5F1*-5HA-reverse primer | GTTTGAATGCATGGGAGAGCCC | Genomic PCR for homology arms |
| *POU5F1*-3HA-forward primer | TGAGGCACCAGCCCTCCC | Genomic PCR for homology arms |
| *POU5F1*-3HA-reverse primer | AGAACTCCCAGAGTGACAAGAGG | Genomic PCR for homology arms |
| *Sox2*-promoter-5HA-forward primer | GAAGACTCCTTCGCCGACAA | Genomic PCR for homology arms |
| *Sox2*-promoter-5HA-reverse primer | TCAGAAACTAGCTTAACATATCCTAACA | Genomic PCR for homology arms |
| *Sox2*-promoter-3HA-forward primer | TGGCAAGCCCTAACCTTGC | Genomic PCR for homology arms |
| *Sox2*-promoter-3HA-reverse primer | CCTCGCCTCAAACCACTCTT | Genomic PCR for homology arms |
| *Sox2*-SCR-5HA-forward primer | TCGGGAAACGTGTGTCCAAT | Genomic PCR for homology arms |
| *Sox2*-SCR-5HA-reverse primer | CGGGCAATGAGATAGCTTACTG | Genomic PCR for homology arms |
| *Sox2*-SCR-3HA-forward primer | AGGATGCTTACAGTTCTAACTAATCACC | Genomic PCR for homology arms |
| *Sox2*-SCR-3HA-reverse primer | TGGAACACAATAGTGCTTCATTTG | Genomic PCR for homology arms |

**Supplementary Table 2**

Primers used for quantitative PCR

| Primer | Sequence | Usage |
|---|---|---|
| *Sox2* gene body forward primer | GATCAGCATGTACCTCCCCG | RT-qPCR |
| *Sox2* gene body reverse primer | GGGGCAGTGTGCCGTTAAT | RT-qPCR |
| *Sox2* 3'UTR forward primer | GGAAAGGGTTCTTGCTGGGT | RT-qPCR |
| *Sox2* 3'UTR reverse primer | ACGAAAACGGTCTTGCCAGT | RT-qPCR |
| GAPDH forward primer | GTGTTCCTACCCCCAATGTGT | RT-qPCR |
| GAPDH reverse primer | AATGTGATACCAGGAAATGAGCTT | RT-qPCR |
| SRR1 forward primer | AAGATGCTTGGCTAGTTCTCG | ChIP-qPCR |
| SRR1 reverse primer | TGTGGAGGTGCGGACTGTA | ChIP-qPCR |
| SRR2 forward primer | CGACAGTTCTTGCTCACACA | ChIP-qPCR |
| SRR2 reverse primer | TCCATTGGCTGGAGTTCCG | ChIP-qPCR |
| SRR18 forward primer | GTGGGTGACTCCATTCTGTCTT | ChIP-qPCR |
| SRR18 reverse primer | AGATTGGCAGCAGCTCGGA | ChIP-qPCR |
| SCR1 forward primer | CCCAGTCCAAAGCGAATACT | ChIP-qPCR |
| SCR1 reverse primer | GCTTCTCCGATGGAAATCTGC | ChIP-qPCR |
| SCR2 forward primer | CAGGACTCTGCTAAGGGCAA | ChIP-qPCR |
| SCR2 reverse primer | TCTGGGTGGTGAAACCTTGG | ChIP-qPCR |
| Neg_pro forward primer | GCACCTCAGTGGCTCTTCAT | ChIP-qPCR |
| Neg_pro reverse primer | GGCCAGACACCCGATTGTTA | ChIP-qPCR |
| Neg_SCR forward primer | GCAATGAGGAAGAAGTTGGCA | ChIP-qPCR |
| Neg_SCR reverse primer | GCCTGGTTCACACAATAGCG | ChIP-qPCR |

**Supplementary Table 3**

Antibodies used in this study

| Name | Source | Identifier | Usage | Dilution |
|---|---|---|---|---|
| Anti-BRD4 | Bethyl | A301-985A100 | ChIP-qPCR | 1:1000 |
| Anti-OCT4 | Abcam | AB181557 | ChIP-qPCR | 1:1000 |
| Anti-RPB1 | Cell signaling | 14958S | Immunofluorescence staining | 1:200 |
| Anti-MED1 | Bethyl | A300-793A | Immunofluorescence staining | 1:200 |
| Anti-P300 | Santa Cruz Biotechnology | sc48343 | Immunofluorescence staining | 1:200 |
| Anti-SOX2 | Abcam | AB97959 | Immunofluorescence staining | 1:200 |
| Anti-NANOG | R&D systems | AF1997 | Immunofluorescence staining | 1:200 |
| Alexa Fluor 555 anti-mouse | Invitrogen | A21422 | Immunofluorescence staining | 1:500 |
| Alexa Fluor 647 anti-rabbit | Invitrogen | A31573 | Immunofluorescence staining | 1:500 |
| Anti-HaloTag | Promega | G921A | Western blot | 1:500 |
| Anti-HAtag | Abcam | ab9110 | Western blot | 1:500 |
| Anti-H3 | Beyotime | AH433 | Western blot | 1:2000 |
| Goat Anti-Rabbit IgG (H&L)-HRP Conjugated secondary antibody | EASYBIO | BE0101 | Western blot | 1:10000 |
| Goat Anti-Mouse IgG (H&L)-HRP Conjugated secondary antibody | EASYBIO | BE0102 | Western blot | 1:10000 |

**Video S1. Dynamic merging and separation of OCT4 and BRD4 assemblies.**

This video demonstrates the dynamic behavior of endogenous OCT4 (green) and BRD4 (magenta) assemblies in live mESCs, captured at 1-second intervals using Live-SR imaging. Arrows highlight representative homotypic merging and separation events. Images were processed with N2V denoising.

**Video S2. 3D Sparse-SIM of OCT4 and BRD4 assemblies**

This video demonstrates a 3D view of simultaneously imaged endogenous OCT4 and BRD4 assemblies. Representative Sparse-SIM images and segmented asseblies are shown. BRD4-Halo was labeled with $BD566_{HTL}$ and OCT4 immunostained with AF488.

**Video S3. Heterotypic interactions between OCT4 and BRD4 assemblies.**

Two-color Live-SR imaging at 2-second intervals reveals dynamic proximity, partial overlap, and separation between OCT4 and BRD4 assemblies. Key interaction events were marked in the boxed regions. Images were processed with Condensate-quant.

**Video S4. Time-lapse imaging of transcriptional bursting at two *Sox2* alleles in the same cell.**

Time-lapse confocal imaging of *Sox2* RNA at 30-second intervals for 1 hour. Real-time RNA intensities of the two alleles are shown on the right.

**Video S5. Three-color live-cell imaging of *Sox2* nascent RNA and assemblies.**

Time-lapse confocal imaging of *Sox2* RNA (blue), OCT4 (green) and BRD4 (magenta) at 2-second intervals. The bottom-right panel displays resolved *Sox2* RNA centroids and OCT4/BRD4 assembly boundaries. Images were processed with Condensate-quant.

**Video S6. Time-lapse confocal imaging of a burst-onset events**

Three-color live-cell imaging of *Sox2* RNA (blue), OCT4 (green) and BRD4 (magenta) at 3-second intervals. The bottom-right panel displays RNA centroids relative to assembly boundaries. Red dots indicate the positions of bursting RNA, whereas yellow dots indicate the inferred locus positions during non-bursting periods, estimated by linear interpolation between the positions of the preceding and subsequent bursts. Images were processed with Condensate-quant.

**Video S7. Time-lapse confocal imaging of a burst-termination events.**

Three-color live-cell imaging of *Sox2* RNA (blue), OCT4 (green) and BRD4 (magenta) at 3-second intervals. The bottom-right panel displays RNA centroids relative to assembly boundaries. Red dots indicate the positions of bursting RNA, whereas yellow dots indicate the inferred locus positions during non-bursting periods, estimated by linear interpolation between the positions of the preceding and subsequent bursts. Images were processed with Condensate-quant.

**Video S8. Four-color live-cell imaging of a bursting Sox2 allele with promoter labeling.**

Time-lapse confocal imaging of the *Sox2* promoter label (blue), *Sox2* RNA (red), OCT4 (green) and BRD4 (magenta) at 2-second intervals at a bursting allele. The bottom-right panel displays resolved promoter-label, RNA centroids and OCT4/BRD4 assembly boundaries. Images were processed with Condensate-quant.

**Video S9. Four-color live-cell imaging of a non-bursting Sox2 allele with promoter labeling.**

Time-lapse confocal imaging of the *Sox2* promoter label (blue), *Sox2* RNA channel (red; no detectable RNA focus), OCT4 (green) and BRD4 (magenta) at 2-second intervals at a non-bursting allele. Images were processed with Condensate-quant.

**Video S10. Four-color live-cell imaging of a bursting Sox2 allele with SCR labeling.**

Time-lapse confocal imaging of the SCR label (blue), *Sox2* RNA (red), OCT4 (green) and BRD4 (magenta) at 2-second intervals at a bursting allele. Images were processed with Condensate-quant.

**Video S11. Four-color live-cell imaging of a non-bursting Sox2 allele with SCR labeling.**

Time-lapse confocal imaging of the SCR label (blue), *Sox2* RNA channel (red; no detectable RNA focus), OCT4 (green) and BRD4 (magenta) at 2-second intervals at a non-bursting allele. Images were processed with Condensate-quant.

**Video S12. Four-color live-cell imaging during a long *Sox2* transcriptional burst.**

Time-lapse confocal imaging of the SCR lable (blue), *Sox2* RNA (red), OCT4 (green) and BRD4 (magenta) at 10-second intervals for 600 s. Images were processed with Condensate-quant.

**Video S13. Long-term time-lapse imaging of *Sox2* bursts with or without JQ1 treatment.**

Live-cell confocal imaging of *Sox2* RNA bursts (red arrows) in untreated (control) or JQ1-treated cells, imaged over 1 hour at 30-second intervals. Maximum-intensity projections of nine z-slices are shown, with smoothing applied for visualization only.
