## Supplementary notes for "Dynamic merging of enhancer- and promoter-associated assemblies is coupled to transcriptional bursting"

### Supplementary Note 1: Rationale and validation of assembly segmentation

Accurate segmentation of nuclear assemblies is critical for downstream quantitative analysis. In this study, raw images were first processed using the *Condensate-quant* pipeline, consisting of Noise2Void (N2V) denoising followed by Lucy-Richardson (LR) deconvolution. The resulting images were then up-sampled to a pixel size of 9.5 nm for Live-SR data or 10 nm for confocal data. We subsequently employed a hidden Markov random field (HMRF) model to classify pixel intensities and segment assemblies. We systematically evaluated its performance using simulated and experimental data and compared it with alternative thresholding methods.

#### HMRF-based assembly segmentation

To quantitatively characterize intensity heterogeneity within the nucleus, we employed an HMRF model to classify pixels into seven discrete intensity classes. Seven classes were selected as a balance between model flexibility and robustness, providing sufficient resolution to distinguish background, diffuse signal, and regions of local enrichment while avoiding overfitting. This approach was adapted from previously established segmentation methods for chromatin organization[1,2].

The HMRF model assumes that the observed pixel intensity $H_i$ follows a Gaussian distribution conditioned on its latent class label, such that each class is characterized by a mean$c_i$ $\mu_c$ and variance $\sigma_c^2$. Under this formulation, the global intensity distribution is effectively modeled as a Gaussian mixture (Fig. 1). In addition, the model incorporates spatial coherence by assuming that neighboring pixels are more likely to belong to the same class. This spatial prior enforces local smoothness and suppresses noise-driven fluctuations, resulting in more stable segmentation and smoother class boundaries.

The classification is formulated within a Bayesian framework. Model parameters were estimated using an expectation–maximization (EM) approach combined with iterative optimization of the spatial prior. Specifically, the log-likelihood function is defined as:

$$\mathcal{L}_i = -\left(H_i - \mu_{c_i}\right)^2/\left(2\sigma_{c_i}^2\right) + \beta \sum I\left(c_i \neq c_j\right)$$

The second term encodes spatial regularization, $i$ and $j$ denote neighboring pixels and $I$ is an indicator function. The parameter $\beta > 0$ controls the strength of spatial regularization. Unlike conventional threshold-based methods, the HMRF approach does not rely on predefined intensity cutoffs. Instead, it provides a data-driven and adaptive classification of intensity levels, enabling robust segmentation across images with heterogeneous backgrounds and varying signal distributions.

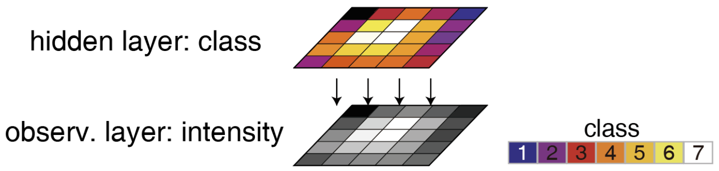


**Supplementary Figure. 1 Schematic of the hidden Markov random field (HMRF)-based segmentation algorithm.** The observation-layer *intensity* is modeled as a Gaussian distribution conditioned on the hidden-layer *class*.

Gaussian mixture modeling of the intensity distribution for real image data revealed that the global mean intensity lies approximately within class 3, indicating that lower classes primarily correspond to background or diffuse signal, whereas classes 4-7 represent signal-enriched regions (Fig. 2). Based on the separation of mixture components, pixels assigned to the selected highest-intensity classes (in this study, classes 5 to 7) were designated as assemblies; these classes correspond to intensity modes with minimal overlap with background-dominated classes. This stringent criterion prioritizes high-confidence signal regions while minimizing inclusion of weak background fluctuations and ambiguous boundary pixels.

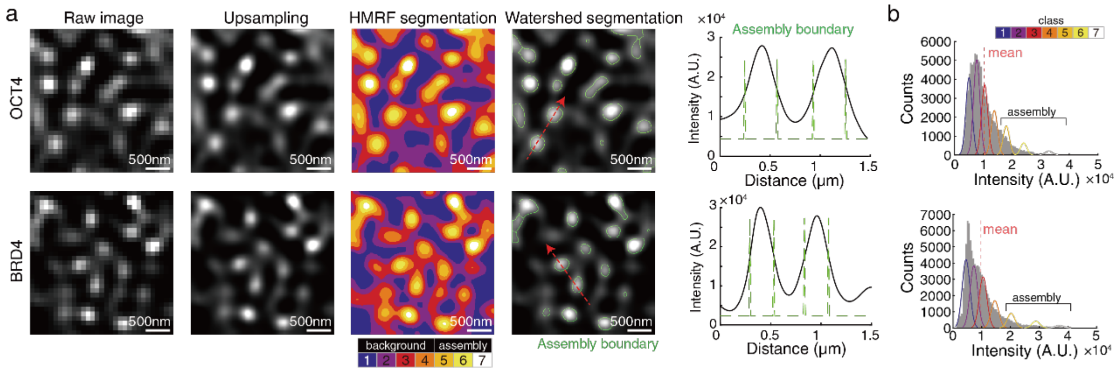


**Supplementary Figure. 2 Workflow for model-based assembly classification and analysis.** (a) Raw images (column 1) were up-sampled (column 2), segmented into seven intensity classes via HMRF (column 3), and finally processed by watershed segmentation to define assembly boundaries (column 4). Corresponding line intensity profiles are shown on the right. (b) Histogram of pixel intensities overlaid with probability density functions of the seven Gaussian components generated by the HMRF model.

#### Simulation-based validation of segmentation performance

To validate the reliability of the segmentation method, we simulated synthetic assembly images with known ground truth, adapting a previously reported pipeline[3]. Briefly, 3D assemblies were simulated with varying physical radii (100-200 nm) and signal-enrichment levels (partition coefficients of 3-7). The ground-truth assemblies were convolved with an experimentally measured PSF, and signals within the depth of focus were projected onto the 2D imaging plane. The resulting images were then downsampled to the camera pixel size and corrupted with Gaussian and Poisson noise, as well as uneven illumination, to mimic experimental imaging conditions. Assembly density and spatial proximity were set to approximate realistic nuclear environments. To account for varying imaging qualities, we set signal-to-noise ratios (SNRs) of 20 dB and 24 dB (Fig. 3a, b).

To evaluate robustness, we compared HMRF-based segmentation with two representative threshold-based segmentation approaches, global Otsu thresholding and local adaptive thresholding. After applying identical N2V denoising and deconvolution preprocessing, segmentation was performed using all three methods and compared against the ground truth.

Segmentation performance was evaluated using the Dice similarity coefficient and boundary error metrics. Boundary error was defined as the difference between the equivalent radius derived from the segmented mask and the ground-truth radius. Negative values reflect conservative boundary estimation, indicating reduced inclusion of background signal. All three methods achieved relatively high Dice similarity scores, indicating accurate overall detection of assembly regions (Fig. 3d). However, boundary analysis revealed clear differences: HMRF produced more stringent and consistent segmentation, as evidenced by a smaller variance in boundary error across noise conditions (Fig. 3c).

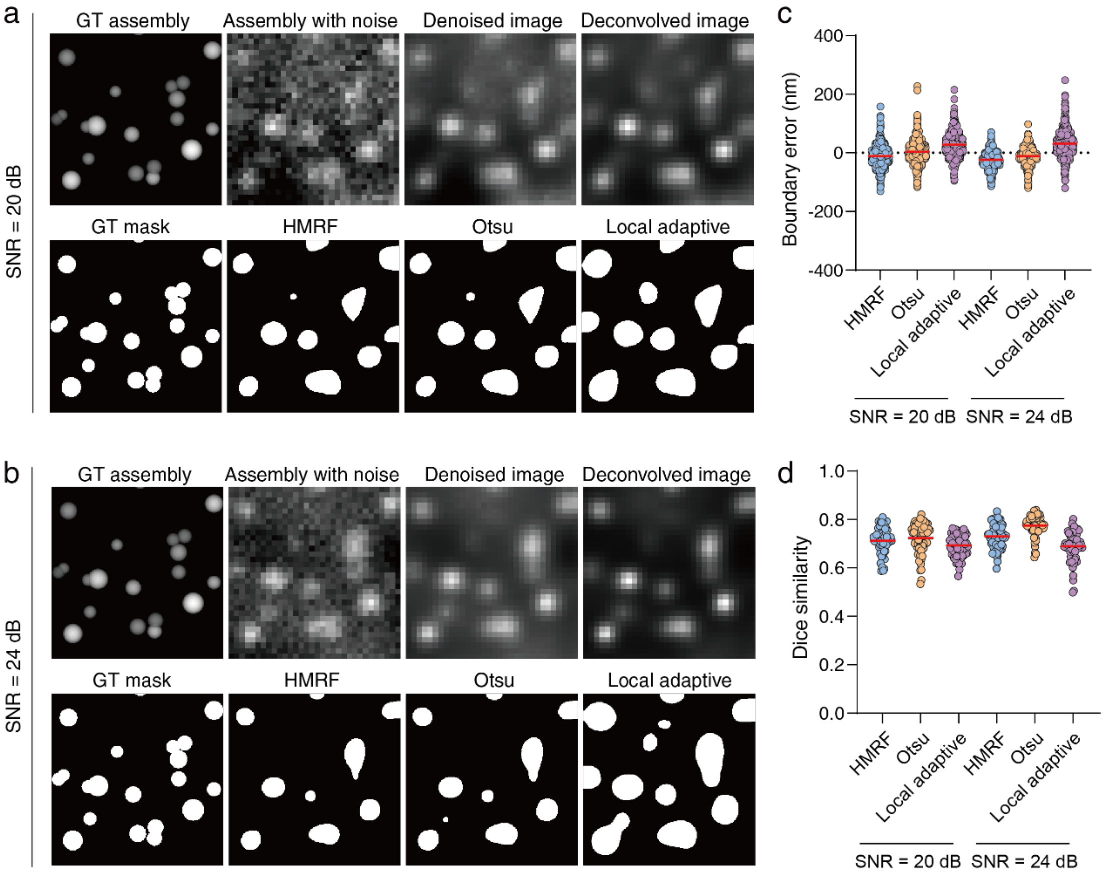


**Supplementary Figure. 3 Comparison of assembly segmentation methods using simulated data.** (a) (Top) Ground-truth assemblies were generated and synthetic images were created at an SNR of 20 dB by convolution with the experimental PSF and addition of noise. The images were then processed using N2V denoising and LR deconvolution. (Bottom) Comparison of ground-truth masks with assembly segmentation results obtained using HMRF, Otsu thresholding, and local adaptive thresholding. (b) Same as in (a), but for SNR = 24 dB. (c) Boundary error across different segmentation methods and noise levels. (d) Dice similarity coefficients across different segmentation methods and noise levels.

**Performance on experimental data**

Consistent with the simulation results, we observed that while Otsu-based methods perform reasonably well under high signal conditions, they are less robust in more complex imaging scenarios (Fig. 4). In particular, when signal contrast is low or when large low-intensity regions (e.g., nucleoli) are present, global thresholding becomes sensitive to background heterogeneity. This often leads to over-segmentation or merging of adjacent assemblies.

In contrast, the HMRF approach incorporates both spatial context and multi-class intensity modeling, enabling more robust segmentation in heterogeneous nuclear environments. This is particularly important for dense assemblies or conditions with non-uniform background, where accurate boundary delineation is critical for downstream quantitative analysis.

Accordingly, the segmented boundaries are used as consistent, intensity-based operational definitions for comparative spatial analyses and should not be interpreted as sharp physical interfaces or as evidence of a particular material state.

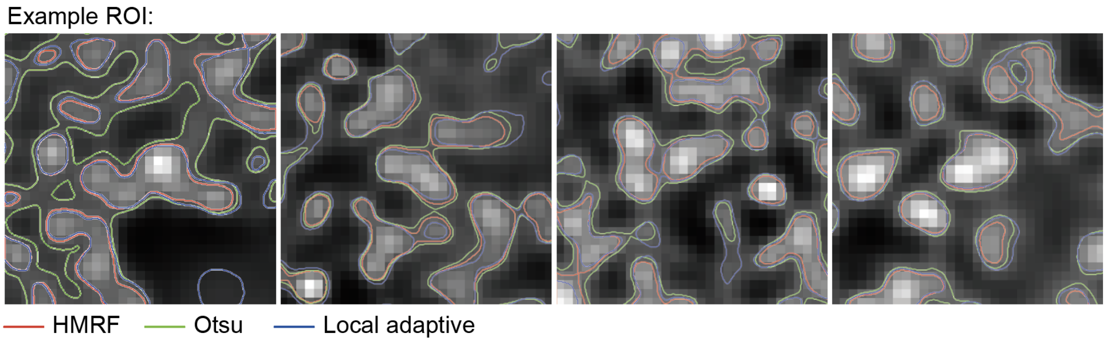


**Supplementary Figure. 4 Comparison of assembly segmentation methods in representative regions of interest (ROIs).** Grayscale images show assembly signals with overlaid boundaries obtained using HMRF, Otsu thresholding, and local adaptive thresholding.

### Supplementary Note 2: Quantification of localization accuracy for assembly centroids and DNA/RNA foci

#### Accuracy of centroid localization across PSF sizes

Boundary-based and centroid-based measurements provide complementary approaches for assessing spatial proximity. In Supplementary Note 1, we describe our strategy for obtaining reliable boundary measurements despite the effects of PSF broadening and image noise. Here, we focus on centroid localization, which is generally less sensitive to variations in apparent object size because the center position of an intensity distribution can be estimated with subpixel precision even when the object boundary is affected by imaging artifacts.

To illustrate this principle, we simulated fluorescent objects with identical center positions but different PSF widths and quantified the resulting localization error of the estimated centroid. As shown in Supplementary Figure 3, increasing PSF size substantially broadened the apparent object boundary, whereas the estimated centroid position remained largely unchanged across a broad range of PSF sizes. These results support the use of centroid-based measurements as a robust complementary metric for spatial proximity analysis.

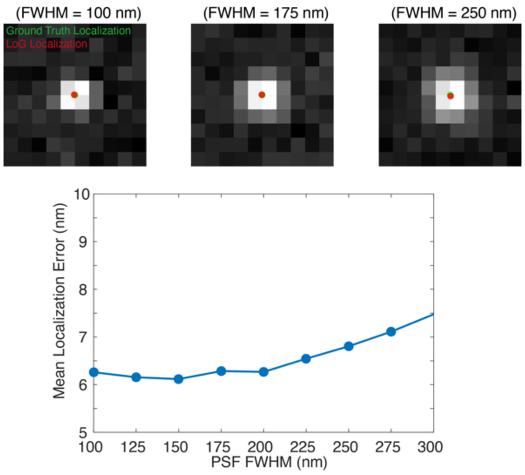


**Supplementary Figure 3. Effect of PSF size on centroid localization accuracy.** Simulated fluorescent objects with identical ground-truth center positions but varying point spread function (PSF) full width at half maximum (FWHM) values were generated and analyzed using the same centroid localization procedure applied to experimental data. Representative examples spanning the tested PSF range are shown.

#### Localization accuracy of assembly centroids

To quantify the accuracy of assembly centroid localization under our experimental imaging conditions, we generated simulated images containing assemblies with known ground-truth positions. Simulation parameters were selected to match experimentally measured signal intensities, background noise levels, assembly densities, and size distributions.

Because OCT4 and BRD4 assemblies exhibit relatively dense nuclear distributions, we simulated 300 assemblies with radii ranging from 100 to 200 nm within a 12.16 μm × 12.16 μm field of view. Assembly sizes were chosen based on measurements obtained using Sparse-SIM imaging. Background intensity and Gaussian noise were modeled using experimentally measured values (background mean intensity = 3500; standard deviation = 500; SNR = 24 dB). To account for freely diffusing nuclear molecules, an additional non-uniform background component was introduced using a Gaussian-filtered intensity field (mean intensity = 7000; standard deviation = 1000).

Simulated assembly images were processed using the same denoising and deconvolution pipeline applied to experimental assembly data, using two-dimensional Laplacian-of-Gaussian (LoG) filtering followed by quadratic fitting for subpixel localization. Comparison between predicted and ground-truth positions yielded a mean centroid localization error of approximately 34 nm (Supplementary Figure 4A–E).

We further estimated localization uncertainties directly from experimental data according to the following relationship[4]:

$$\sigma_{loc}^2 = \frac{2\sigma_{noise}}{|f''(r)|} ,$$

where $f(r)$ represents the fitted quadratic function, $r$ denotes the estimated centroid position, and $\sigma_{noise}$ is the standard deviation of background noise. Application of this method to experimental data yielded a mean localization error of 38 nm, in close agreement with the simulation-based estimate (Supplementary Figure 4F, G).

Together, these results indicate that assembly centroids can be localized with a mean error or estimated uncertainty of approximately 34-38 nm under our imaging conditions. Therefore, for analyses focused on spatial proximity, focus-to-centroid distance measurements provide a robust and reproducible metric.

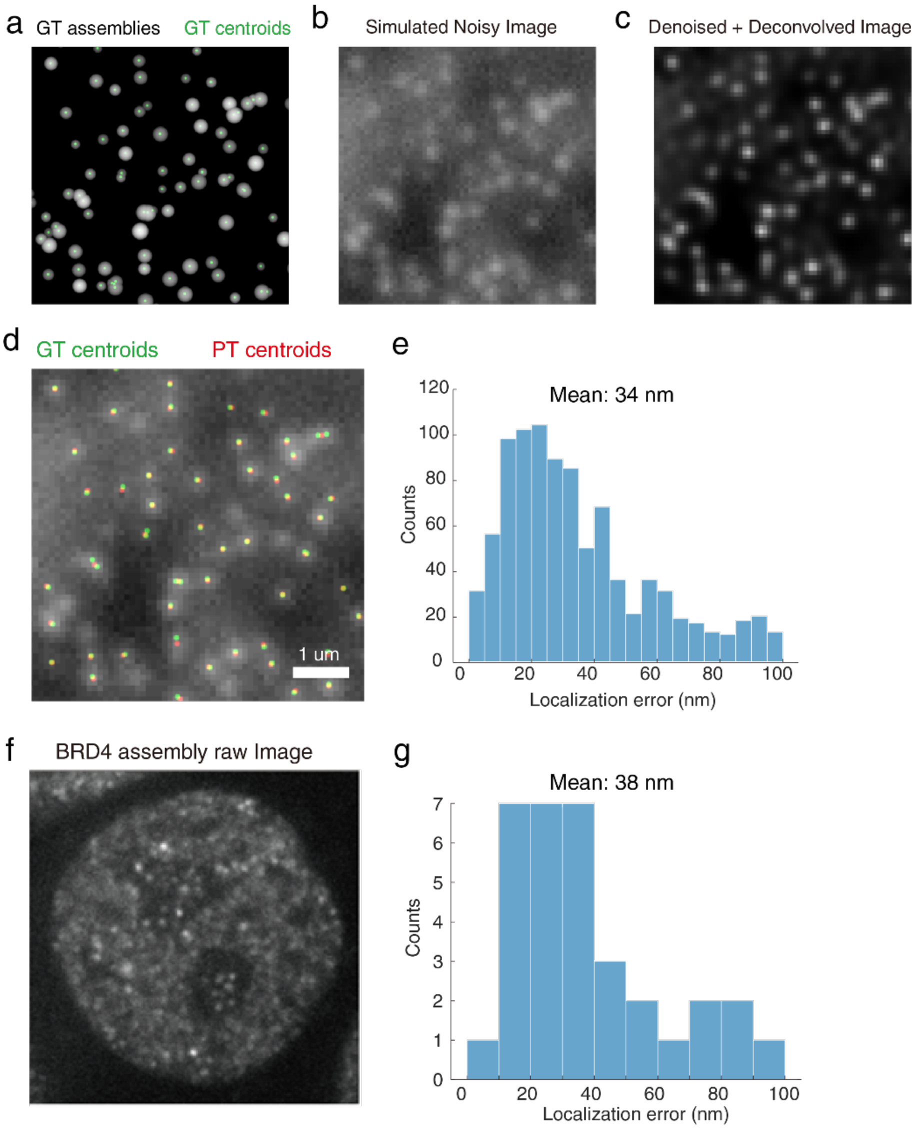


**Supplementary Figure 4. Estimating assembly centroid localization error using simulated and experimental data. (A)** Ground-truth (GT) distribution of assemblies; corresponding centroid positions are indicated in green. **(B)** Simulated noisy image generated under experimental imaging conditions. **(C)** Image processed using the same denoising and deconvolution pipeline applied to experimental data. **(D)** Overlay of GT and predicted (PT) centroid positions on the simulated noisy image. **(E)** Distribution of localization errors between PT and GT centroids. **(F)** Representative raw experimental image of BRD4 assemblies. **(G)** Distribution of localization errors estimated from experimental data using the described equation.

### Localization accuracy of DNA/RNA foci

To estimate localization accuracy for DNA and RNA foci, we simulated spatially separated puncta with radii ranging from 25 nm to 50 nm within a 12.16 μm × 12.16 μm field of view, with known ground-truth coordinates. Background intensity and Gaussian noise were modeled using experimentally measured values (mean intensity = 3500; standard deviation = 500). An SNR of 24 dB, corresponding to approximately threefold signal enrichment above background, was used to match experimental observations.

Simulated foci images were processed using the same denoising procedure applied to experimental foci data, followed by a 3D LoG detector and 3D quadratic fitting to achieve subpixel localization precision[5]. Comparison between predicted and ground-truth positions yielded mean localization errors of approximately 15 nm in 2D (xy) and 37 nm in 3D (xyz) (Supplementary Figure 5 A to F).

To assess localization uncertainty in experimental data, we estimated localization uncertainties directly from experimental data according to the following relationship, $\sigma_{loc}^2 = \frac{2\sigma_{noise}}{|f''(r)|}$ , as above. The resulting estimated localization errors of labeled promoter, RNA, and enhancer foci were consistently slightly larger than those obtained from simulations (Supplementary Figure 5G–I), likely reflecting additional sources of variability not captured by the simulations, including heterogeneous focus morphology, deviations from ideal Gaussian intensity distributions, residual motion blur, and local background fluctuations.

The simulation-derived localization errors are comparable to values previously reported from fluorescent bead calibration experiments[6], whereas the experimentally derived values are consistent with localization uncertainties reported for chromatin loci in living cells[7]. We therefore regard the simulation-derived values as lower-bound estimates under the modeled imaging conditions and use the experimentally derived localization errors as conservative estimates for subsequent analyses.

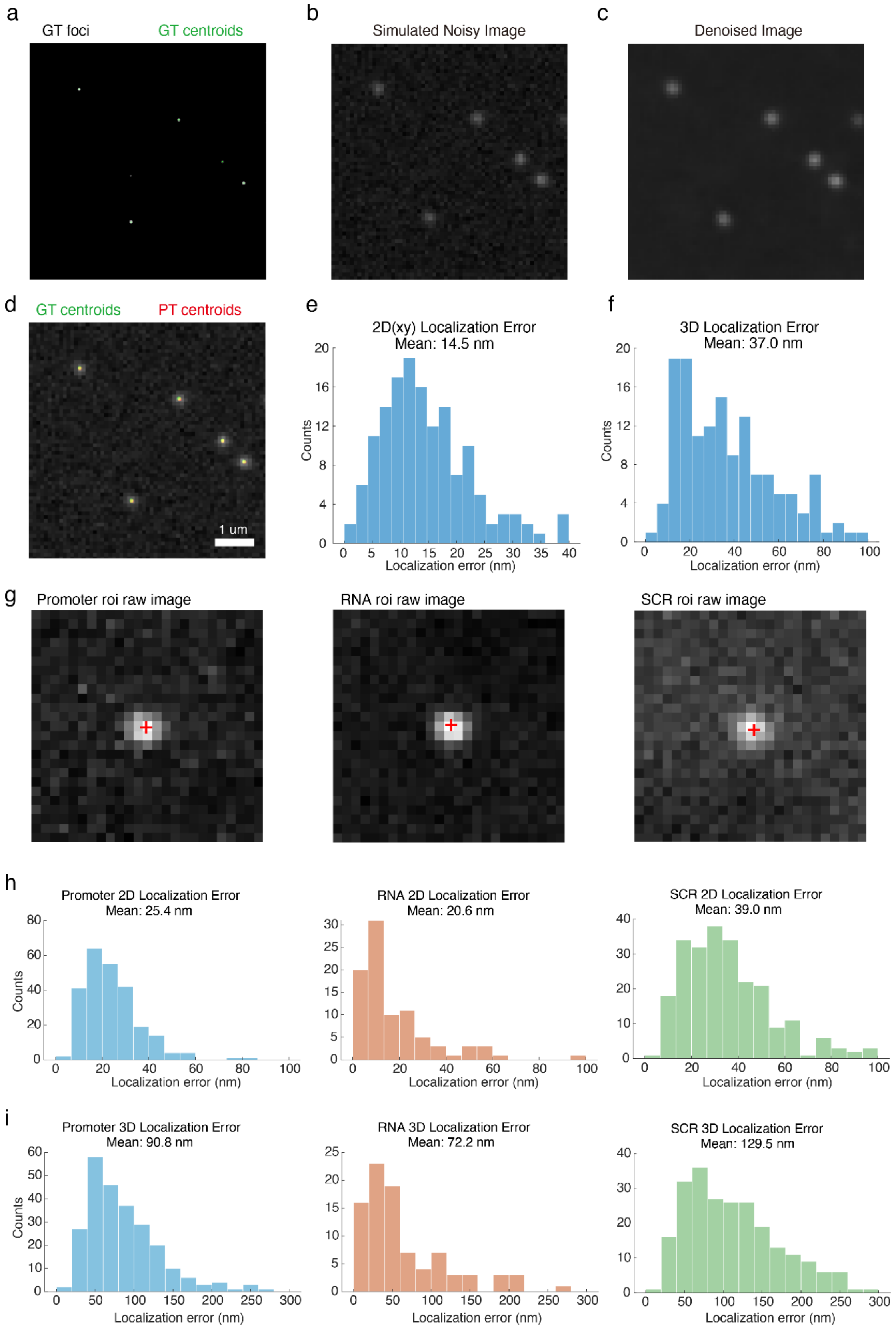


**Supplementary Figure 5. Estimating DNA/RNA foci localization error using simulated and experimental data.** **(A)** Ground-truth (GT) distribution of foci; corresponding positions are indicated in green. **(B)** Simulated noisy image generated under experimental imaging conditions. **(C)** Image processed using the same denoising pipeline applied to experimental data. **(D)** Overlay of GT and predicted (PT) centroid positions on the simulated image. **(E-F)** Distributions of 2D (xy, **E**) and 3D (xyz, **F**) localization errors between PT and GT positions. **(G)** Representative images of indicated DNA/RNA foci; red cross indicates the fitted position. **(H-I)** Distributions of localization errors estimated from experimental data in 2D (xy, **H**) and 3D (xyz, **I**).

**Supplementary Note 3: Error analysis of direct 3D distance measurements and 2D projection-based distance estimates**

Although the spatial relationships between genomic loci are inherently three-dimensional, localization precision in confocal microscopy is substantially poorer along the axial (z) dimension than along the lateral (xy) dimensions. Consequently, direct 3D distance measurements are subject to considerable uncertainty arising from axial localization error. In contrast, distance estimates derived from 2D projections eliminate axial localization error but introduce geometric projection error because the true 3D separation is projected onto a two-dimensional plane.

To quantitatively evaluate this trade-off, we compared the expected measurement uncertainties associated with direct 3D distance measurements and 3D distance estimates derived from 2D projections. Under our imaging conditions, 2D projection-based estimates showed lower expected error for true spatial separations below approximately 200-300 nm, whereas direct 3D measurements showed lower expected error at larger distances. The theoretical framework underlying this analysis is described below.

**Evaluating the error in direct 3D distance measurements**

Assume two foci separated by a true 3D distance:

$$r = \sqrt{x^2 + y^2 + z^2},$$

where $x, y,$ and $z$ denote the spatial separations along each axis.

The variance associated with direct 3D distance measurements is:

$$\sigma^2_{Direct\ 3D} = Var(r) = Var\left(\sqrt{x^2 + y^2 + z^2}\right)$$

When $r \gg Var(x), Var(y), Var(z)$, the variance can be approximated using first-order uncertainty propagation as

$$\sigma^2_{Direct\ 3D} = \left(\frac{x}{r}\right)^2 Var(x) + \left(\frac{y}{r}\right)^2 Var(y) + \left(\frac{z}{r}\right)^2 Var(z)$$

Assuming isotropic orientation of spatial separations,

$$E\left(\left(\frac{r_i}{r}\right)^2\right) = \frac{1}{3},$$

for each axis $i$.

Because both localized foci contribute localization uncertainty,

$$Var(r_i) = 2\sigma^2_{loc,i},$$

where $\sigma_{loc,i}$ denotes the localization uncertainties along axis $i$.

For locus pairs with different localization uncertainties, $\sigma^2_{loc,i}$denotes the mean of the two per-focus localization-error variances along axis i, such that $2\sigma^2_{loc,i} = \sigma^2_{1,i} + \sigma^2_{2,i}$

Substituting these values yields

$$\sigma^2_{Direct\ 3D} = \frac{4}{3}\sigma^2_{loc,xy} + \frac{2}{3}\sigma^2_{loc,z}.$$

**Evaluating the error in 2D projection-based distance estimates**

The total variance associated with the projected-distance estimation contains localization error and geometric projection error,

$$\sigma^2_{Projected\ 2D} = \sigma^2_{localization} + \sigma^2_{projection}(r).$$

For 2D projection-based analysis, the projected distance is:

$$r_{2D} = \sqrt{x^2 + y^2}$$

When $r_{2D} \gg Var(x), Var(y),$ the variance can be approximated using first-order uncertainty propagation as

$$\sigma^2_{localization} = Var\left(\sqrt{x^2 + y^2}\right) = \left(\frac{x}{r_{2D}}\right)^2 Var(x) + \left(\frac{y}{r_{2D}}\right)^2 Var(y)$$

Assuming isotropic orientation within the xy plane:

$$\sigma^2_{localization} = \frac{1}{2}Var(x) + \frac{1}{2}Var(y) = \sigma^2_{loc,xy}(x,y)$$

The second component arises because projection removes information about the z coordinate. For a true 3D distance $r$, the probability distribution of $r_{2D}$ is:

$$P(r_{2D}) = \frac{r_{2D}}{r\sqrt{r^2 - r^2_{2D}}}$$

From this distribution,

$$\sigma^2_{projection}(r) = Var(r_{2D}) = \left(\frac{2}{3} - \frac{\pi^2}{16}\right) r^2$$

Thus, the total variance of the 2D projection-based estimate is

$$\sigma^2_{Projected\ 2D} = \sigma^2_{loc,xy}(x,y) + \left(\frac{2}{3} - \frac{\pi^2}{16}\right) r^2$$

To compare the error associated with 2D projection-based measurements directly with $\sigma_{Direct\ 3D}$, the systematic shortening introduced by projection must first be corrected. Based on the distribution $P(r_{2D})$, the expected mean projected distance is:

$$E(r_{2D}) = \frac{\pi}{4} r$$

Thus, the projected distance was rescaled by a factor of $\frac{4}{\pi}$ to obtain an unbiased estimator of the true 3D distance. Correspondingly, the variance of the corrected 2D measurement becomes

$$\frac{4}{\pi} \sigma_{Projected\ 2D}$$

Because the projected distance is multiplied by $\frac{4}{\pi}$, its standard deviation is multiplied by the same factor (and its variance by $\left(\frac{4}{\pi}\right)^2$). This rescaled uncertainty was used for the analytical comparison between 2D projection-based and direct 3D measurement strategies.

The rescaling was used for error comparison and model-based inference of 3D distance; the experimental 2D distance distributions presented in the main figures remain unscaled projected distances.

### Comparison between 3D and 2D measurement strategies

Using the analytical framework described above, we next compared the expected errors associated with direct 3D measurements and 2D projection-based measurements.

We first evaluated the theoretical standard deviation of each strategy as a function of the true 3D distance (Supplementary Fig. 6A). We further converted the calculated mean localization errors into localization uncertainties. Assuming independent, zero-mean Gaussian localization errors with equal variance along the lateral axes, the mean radial localization error was converted into the standard deviation of each lateral coordinate using $\sigma_{loc,xy} = \sqrt{\frac{2}{\pi}} E(r_{xy})$, where $E(r_{xy})$ represents the measured mean radical 2D localization error in Supplementary Note 2. Similarly, the axial localization uncertainty was calculated as $\sigma_{loc,z} = \sqrt{\frac{\pi}{2}} E(r_z)$, where $E(r_z)$ represents the mean absolute axial localization error.

Incorporating the experimentally determined localization errors of each labeled element—for example, promoter mean labeling errors of 25 nm in 2D and 91 nm in 3D, and RNA mean labeling errors of 21 nm in 2D and 72 nm in 3D—revealed a distance-dependent trade-off in measurement precision. At short distances, the uncertainty introduced by projection into two dimensions is smaller than the localization uncertainty associated with direct 3D measurements. For promoter–RNA measurements, the predicted crossover occurs at a true distance of approximately 210 nm (Supplementary Fig. 6A). Beyond this distance, direct 3D measurements become progressively more precise. Because the crossover distance depends on the overall localization uncertainty, it shifts to approximately 270 nm for the SCR–RNA pair and 280 nm for the promoter–SCR pair, reflecting the larger mean localization errors associated with SCR labeling (39 nm in 2D and 130 nm in 3D).

Because the analytical derivation relies on the simplified assumption of first-order error propagation, which becomes less accurate when the true separation is comparable to or smaller than the localization error and does not fully capture the positive bias of non-negative distance measurements at very short separations, we further evaluated this trade-off using Monte Carlo simulations. For pairs of loci separated by a defined true distance, localization uncertainties for each locus were introduced according to the experimentally determined localization errors, and the resulting distances were repeatedly sampled to calculate the root-mean-square error (RMSE) of each measurement strategy (Supplementary Fig. 6B). This yielded crossover ranges similar to those predicted by the analytical model. At larger distances, the simulation results agreed closely with the theoretical predictions. At short distances, however, the errors associated with direct 3D measurements increased substantially beyond those predicted by the analytical model and became progressively larger as the true distance approached zero. In contrast, 2D projection-based measurements exhibited only a modest increase in error. Nevertheless, direct 3D measurements remained considerably more error-prone than 2D projection-based measurements throughout this regime.

Notably, the large measurement errors associated with direct 3D measurements at short distances also introduce a substantial positive bias in the measured distances. Consequently, the mean measured 3D distance increasingly overestimates the true spatial separation as the underlying distance approaches zero (Supplementary Fig. 6C). This effect partially explains why the mean value of experimentally measured promoter–RNA 3D distances is approximately 200 nm, despite the expectation that promoter and nascent RNA signals should be nearly colocalized. The observed discrepancy is further influenced by the genomic offset between the promoter and the labeled RNA sequence, as discussed in detail in Supplementary Note 4.

In summary, these results indicate that below the predicted crossover distances, 2D projection-based measurements have lower expected measurement error than direct 3D measurements under our imaging conditions.

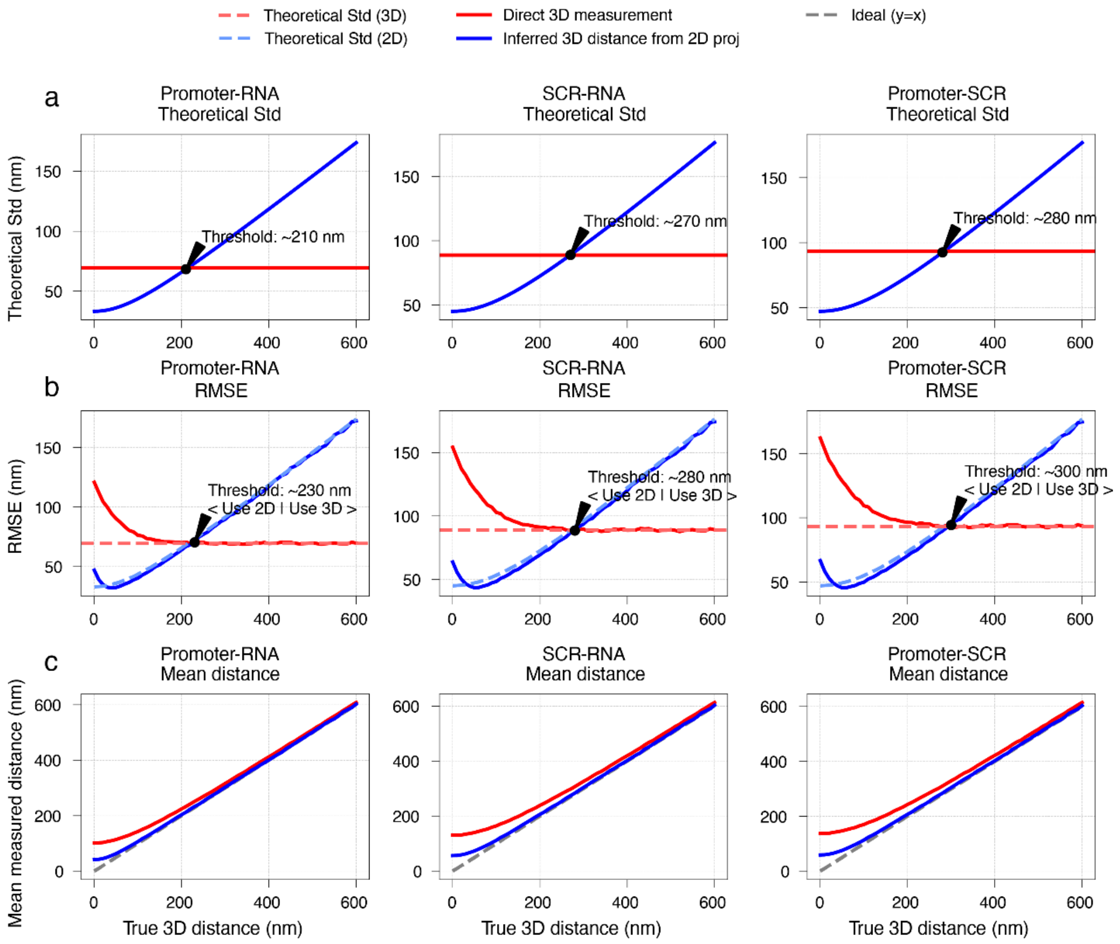


**Supplementary Figure 6.** Comparison of measurement errors between direct 3D measurements and 3D distances inferred from 2D projections. **(A)** Theoretical standard deviation versus true 3D distance for direct 3D measurements and 3D distances inferred from 2D projections. The crossover point is marked with a black dot. **(B)** RMSE versus true 3D distance for direct 3D measurements and 3D distances inferred from 2D projections. The crossover point is marked with a black dot. **(C)** Mean measured distance versus true 3D distance for direct 3D measurements and 3D distances inferred from 2D projections. In each panel, the corresponding experimentally determined localization errors for the indicated DNA/RNA element pair were incorporated into the calculations or simulations. Distances inferred from 2D projections were obtained by multiplying the projected distance by a scaling factor of $\frac{4}{\pi}$.

**Supplementary Note 4: Quantification of distance measurement bias arising from genomic separation between labeled sites and target sites**

To minimize perturbation of endogenous regulatory function, the DNA-labeling arrays were inserted a few kilobases (kb) from, rather than directly within, the regulatory elements of interest. In our system, the promoter-proximal array is located approximately 8 kb upstream of the Sox2 transcription start site (TSS), the SCR-proximal array approximately 4 kb downstream of the SCR, and the MS2 cassette in the Sox2 3′ UTR approximately 2 kb downstream of the TSS. These labeling positions were selected to reduce disruption of core regulatory sequences, but they introduce genomic offsets between the labeling sites and their corresponding functional elements. Consequently, measured distances between fluorescent signals do not directly report the underlying spatial distances between the corresponding functional elements (Supplementary Figure 9).

To quantify the impact of these labeling offsets, we used chromatin polymer simulations to estimate how genomic separation between labeling sites and target sites affects the observed distances. We then used the resulting relationship to infer the underlying functional-element distance distributions from the measured label-to-label distances under the stated polymer assumptions.

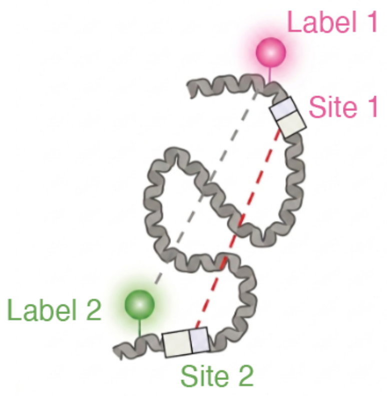


**Supplementary Figure 9. Relationship between target sites and fluorescent labeling sites**.

**Chromatin polymer simulations**

Chromatin dynamics were simulated using the Polychrom framework (OpenMM-based) following a previously established live-cell chromatin modeling strategy[8]. Chromatin was represented as a 1000-kb polymer chain discretized into 1-kb monomers, corresponding to a contour length of 25 nm per monomer. Adjacent monomers were connected by harmonic bonds, and excluded-volume interactions were implemented using parameters identical to those described previously. Experimentally determined localization uncertainties (Supplementary Note 2) were incorporated into simulated coordinates prior to distance measurements. Unless otherwise indicated, distance distributions were generated from 100,000 simulation iterations.

**Validation of the simulation framework using promoter-RNA distances**

We first evaluated whether the chromatin simulation framework could account for distance bias arising from labeling offsets using promoter-RNA distance measurements. Because the nascent RNA focus reports the transcription site of the short Sox2 gene, its underlying spatial separation from the functional promoter is expected to be small relative to the measured label-to-label distance.

However, the experimentally measured promoter-RNA distances are influenced by both localization errors and the genomic offset between their labeling positions. Specifically, the promoter label is positioned approximately 8 kb upstream of the TSS, whereas the MS2 cassette generating the nascent RNA signal is located in the 3′ UTR, approximately 2 kb downstream of the TSS. Consequently, the promoter and RNA labeling positions are separated by approximately 10 kb along the chromatin fiber, introducing a systematic spatial offset even when the promoter and RNA themselves are effectively apposed.

To determine whether this labeling geometry can account for the experimentally observed promoter-RNA distances, we simulated a 10-kb genomic separation using the chromatin polymer model and incorporated experimentally determined localization uncertainties for promoter and RNA labeling (Supplementary Note 2). The simulated promoter-RNA distance distributions closely matched the experimentally measured distributions in both 2D and 3D (Supplementary Figure 10).

This agreement supports the use of the simulation framework to estimate the principal distance bias arising from labeling offsets and localization uncertainty across a range of underlying distances.

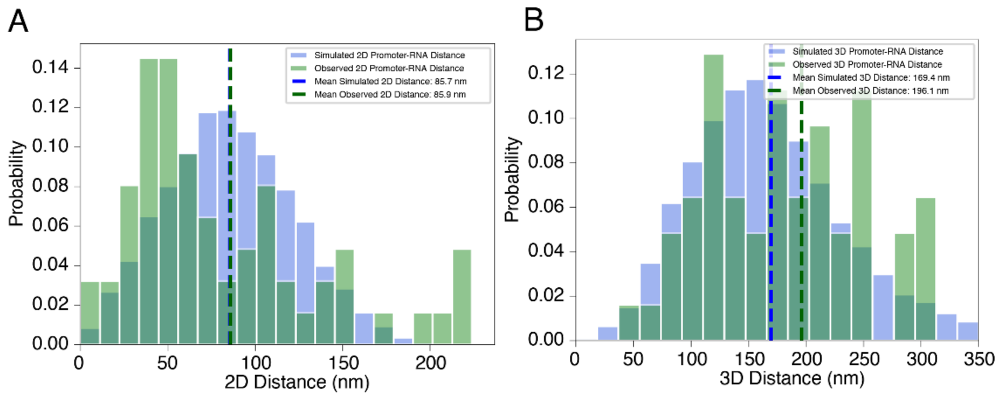


**Supplementary Figure 10. Comparison of simulation-generated and experimentally observed promoter-RNA distance distributions.** Simulated and experimentally measured promoter-RNA distance distributions in 2D (**A**) and 3D (**B**).

**Mapping the relationship between measured and underlying distances**

Having established that the simulation framework reproduced the promoter–RNA distance distributions under the modeled conditions, we next quantified how labeling-site offsets and localization uncertainty influence measured distances across a broad range of underlying functional-element separations, hereafter referred to as true distances within the simulation.

To map this relationship, we arranged key sites on the simulated chromatin according to the Sox2 locus, incorporating both the genomic separation between the SCR enhancer and Sox2 promoter and the experimentally used labeling offsets (8 kb at the promoter and 4 kb at the SCR) (Supplementary Figure 9). We then generated 100,000 chromatin conformations spanning a range of underlying SCR-promoter distances across the experimentally relevant range (approximately 0-350 nm). Analysis of these conformations yielded the relationship between measured label-to-label distances and underlying functional-element distances (Supplementary Figure 11).

For each simulated chromatin conformation, we recorded both the true distance between the target sites and the corresponding distance between their label sites, either without (Figure 11A) or with (Figure 11B, C) incorporation of experimentally determined localization uncertainties (Supplementary Note 2). We further examined the relationship between true and measured distances using both 2D and 3D measurement strategies (Supplementary Note 3). As shown in Supplementary Figure 11, genomic labeling offsets and localization uncertainty increased the mean measured distances relative to the corresponding true distances, with the deviation becoming more pronounced at shorter true separations. In addition, over the short-distance range examined here, distances inferred from 2D measurements more closely tracked the underlying true 3D distances than direct 3D measurements, consistent with the findings described in Supplementary Note 3.

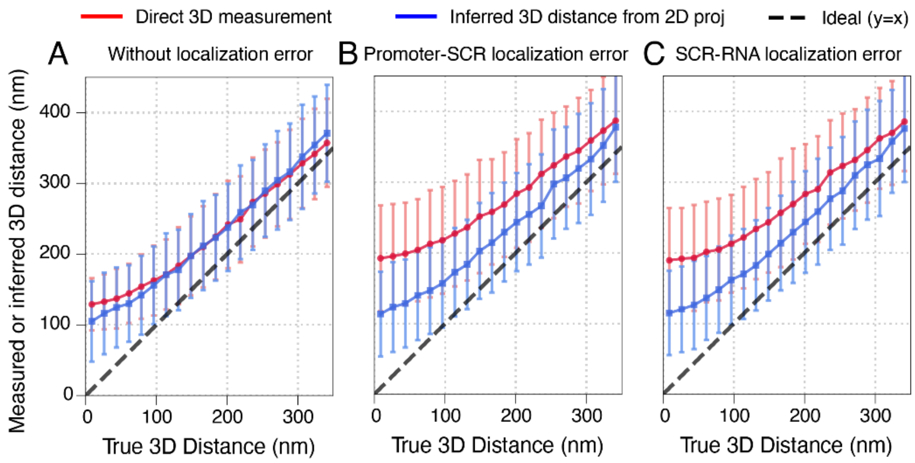


**Supplementary Figure 11. Relationship between measured and true distances derived from chromatin simulations.** Simulated measured distance is plotted as a function of the underlying true distance in the absence of localization error **(A)**, after incorporation of promoter and SCR localization uncertainties **(B)**, or after incorporation of SCR and RNA localization uncertainties **(C)**. Genomic labeling offsets and localization uncertainty cause measured distances to systematically exceed true distances, particularly at short spatial separations.

**Inferring underlying SCR-promoter distances from measured separations**

Using this simulated relationship, we next inferred the underlying spatial-distance distributions for experimentally observed DNA/RNA element pairs under the stated polymer assumptions.

For SCR-promoter distances, this mapping revealed broad underlying 3D distance distributions centered at approximately 150 nm in both bursting and non-bursting alleles, regardless of whether inference was based on 2D or 3D measurements (Supplementary Figure 12). This result was robust to the choice of measurement strategy and consistent across the full range of observed experimental distances. Importantly, the inferred SCR-promoter distances were substantially larger than those obtained for promoter-RNA pairs, indicating that the SCR and promoter were not uniformly in the closely apposed configuration as the promoter and nascent RNA. Very short SCR-promoter distances, potentially compatible with direct contact, were nevertheless observed in both states, indicating that such close configurations can occur do not uniquely specify the bursting state.

Because the functional promoter and nascent transcription site are expected to be closely apposed, much of the apparent promoter-RNA label separation reflects labeling-site offsets and localization uncertainty rather than underlying functional separation. Consistent with the findings in Supplementary Note 3, promoter-RNA distances inferred from 2D measurements were substantially smaller than those obtained from direct 3D measurements. This difference arises because, within the short-distance regime (<230 nm), projection-based 2D measurements have lower measurement error than direct 3D measurements, resulting in more reliable inference of the underlying 3D distance.

Notably, the inferred SCR-promoter separation (~150 nm) is comparable to the assembly diameters measured in this study. Together with the four-color imaging data, these results are consistent with a model in which transcriptional bursting is associated with a shared regulatory assembly environment rather than with a uniformly close SCR-promoter configuration.

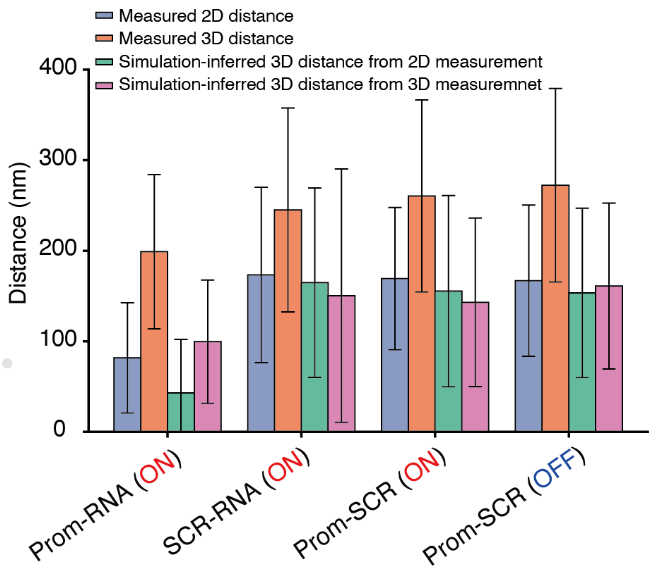


**Supplementary Figure 12. Comparison of measured distances and simulation-inferred underlying distances for DNA/RNA element pairs.**
